# RNA polymerase II links arsenic exposure to chromosome-wide γH2A signaling

**DOI:** 10.64898/2026.11.16.767017

**Authors:** Adrianna Nawrot, Paulina Tomaszewska, Ewa Maciaszczyk-Dziubińska, Karol Kramarz, Michał Tracz, Aleksandra Domaradzka, Dharshanbaabu Pennagaram Balaji, Tomasz W. Turowski, Artur P. Kaczmarczyk, Ireneusz Litwin

## Abstract

Arsenic is a widespread environmental carcinogen whose genotoxicity is commonly attributed to oxidative DNA damage and replication-associated lesions. Here, we identify RNA polymerase II (RNAPII) as an important determinant of early arsenic-induced DNA damage checkpoint signaling. In budding yeast, acute As(III) exposure induced rapid, chromosome-wide γH2A accumulation under low oxidative stress, without requiring ongoing bulk DNA replication or causing detectable chromosome fragmentation. Unexpectedly, depletion of Rpb1, the largest subunit of RNAPII and a key component of its catalytic center, markedly attenuated this response. Comparison of different transcriptional perturbations revealed that only selected disruptions of RNAPII function, including Rpb1 inactivation in the *rpb1-1* mutant, induced widespread γH2A accumulation. These contrasting effects suggest that checkpoint activation depends on the presence and particular functional states of RNAPII. Consistent with its involvement, As(III) reduced RNAPII chromatin occupancy and Rpb1 abundance, while arsenic-affinity assays recovered the RNAPII complex. However, As(III) did not measurably inhibit RNA elongation by purified RNAPII *in vitro*. In human cells, acute As(III) exposure induced ATR phosphorylation and γH2AX accumulation under low-ROS conditions, while pharmacological inhibition of transcription also triggered γH2AX phosphorylation. Together, our findings establish an important contribution of RNAPII to early arsenic-induced checkpoint signaling and suggest that the transcription machinery can couple environmental stress to a chromosome-wide DNA damage response.

**Significance Statement:** Arsenic is a widespread environmental carcinogen, but how it activates cellular responses to genome stress remains unclear. Its genotoxicity has predominantly been explained by oxidative DNA damage and replication-associated DNA breaks. Here we show that arsenic triggers widespread DNA damage signaling in budding yeast despite low oxidative stress and without requiring ongoing DNA replication. Unexpectedly, depletion of a key subunit of RNA polymerase II, which transcribes protein-coding genes, weakens this response, whereas certain disruptions of its function can induce similar signaling. In human cells, both arsenic and transcription inhibitors induce DNA damage signaling. These findings suggest an underappreciated role for the transcription machinery in cellular responses to environmental genotoxic stress.

## INTRODUCTION

Genomes of all organisms are continuously challenged by DNA-damaging factors originating from endogenous metabolic processes and environmental sources, posing a threat to genomic stability. Eukaryotic cells preserve genome integrity through DNA damage checkpoint (DDC) signaling and DNA repair pathways. The sensing and repair of DNA double-strand breaks (DSBs) are among the most thoroughly characterized components of this response (1, 2). In the budding yeast *Saccharomyces cerevisiae*, the Mre11-Rad50-Xrs2 (MRX) complex recognizes DSBs and recruits Tel1 kinase to initiate DDC activation (3–5). The majority of DSBs arising in S/G2 undergo DNA end resection, initiated by MRX and extended by Exo1 or Dna2–Sgs1, generating 3′ single-stranded DNA (ssDNA) (5, 6). The resulting ssDNA is rapidly bound by the RPA complex (Rfa1–Rfa3) that promotes recruitment and activation of the major DDC kinase Mec1 (4, 7–9). Importantly, Mec1 can also be activated by RPA-coated ssDNA formed at stalled replication forks or DNA gaps (10–12). Beyond replication-associated structures, perturbations of transcription–DNA transactions can also expose ssDNA and trigger checkpoint signaling (11, 13, 14). For example, RNAPII stalling at UV-induced lesions promotes Exo1-dependent ssDNA formation and Mec1 activation. Under these conditions, the largest catalytic subunit of RNAPII, Rpb1, can also undergo degradation, linking transcriptional arrest to checkpoint activation and RNAPII turnover (11, 15).

Histone H2A is among the earliest targets of Tel1 and Mec1 and is phosphorylated at serine 129, a modification referred to as H2A-S129-P or γH2A. Much of our mechanistic understanding of γH2A formation and spreading comes from studies using site-specific, endonuclease-induced DSBs. In these systems, γH2A can be detected within minutes after DSB induction and spreads over chromosomal domains extending up to ∼50 kb on both sides of the lesion (16, 17). However, γH2A formation is not restricted to DSBs and can also occur during replication stress, when RPA-coated ssDNA activates Mec1 (10, 12).

While DDC and DNA repair pathways have been defined primarily in response to engineered DSBs and replication stress, far less is known about how environmental genotoxins engage this network. Arsenic (As) is a widespread environmental toxin that contaminates groundwater and crops worldwide. It is among the WHO’s 10 chemicals of major public-health concern, with ∼200 million people exposed above the provisional 10 μg/L limit (18–21). At the same time, As-containing compounds are used therapeutically, including arsenic trioxide for acute promyelocytic leukemia and melarsoprol for human African trypanosomiasis (22). Nevertheless, prolonged As exposure is associated with multisystem disorders and elevates the risk of skin, lung, bladder, kidney, and liver cancers (23). Despite extensive research, the mechanisms underlying As-induced genome instability remain unclear. A prevailing model invokes both DNA damage and inhibition of DNA repair. Arsenic-induced reactive oxygen species (ROS) can generate oxidative DNA lesions and single-strand breaks (SSBs), potentially promoting replication fork collapse (24–26). In parallel, As(III) can inhibit repair proteins including XPA and PARP1, compromising NER and PARP-dependent repair (27–30).

We have previously shown that in budding yeast, short-term exposure to As(III) triggers Mec1-dependent activation of the DDC (31). However, the proximal trigger that engages this response has remained undefined. Here, we show that As(III) induces a rapid and widespread γH2A response that is largely replication-independent and occurs without detectable chromosome fragmentation or a major contribution from oxidative stress. Unexpectedly, depletion of RNAPII markedly attenuates this response, whereas specific forms of RNAPII dysfunction are sufficient to trigger widespread γH2A accumulation. Together, these findings reveal an underappreciated contribution of RNAPII-associated processes to arsenic-induced DNA damage checkpoint signaling, extending current models of arsenic genotoxicity beyond oxidative damage and replication-associated lesions.

## RESULTS

### As(III) induces a rapid and widespread γH2A response

In *S. cerevisiae*, exposure to 0.5 mM As(III) for 1 h induced robust H2A S129 phosphorylation (Fig. 1A) (31). At this dose, viability was unaffected (SI Appendix, Fig. S1A), and intracellular As was estimated to reach ∼40 μM after 1–2 h of treatment (SI Appendix, Fig. S1B). This concentration is comparable to intracellular As levels reported in human cells exposed to 5–10 μM extracellular As(III) for 1–24 h (32–35). The response was specific to As(III), as Fe, Cu, Co, Ni, and Cd did not increase H2A phosphorylation under similar conditions (SI Appendix, Fig. S1C). γH2A induction was dose-dependent (SI Appendix, Fig. S1D), detectable within 15 min (SI Appendix, Fig. S1E), and persisted for up to 2 h before declining markedly by 4 h (SI Appendix, Fig. S1F). Functionally, H2A-S129A cells showed improved growth in the presence of As(III), but not after treatment with the DSB-inducing radiomimetic antibiotic phleomycin (Phl) (SI Appendix, Fig. S1G), consistent with previous observations (36). Thus, H2A-S129 phosphorylation contributes modestly to As(III)-induced growth inhibition.

**Fig. 1.**
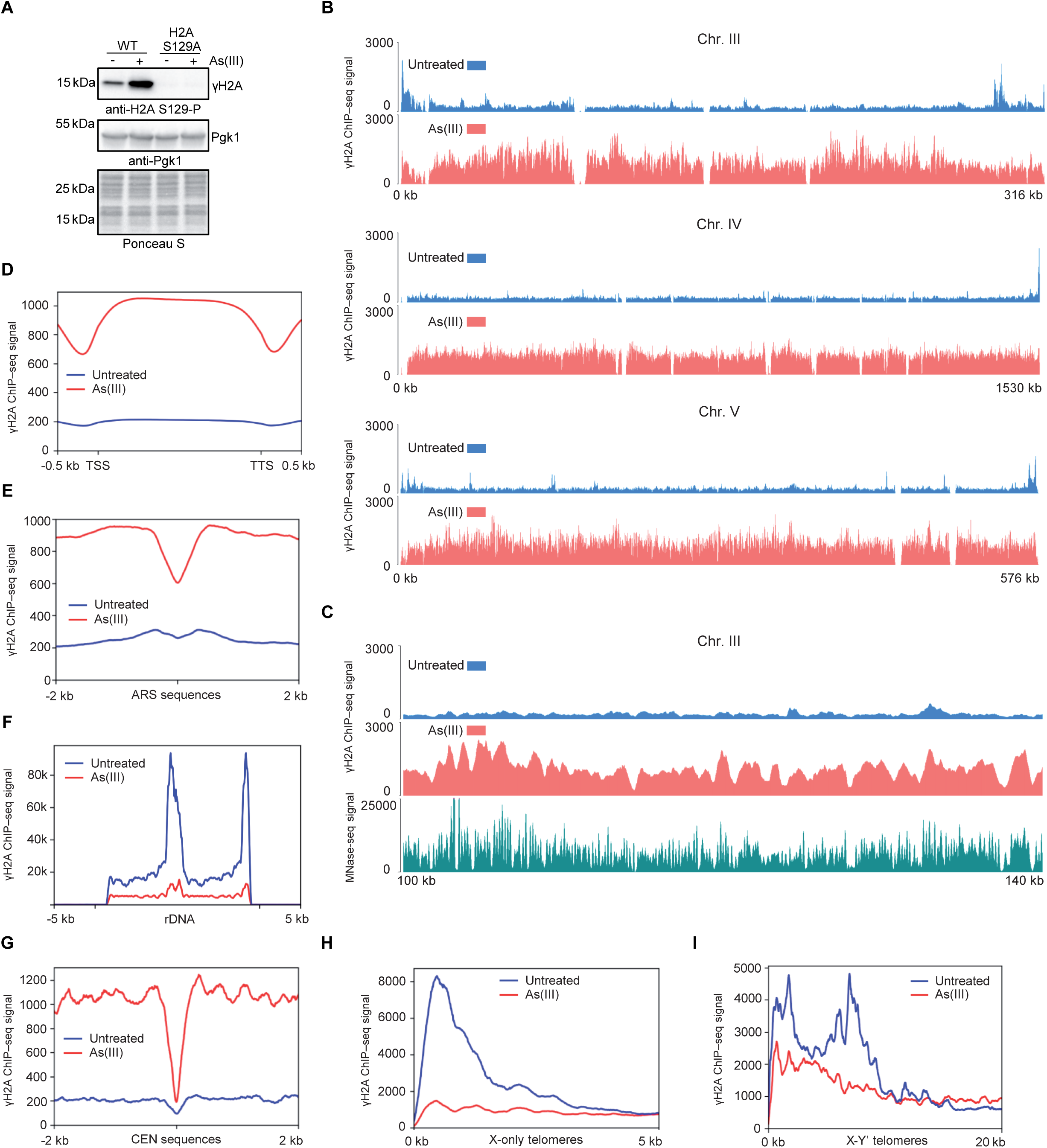
Broad As(III)-induced γH2A accumulation across the yeast genome. (A) Immunoblot analysis of γH2A in WT and H2A-S129A cells left untreated or exposed to 0.5 mM As(III) for 1 h. Pgk1 and Ponceau S staining served as loading and total-protein controls, respectively. (B–I) Spike-in-normalized γH2A ChIP–seq analyses of logarithmically growing WT cells left untreated or exposed to 0.5 mM As(III) for 1 h. γH2A profiles and metaprofiles are shown for representative biological replicate 1. Chromosome III profiles from all three independent biological replicates and their genome-wide correlations are presented in SI Appendix, Fig. S2 A and B, respectively. (B) Profiles along chromosomes III, IV, and V. (C) Expanded view of a representative chromosomal region showing γH2A profiles alongside MNase-seq signal from untreated WT cells. (D-I) γH2A metaprofiles across scaled TSS–TTS gene bodies (D), ARSs (E), rDNA (F), centromeres (G), and X-only (H) and X–Y′ telomere-proximal regions (I). The two rDNA repeat units annotated in the S288C reference genome are shown. Because multimapping reads were retained (MAPQ 0), the signal includes contributions from multiple copies of the endogenous rDNA array, which contains approximately 150 tandem repeats.

To characterize the genomic distribution of As(III)-induced γH2A, we performed spike-in-calibrated γH2A ChIP–seq in logarithmically growing WT cells. Under control conditions, γH2A levels were generally low across chromosomal regions, with several discrete enriched sites. After 1 h of As(III) exposure, γH2A ChIP–seq signal increased broadly across chromosomes, corresponding to an approximately 4.6-fold genome-wide increase (Fig. 1B). The increase was reproducible across three independent biological replicates which showed strong genome-wide correlations (SI Appendix, Fig. S2 A and B). ChIP–seq in H2A-S129A cells yielded only background-level signal, confirming γH2A specificity (SI Appendix, Fig. S2C). Closer inspection of chromosomal profiles revealed local peaks and troughs superimposed on this broad increase (Fig. 1C). Because γH2A is a nucleosomal histone modification, we asked whether this variation reflects underlying nucleosome organization. Visual inspection revealed partial correspondence between γH2A and MNase-seq profiles (Fig. 1C), and genome-wide γH2A signal after As(III) correlated with nucleosome occupancy (Spearman’s ρ = 0.38). Nevertheless, As(III)-dependent induction extended across the full occupancy range, including a 3.8-fold increase in the lowest MNase-occupancy decile. Thus, nucleosome organization contributes to local γH2A variation but does not explain the broad As(III)-induced increase.

Next, we examined γH2A enrichment across transcribed regions, replication origins, rDNA locus, centromeres, and telomeres. Across genes, γH2A was distributed throughout gene bodies but reduced around promoters and terminators (Fig. 1D). At replication origins, As(III) increased γH2A in surrounding chromatin, whereas origin-centered regions showed locally reduced signal (Fig. 1E). These local reductions are consistent with the lower nucleosome occupancy characteristic of promoter, terminator and ARS-associated regions (37, 38). At rDNA, a site of constitutive γH2A enrichment linked to replication–transcription conflicts (39), γH2A decreased markedly after As(III) (Fig. 1F). This observation is consistent with previous reports of arsenite-induced suppression of rRNA synthesis and ribosome biogenesis (40, 41). Centromeres showed a distinct pattern, with low γH2A at CEN-centered regions under both conditions but increased signal in surrounding pericentromeric chromatin after As(III) treatment (Fig. 1G). γH2A (γH2AX in human cells) is constitutively enriched in telomere-adjacent chromatin in budding yeast and human cells (39, 42, 43). Surprisingly, γH2A decreased after As(III) in telomere-proximal chromatin at both X-only and X–Y′ telomeres (Fig. 1 H and I). Importantly, ChIP–qPCR measurements at selected loci were consistent with the corresponding ChIP–seq changes (SI Appendix, Fig. S2D). Together, these findings reveal distinct patterns of γH2A responses to As(III) across different genomic regions.

Previous work showed that As(III) induces Rfa1-YFP foci (31). Therefore, we profiled RPA genome-wide and generated ORF metaprofiles. These analyses did not reveal prominent new RPA-enriched domains; instead, mean RPA signal decreased modestly (SI Appendix, Fig. S3 A and B). This decrease may partly reflect the increased fraction of cells in G1 (SI Appendix, Fig. S3C), where DNA end resection and RPA loading onto ssDNA are largely suppressed (44). Alternatively, the decrease in RPA occupancy may reflect its displacement by HR factors at a subset of DNA lesions.

Building on previous evidence that As(III) induces Rad52-YFP foci and that HR promotes survival during As(III) stress (31), we profiled Rad52, Rad51, and Rad59 occupancy with or without As(III). The distributions of all three HR factors were similar between conditions, and metaprofiles over ORFs showed modest, broadly distributed As(III)-dependent increases rather than the discrete enrichment domains (SI Appendix, Fig. S4 A-F).

Taken together, these results indicate that As(III) induces widespread γH2A accumulation across chromosomes, with distinct local patterns at rDNA, centromeres, and subtelomeric regions. In contrast, genome-wide profiling of RPA and HR factors revealed only modest changes in chromatin occupancy, without prominent new enrichment domains.

### As(III) elicits checkpoint response distinct from canonical DSB signaling

To determine whether the As(III)-induced response resembles canonical DSB-associated damage, we used Phl as a reference and matched Phl exposure separately to two As(III)-induced readouts: bulk γH2A induction and Rad52–YFP focus formation, a marker of homologous recombination-associated DNA repair. Matching As(III)-induced γH2A required 50 µg/ml Phl (Fig. 2 A and B). At this dose, γH2A remained elevated for at least 4 h (SI Appendix, Fig. S5A), while viability was markedly reduced and Rad52–YFP foci were more than twice as frequent as after As(III) (Fig. 2 C and D). Conversely, 2.5 µg/ml Phl produced Rad52–YFP foci at levels similar to As(III) (Fig. 2D), but only a weak γH2A response (Fig. 2E) with no progressive increase over 4 h (SI Appendix, Fig. S5B) and caused a modest viability defect (Fig. 2C). Genome-wide profiling likewise showed little γH2A accumulation after 2.5 µg/ml Phl but pronounced accumulation after 50 µg/ml Phl (SI Appendix, Fig. S5C). Taken together, the distinct response profiles elicited by As(III) and Phl argue against a simple Phl-like burden of canonical DSBs.

**Fig. 2.**
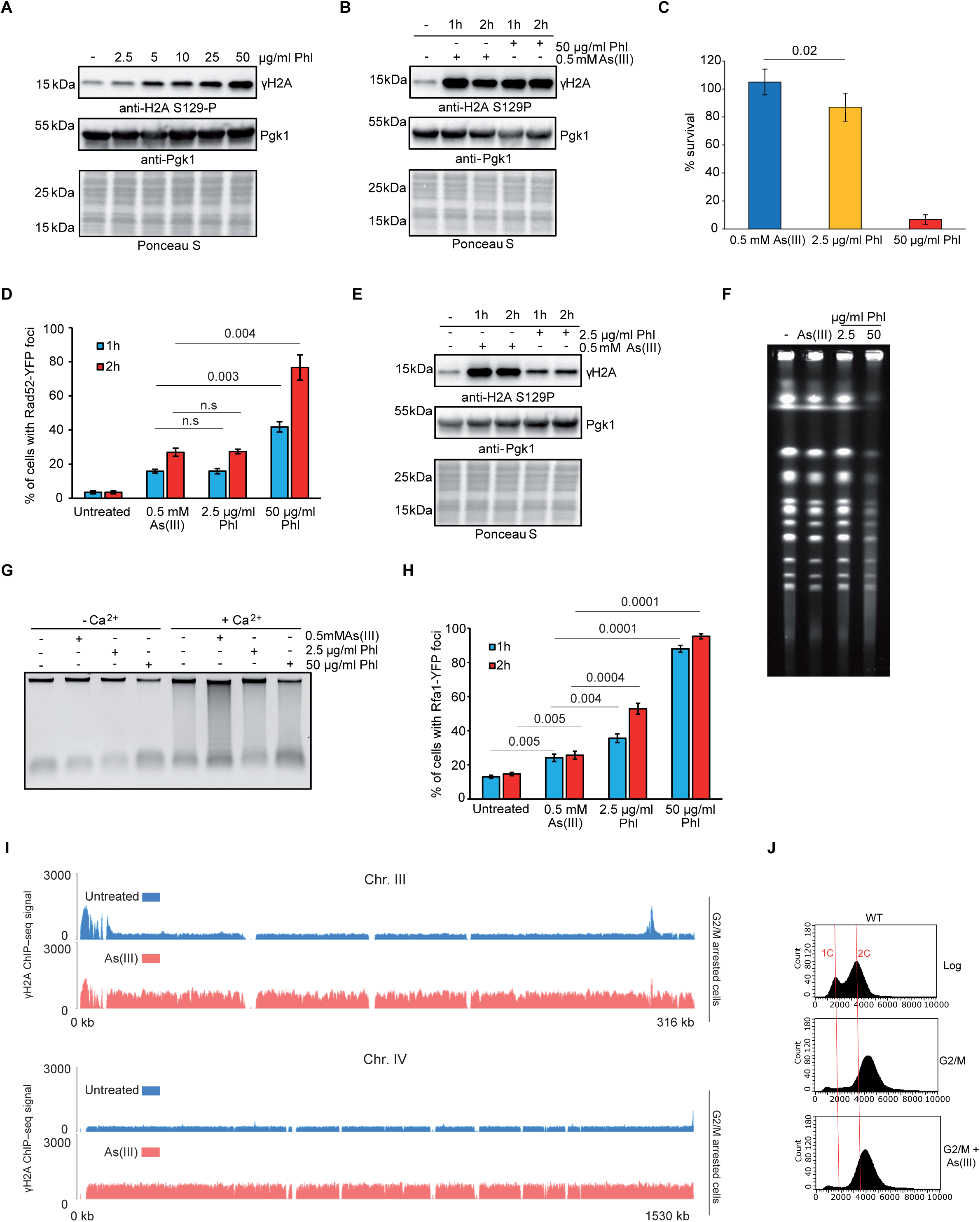
As(III) and phleomycin show distinct relationships between γH2A induction, Rad52 foci, and cell survival. Phl was used at 50 µg/ml as a γH2A-matched reference and at 2.5 µg/ml as a Rad52–YFP-focus-matched reference to 0.5 mM As(III). (A) γH2A immunoblot of logarithmically growing WT cells treated with the indicated concentrations of phleomycin (Phl) for 1 h. (B) γH2A immunoblot of logarithmically growing WT cells left untreated or exposed to 50 µg/ml Phl or 0.5 mM As(III) for 1 or 2 h. Pgk1 and Ponceau S staining served as loading and total-protein controls, respectively, in the immunoblots. (C) Clonogenic survival after 2 h exposure to 0.5 mM As(III), 2.5 µg/ml Phl, or 50 µg/ml Phl, expressed relative to untreated cells. Data are shown as mean ± SD of three biological replicates. Statistical differences were determined by two-tailed Student’s t test. (D) Quantification of Rad52–YFP foci after 1 or 2 h exposure to 0.5 mM As(III), 2.5 µg/ml Phl, or 50 µg/ml Phl. Data are shown as mean ± SD of three biological replicates. Statistical differences were determined by two-tailed Student’s t test. (E) γH2A immunoblot of logarithmically growing WT cells left untreated or exposed to 2.5 µg/ml Phl or 0.5 mM As(III) for 1 or 2 h. (F) PFGE analysis of whole chromosomes from logarithmically growing WT cells after 2 h exposure to 0.5 mM As(III), 2.5 µg/ml Phl, or 50 µg/ml Phl. (G) Rad52–MN ChEC assay in G2/M-arrested cells treated for 2 h with 0.5 mM As(III), 2.5 or 50 µg/ml Phl, in the presence or absence of Ca²⁺. (H) Quantification of Rfa1–YFP foci after 1 or 2 h exposure to 0.5 mM As(III), 2.5 µg/ml Phl, or 50 µg/ml Phl. Data are shown as mean ± SD of three biological replicates. Statistical differences were determined by two-tailed Student’s t test. (I) Spike-in-normalized γH2A ChIP–seq analyses of G2/M arrested WT cells left untreated or exposed to 0.5 mM As(III) for 1 h. Profiles along chromosomes III and IV are shown. A representative biological replicate is shown. (J) Representative DNA-content profiles of logarithmically growing WT cells and nocodazole-arrested G2/M cells before and after exposure to 0.5 mM As(III) for 1 h.

We next sought to further characterize the As(III)-induced checkpoint response and explore potential contributing mechanisms. Because ROS-mediated oxidative damage may drive arsenic genotoxicity (25), we tested its contribution to the acute checkpoint response. In yeast, As(III) caused only minimal increases in oxidative stress (SI Appendix, Fig. S6A) or 8-oxoguanine (31), and neither inhibition of mitochondrial respiration (*ρ0 cells*) nor deletion of *YNO1*, the sole yeast NADPH oxidase (45), affected γH2A accumulation (SI Appendix, Fig. S6 B and C).

Broad γH2A could reflect Mec1 recruitment at numerous lesions or long-range phosphorylation driven by higher-order chromatin organization. To distinguish these possibilities, we depleted cohesin, the major loop-forming complex in *S. cerevisiae* (46). Cohesin depletion did not reduce γH2A (SI Appendix, Fig. S6D), arguing against a major contribution of cohesin-mediated looping to broad As(III)-induced H2A phosphorylation.

We next asked whether this diffuse response was accompanied by overt chromosome fragmentation. To this end, we assessed chromosome integrity using pulsed-field gel electrophoresis (PFGE). PFGE revealed no detectable chromosome smearing after 0.5 mM As(III) or 2.5 µg/ml Phl, whereas high-dose Phl caused pronounced smearing indicative of DNA fragmentation (Fig. 2F). To detect ssDNA-containing intermediates, we used Rad52–MN ChEC. In this assay, Ca²⁺-activated MNase cleaves DNA near Rad52-associated sites, with the resulting fragmentation assessed by genomic DNA electrophoresis. Because pre-existing resected DSBs alone do not increase bulk fragmentation, enhanced DNA smearing primarily reflects Rad52–MN engagement with ssDNA-containing intermediates (47, 48). Phl had little effect on Ca²⁺-dependent digestion, as reported previously (48), whereas As(III) increased DNA fragmentation (Fig. 2G and SI Appendix, Fig. S6E). Rfa1–YFP foci provided an independent readout of ssDNA-associated structures (4, 49). High-dose Phl (50 µg/ml) induced foci in nearly all cells, whereas As(III) produced a modest, approximately twofold increase; notably, 2.5 µg/ml Phl caused an approximately threefold increase relative to untreated cells (Fig. 2H).

Because many genotoxic agents activate the DDC during S phase by inducing replication-fork stalling and ssDNA accumulation (10, 50, 51), we tested whether ongoing DNA replication is required for As(III)-induced γH2A accumulation by arresting cells outside S phase before As(III) exposure. WT cells pre-arrested in G2/M with nocodazole showed a clear γH2A increase after As(III) exposure (Fig. 2 I and J; SI Appendix, Fig. S7A). In α-factor–arrested G1 cells (SI Appendix, Fig. S7B), γH2A induction remained low in WT cells (SI Appendix, Fig. S7C), consistent with yKu-dependent suppression of As(III)-induced DDC activation in G1 (31) but was largely restored by *KU70* deletion (SI Appendix, Fig. S7 D and E). Together, these results argue that ongoing bulk DNA replication at the time of As(III) exposure is not obligatory for γH2A induction.

DDC activation often requires nuclease-mediated formation or extension of ssDNA (11, 52). We therefore tested whether DNA resection contributes to As(III)-induced signaling by examining γH2A and Rad53 phosphorylation in *sgs1*Δ*exo1*Δ or *mre11*Δ mutants. γH2A remained largely WT-like, whereas Rad53 phosphorylation was enhanced in these backgrounds (SI Appendix, Fig. S8 A and B). Consistently, *exo1*Δ and the resection-defective alleles *sgs1-D664*Δ and *mre11-H125N* (53, 54) did not increase As(III) sensitivity (SI Appendix, Fig. S8C).

Taken together, these results indicate that As(III) elicits a characteristic checkpoint response whose overall profile differs from that typically associated with DSBs.

### RNAPII contributes to As(III)-induced γH2A and can trigger widespread signaling when dysfunctional

Previous studies have shown that transcriptional inhibition can suppress DNA damage checkpoint signaling even in nonreplicating cells (55, 56). Given that ongoing bulk DNA replication was not required for As(III)-induced γH2A accumulation, we next asked whether RNAPII contributes to this response.

RNAPII is a conserved, 12-subunit complex whose two largest subunits, Rpb1 and Rpb2, form its catalytic centre. It transcribes protein-coding and many noncoding genes, whereas Pol I and III mainly transcribe more restricted gene classes, including 35S rRNA, 5S rRNA, and tRNAs (57, 58). To test whether RNAPII contributes to As(III)-induced γH2A accumulation, we used an auxin-inducible degron to deplete Rpb1. Although Rpb1 depletion alone elicited only a weak γH2A response, it reduced As(III)-induced γH2A accumulation by approximately 60% (Fig. 3A and B). Flow cytometry revealed comparable cell-cycle distributions in As(III)-treated cells with and without Rpb1 pre-depletion (SI Appendix, Fig. S9A). Importantly, depletion of Rpb2 produced a similar reduction in As(III)-induced γH2A accumulation (SI Appendix, Fig. S9B). To determine whether Rpb1 depletion generally impairs γH2A signaling, we treated cells with Phl. By contrast, Rpb1 pre-depletion increased γH2A levels in cells exposed to phleomycin (Fig. 3C). Together, these findings support a specific contribution of RNAPII to As(III)-induced γH2A signaling.

**Fig. 3.**
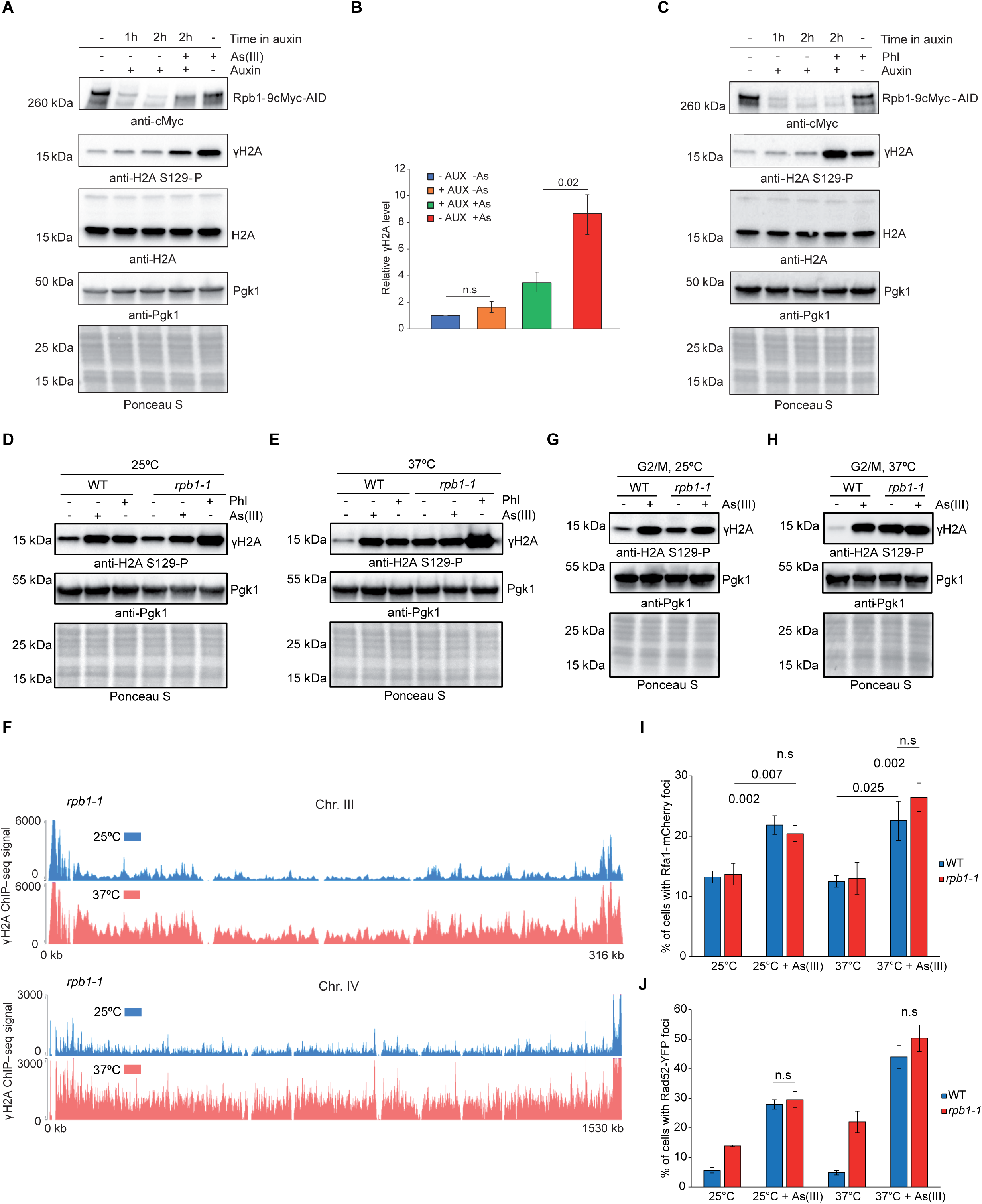
RNAPII depletion attenuates As(III)-induced γH2A, whereas rpb1-1 dysfunction triggers widespread γH2A accumulation. (A) Immunoblot analysis of Rpb1–9cMyc–AID, γH2A, and total H2A. Cells were pretreated with auxin for 1 h to deplete Rpb1, followed by exposure to 0.5 mM As(III) for an additional 1 h in the continued presence of auxin. Control samples included untreated cells, cells treated with auxin alone for 1 or 2 h, and cells exposed to As(III) for 1 h without auxin pretreatment. (B) Densitometric quantification of γH2A levels for four conditions shown in (A): untreated cells, auxin alone (2 h), auxin (2 h) with As(III) added during the final 1 h, and As(III) alone (1 h). γH2A signals were normalized to total H2A and expressed relative to untreated controls, set to 1. Data are presented as mean ± SD from three independent biological replicates. (C) Immunoblot analysis of γH2A and total H2A in Rpb1-AID cells treated as in (A), except that 50 µg/ml Phl was used instead of As(III). (D and E) γH2A immunoblot analysis of logarithmically growing WT and *rpb1-1* cells at 25°C (D) or 37°C (E), left untreated or exposed to 0.5 mM As(III) or 50 µg/ml Phl for 1 h. (F) Spike-in-normalized γH2A ChIP–seq profiles along chromosomes III and IV in *rpb1-1* cells maintained at 25°C or shifted to the restrictive temperature of 37°C. A representative biological replicate is shown. (G and H) γH2A immunoblot analysis of WT and *rpb1-1* cells pre-arrested in G2/M with nocodazole at 25°C (G) or 37°C (H), with or without subsequent exposure to 0.5 mM As(III) for 1 h. Pgk1 and Ponceau S staining served as loading and total-protein controls, respectively, in the immunoblots. (I and J) Quantification of Rfa1–mCherry (I) and Rad52–YFP (J) foci in WT and *rpb1-1* cells at 25°C or 37°C, with or without exposure to 0.5 mM As(III). Data are shown as mean ± SD of three independent biological replicates. Statistical significance was assessed using two-tailed t tests; n.s., not significant.

Because Rpb1 depletion alone induced little γH2A, we tested whether other RNAPII perturbations could reproduce an As(III)-like response. In our previous study (31), neither thiolutin, a transcription inhibitor (59), nor RNAPII inactivation using the temperature-sensitive Rpb1-G1437D (*rpb1-1*) mutant (60–62) measurably altered the As(III)-induced γH2A response (31). We revisited these findings, first examining thiolutin at higher concentrations previously used to inhibit transcription (59, 63, 64). As shown previously (31), 3 µg/ml thiolutin had no detectable effect on γH2A (SI Appendix, Fig. S10A). In contrast, 20 µg/ml thiolutin induced γH2A to levels comparable to As(III) (SI Appendix, Fig. S10B).

Given thiolutin’s pleiotropic effects *in vivo* (59), we next examined RNAPII dysfunction using the *rpb1-1* mutant at permissive (25°C) and restrictive (37°C) temperatures, following an established shift regimen (60–62). At 25°C, WT and *rpb1-1* cells showed low basal γH2A and similar As(III)-induced increases (Fig. 3D). In contrast, shifting cells to 37°C in the absence of As(III) strongly increased γH2A in *rpb1-1* but not WT cells (Fig. 3E), consistent with previous evidence linking *rpb1-1* dysfunction to H2A phosphorylation (14). To determine whether this response is a general consequence of RNAPII dysfunction, we also examined *rpb1-S751F* and *rpb9*Δ cells, which exhibit increased transcriptional slippage and impaired transcriptional elongation, respectively (65, 66). Neither mutant displayed elevated basal γH2A at 30 or 37°C, although both retained γH2A induction following As(III) exposure (SI Appendix, Fig. S10 C and D).

To determine whether *rpb1-1* dysfunction elicits a similarly widespread genomic response, we performed calibrated γH2A ChIP–seq in cells maintained at 25°C or shifted to 37°C. The temperature shift increased the genome-wide mean γH2A signal approximately 3.2-fold (Fig. 3F and SI Appendix, Fig. S11A), while the profiles remained strongly correlated (Spearman ρ = 0.84), indicating amplification of a largely pre-existing distribution. ORF metaprofiles showed increased γH2A across gene bodies and flanking chromatin, with locally reduced signal near transcription start and termination sites (SI Appendix, Fig. S11B). Both *rpb1-1* dysfunction and As(III) treatment of WT cells increased γH2A across nearly the entire analyzed genome (98.6% and 98.4% of bins, respectively), although the magnitudes of these changes were only weakly correlated (Spearman ρ = 0.14). Thus, both inactivation of RNAPII and As(III) exposure elicit widespread γH2A accumulation, but their local patterns of induction differ.

Because both *rpb1-1* dysfunction and As(III) independently induced γH2A, we next tested whether their effects were additive. Addition of 0.5 mM As(III) did not enhance γH2A accumulation in *rpb1-1* cells at 37°C (Fig. 3E). This non-additivity persisted also at 0.25 mM As(III), which induced only weak γH2A in WT cells at 37°C (SI Appendix, Fig. S11 C and D). Non-additivity was unlikely to reflect saturation of H2A phosphorylation, because Phl further increased γH2A in *rpb1-1* cells at 37°C (Fig. 3E). Similarly, methyl methanesulfonate (MMS) or Phl enhanced γH2A in As(III)-treated WT cells, indicating that As(III) does not generally suppress H2A phosphorylation induced by other genotoxins (SI Appendix, Fig. S11 E and F). Interestingly, combined treatment with As(III) and 20 µg/ml thiolutin was also non-additive (SI Appendix, Fig. S10B). Thus, the non-additivity between As(III) and *rpb1-1* dysfunction suggests that these perturbations may converge on overlapping γH2A-inducing processes.

We next examined whether *rpb1-1*-induced γH2A requires ongoing DNA replication. Shifting G2/M-arrested *rpb1-1* cells to 37°C induced strong γH2A without further enhancement by As(III) (Fig. 3G and H). Thus, as in cycling cells, non-additivity persists outside S phase, and neither response requires ongoing bulk DNA replication.

We also examined whether *rpb1-1* dysfunction produces DNA damage-associated features similar to those induced by As(III). PFGE did not reveal detectable DSBs under restrictive conditions (SI Appendix, Fig. S11G). We therefore examined markers associated with ssDNA and HR. After 2 h at 37°C, *rpb1-1* cells showed Rfa1–mCherry focus levels similar to WT, whereas Rad52–YFP foci were elevated at both 25°C and 37°C, consistent with previous observations (14). As(III) further increased both markers, but not beyond the levels observed in As(III)-treated WT cells (Fig. 3 I and J; SI Appendix, Fig. S11H).

RNAPII stalling can promote R-loop formation (67). We therefore investigated their contribution to As(III)-induced γH2A by overexpressing *RNH1*, which encodes an RNase H that degrades aberrantly accumulated R-loops (68). *RNH1* overexpression had no effect on γH2A levels (SI Appendix, Fig. S11I), arguing against a major contribution of R-loops under these conditions. Interestingly, *RNH1* overexpression was previously shown not to reduce the elevated Rad52–YFP foci observed in *rpb1-1* cells (14).

Together, these findings indicate that RNAPII contributes substantially to As(III)-induced γH2A accumulation, prompting us to examine how As(III) affects RNAPII.

### As(III) reduces RNAPII abundance and alters its chromatin distribution

We first examined whether As(III) affects cellular Rpb1 abundance. Immunoblot analysis revealed a progressive decline in Rpb1 levels during As(III) exposure, accompanied by a marked increase in γH2A. Notably, γH2A accumulation was already detectable before substantial Rpb1 loss became apparent (Fig. 4A and SI Appendix, Fig. S12A). Phleomycin also reduced Rpb1 abundance (SI Appendix, Fig. S12 B and C), consistent with previous reports (69, 70), indicating that Rpb1 loss is not specific to As(III). The C-terminal domain (CTD) of Rpb1 undergoes dynamic phosphorylation, with Ser2 and Ser5 phosphorylation associated predominantly with transcription elongation and initiation/early elongation, respectively (71). To assess how As(III) affects RNAPII forms associated with these transcriptional stages, we examined Ser2- and Ser5-phosphorylated Rpb1. Both phospho-specific signals decreased following As(III) exposure, alongside the reduction in total Rpb1 abundance (SI Appendix, Fig. S12 D and E). We next examined whether As(III)-induced Rpb1 loss was accompanied by changes in other RNAPII subunits. Despite a marked reduction in Rpb1 following As(III) exposure, levels of Rpb2 and the noncatalytic core subunit Rpb3 remained largely unchanged (SI Appendix, Fig. S12F). Similarly, auxin-induced Rpb2 depletion had little effect on Rpb1 or Rpb3 abundance (SI Appendix, Fig. S12G).

**Fig. 4.**
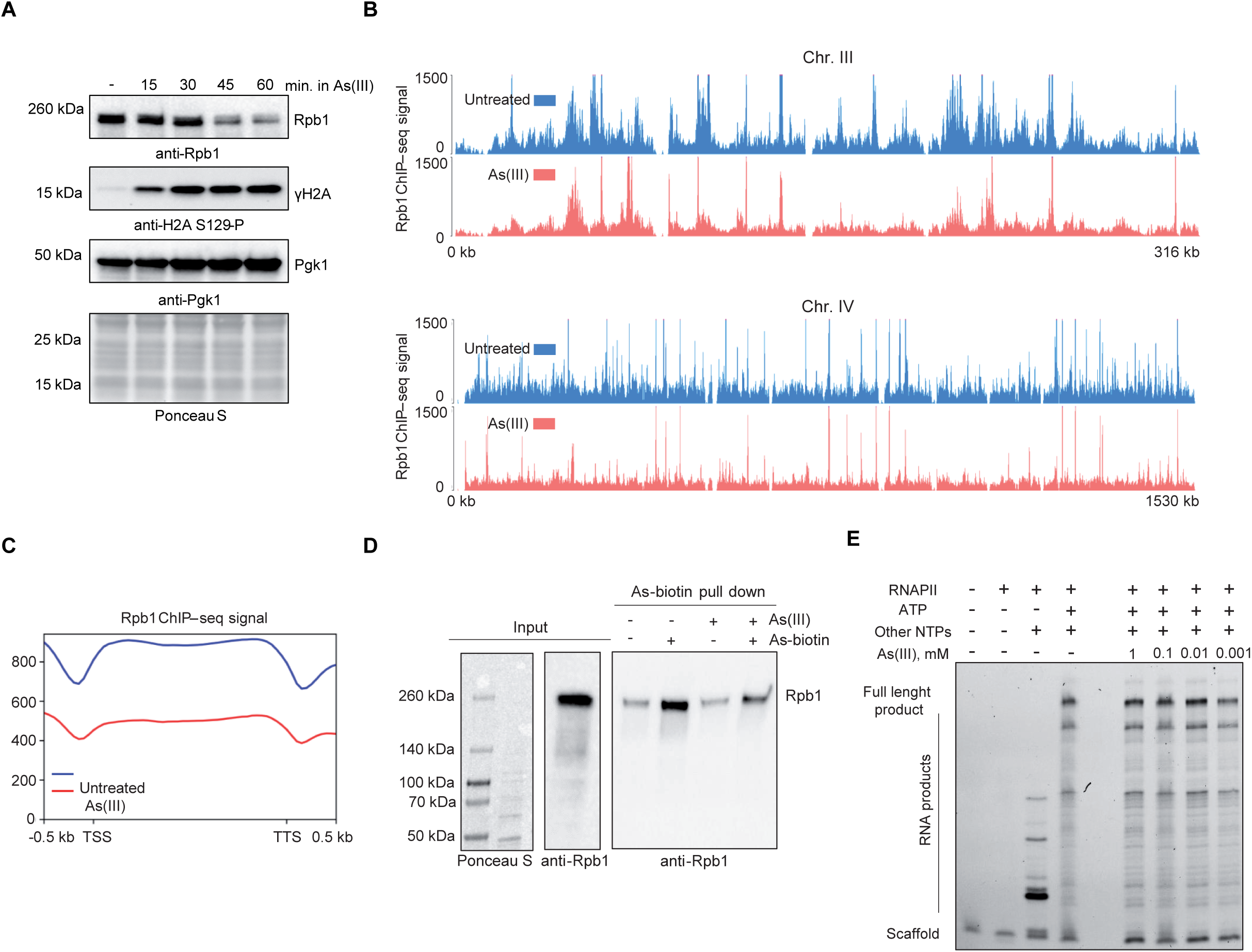
As(III) alters RNAPII abundance and chromatin association. (A) Immunoblot analysis of Rpb1 and γH2A in logarithmically growing WT cells exposed to 0.5 mM As(III) for the indicated times (min). Pgk1 and Ponceau S staining served as loading and total-protein controls, respectively. (B) Spike-in-normalized Rpb1 ChIP–seq profiles along chromosomes III and IV in logarithmically growing WT cells left untreated or exposed to 0.5 mM As(III) for 1 h. A representative biological replicate is shown. (C) Rpb1 ChIP–seq metaprofiles across scaled ORFs under the same conditions. (D) As-biotin pull-down from native yeast cell lysates using 100 µM As-biotin, with or without 100 µM unmodified As(III) as a competitor. Rpb1 recovery was assessed by immunoblotting with anti-Rpb1 antibodies. Input material and pull-down controls are shown. (E) *In vitro* transcription elongation assay using purified yeast RNAPII and a defined nucleic acid scaffold in the presence of the indicated As(III) concentrations. RNA extension products were analyzed by gel electrophoresis.

Next we tested RNAPII chromatin association using Rpb1 ChIP–seq. As(III) exposure caused a widespread reduction in Rpb1 ChIP–seq signal, with the mean signal across ORF bodies decreasing by approximately 35% (Fig. 4B and C; SI Appendix, Fig. S13A). Among 5,764 analyzed genes, 3,945 (68.4%) showed reduced Rpb1 occupancy (log₂FC ≤ −0.5), whereas only 124 (2.2%) exhibited increased occupancy (log₂FC ≥ +0.5). The remaining 1,695 genes (29.4%) showed relatively minor changes in Rpb1 occupancy (−0.5 < log₂FC < +0.5) (SI Appendix, Dataset S1). Gene Ontology (GO) enrichment analysis revealed that genes with increased Rpb1 occupancy following As(III) exposure were enriched for sulfur compound metabolism, sulfate assimilation, glutathione metabolism, detoxification, and oxidative stress responses. In contrast, genes with reduced Rpb1 occupancy were enriched for ribosome biogenesis, rRNA processing, and mRNA processing (SI Appendix, Dataset S2). These patterns are consistent with previously reported transcriptional responses to arsenite (40, 72).

To investigate whether pre-existing RNAPII occupancy was associated with the genomic distribution of As(III)-induced γH2A, we compared basal Rpb1 occupancy with γH2A levels before and after As(III) exposure. Basal Rpb1 occupancy positively correlated with γH2A levels following As(III) treatment (Spearman’s ρ = 0.60), whereas only a weak negative correlation was observed under untreated conditions (ρ = −0.12). These findings indicate that genomic regions with higher basal RNAPII occupancy tend to exhibit higher γH2A levels following As(III) exposure.

Based on the findings described above, we hypothesized that As(III) may impair RNAPII function. Several RNAPII subunits, including Rpb1, harbour cysteine-rich zinc-binding motifs, raising the possibility that these regions may interact with arsenic (71, 73). To investigate this, we purified endogenous RNAPII from *S. cerevisiae* (SI Appendix, Fig. S13B) and incubated it with N-biotinyl p-aminophenyl arsenic acid (As-biotin). Streptavidin-captured proteins were subsequently analysed by mass spectrometry. As-biotin increased the recovery of Rpb1 and Rpb2 compared with beads-only controls, while several additional RNAPII subunits, including Rpb3, Rpb5, Rpb9, Rpb11, and Rpb4, were also detected (SI Appendix, Fig. S13C). To further examine this association in a more physiological environment, we performed As-biotin pull-downs from native yeast lysates. Immunoblot analysis confirmed the recovery of Rpb1, which was reduced in the presence of unmodified As(III) (Fig. 4D). Together, these findings support an arsenic-dependent association with RNAPII-containing material, although they do not establish direct binding to a specific RNAPII subunit *in vivo*.

We next asked whether As(III) directly affects RNAPII catalytic activity. An *in vitro* transcription assay revealed no detectable impairment of RNA extension in the presence of As(III) (Fig. 4E and SI Appendix, Fig. S13D). Thus, despite the association detected using As-biotin, As(III) did not measurably inhibit RNAPII elongation under the conditions tested.

### Short-term As(III) exposure elicits a γH2AX response in human cancer cells

Core features of DNA damage checkpoint signaling are conserved from yeast to humans. ATM (Tel1 in yeast) responds primarily to DSBs, whereas ATR (Mec1 in yeast) is recruited through ATRIP (Ddc2 in yeast) to RPA-coated ssDNA generated during replication stress and other DNA-processing events. Both kinases phosphorylate H2AX on Ser139 (γH2AX), analogous to phosphorylation of yeast H2A on Ser129 (1). Because As(III)-induced γH2AX in human cells has mainly been described after prolonged exposure and linked to ROS (24, 74, 75), we asked whether acute exposure elicits an early checkpoint response. Exposure of A549 cells to 12.5 μM As(III) induced γH2AX within 1 h (Fig. 5 A and E). Using phosphorylation of ATM Ser1981 and ATR Thr1989 as activation readouts, we found that, unlike Phl, As(III) increased ATR Thr1989 phosphorylation without a detectable increase in ATM Ser1981 phosphorylation and caused a modest increase in RPA foci (Fig. 5 B–E). This response occurred without a significant increase in ROS (Fig. 5F), arguing against oxidative stress as its dominant early trigger. We next tested whether transcriptional perturbation could evoke a similar γH2AX response. Combined inhibition of transcription initiation and elongation with triptolide, an XPB/TFIIH inhibitor, and DRB (5,6-dichloro-1-β-D-ribofuranosylbenzimidazole), a CDK9/P-TEFb inhibitor, induced γH2AX. Addition of As(III) did not further increase the signal under these conditions (Fig. 5G). Further supporting a connection to RNAPII, As(III)-biotin recovered endogenous RPB1 from native cell lysates, and unmodified As(III) reduced this recovery (Fig. 5H). Together, these findings indicate that acute As(III) exposure induces ATR activation accompanied by an early γH2AX response, consistent with ssDNA-associated signaling under low-ROS conditions.

**Fig. 5.**
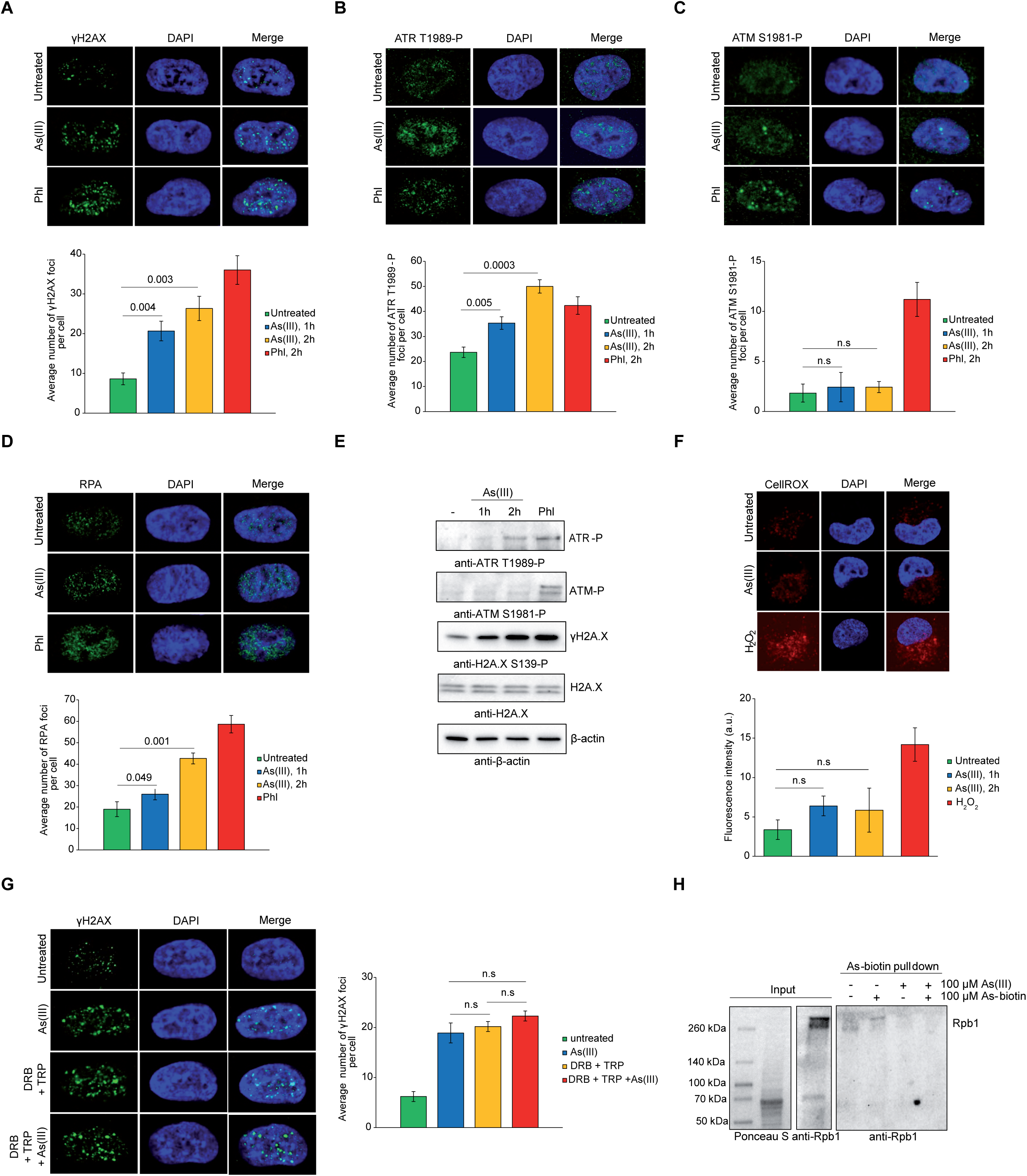
Acute As(III) activates ATR signaling and γH2AX with minimal ROS induction. (A–D) Immunofluorescence analysis of γH2AX (Ser139) (A), phosphorylated ATR (Thr1989) (B), phosphorylated ATM (Ser1981) (C), and RPA (D) in A549 cells left untreated or exposed to 12.5 µM As(III) for 1 or 2 h or 15 µg/ml Phl for 2 h. Top, representative images of untreated cells and cells treated with As(III) or Phl for 2 h. Bottom, quantification of foci per cell from more than 200 cells per condition across three independent biological experiments (n = 3). Data are shown as mean ± SD. Statistical significance was assessed using two-tailed Student’s t tests; n.s., not significant. (E) Immunoblot analysis of phosphorylated ATR (Thr1989), phosphorylated ATM (Ser1981), γH2AX, and total H2AX in A549 cells left untreated or exposed to 12.5 µM As(III) for 1 or 2 h or 15 µg/ml Phl for 2 h. β-actin served as a loading control. (F) CellROX DeepRed–based detection of ROS in A549 cells left untreated or exposed to 12.5 µM As(III) for 1 or 2 h or 100 µM H₂O₂ for 30 min. Top, representative images. Bottom, quantification of CellROX Deep Red fluorescence intensity from more than 100 cells per condition across three independent biological experiments (n = 3). Data are shown as mean ± SD. Statistical significance was assessed using two-tailed Student’s t tests; n.s., not significant. (G) Immunofluorescence analysis of γH2AX in A549 cells pretreated with the transcription inhibitors triptolide (TRP, 1 µM) and DRB (100 µM) for 1 h, followed by exposure to 12.5 µM As(III) for 2 h. Untreated cells, As(III) alone, and TRP + DRB without As(III) served as controls. Left, representative images. Right, quantification of γH2AX foci per cell from 200 cells per condition across three independent biological experiments (n = 3). Data are shown as mean ± SD. Statistical significance was assessed using two-tailed Student’s t tests; n.s., not significant. (H) Streptavidin pull-down from native A549-cell lysates using 100 µM As-biotin, with or without 100 µM unmodified As(III) as a competitor, followed by immunoblotting for endogenous RPB1. Input material and pull-down controls are shown.

## DISCUSSION

Arsenic genotoxicity is commonly attributed to ROS-mediated DNA damage and impaired DNA repair, potentially leading to replication-associated lesions (24–26). Here, we identify an important contribution of RNAPII-associated processes to early As(III)-induced checkpoint signaling, suggesting an additional mechanism linking arsenic exposure to genome stress. The early As(III)-induced checkpoint response differs from that elicited by phleomycin.

At doses inducing comparable γH2A levels, As(III) produces substantially fewer RPA and Rad52 foci, without detectable chromosome fragmentation or loss of viability, indicating that similar checkpoint responses need not reflect comparable burdens of canonical DSBs. Moreover, γH2A induction by As(III) does not require ongoing bulk DNA replication or extensive DNA end resection, suggesting that the response differs from conventional replication-associated DSB signaling. The absence of prominent new RPA- or HR-factor-enriched domains, typically observed around defined DSB sites, may reflect transient or spatially heterogeneous checkpoint-inducing events that are difficult to resolve by population-averaged ChIP-seq.

Although ROS-mediated DNA damage is widely considered a major contributor to arsenic genotoxicity, recent studies have questioned the role of mitochondrial ROS as a direct source of chromosomal DNA damage. Notably, sustained chemogenetic production of mitochondrial H₂O₂, even at supraphysiological levels, failed to induce detectable DNA strand breaks or DDR activation in viable human cells, even during mitosis (76) and similarly did not induce γH2A accumulation in fission yeast (77). Moreover, at least in some cellular models, ROS accumulation can be uncoupled from arsenic-induced cytotoxicity, as antioxidant interventions have been shown to suppress ROS without preventing cell death (78, 79). In our study, As(III) induced only minor changes in oxidative stress markers, while previous measurements revealed little 8-oxoguanine accumulation (31). Furthermore, neither loss of mitochondrial respiration nor deletion of *YNO1* attenuated γH2A induction. Together, these findings suggest that ROS-mediated oxidative DNA damage is unlikely to be the principal driver of early As(III)-induced checkpoint signaling, although oxidative stress may contribute to arsenic genotoxicity during prolonged exposure.

Unexpectedly, our findings implicate RNAPII-associated processes in early As(III)-induced checkpoint signaling. The contrasting effects of Rpb1 depletion, transcriptional shutdown in *rpb1-1*, and defects in transcription elongation and fidelity (*rpb1*-S751F and *rpb9*Δ) suggest that checkpoint activation may depend on particular states of RNAPII rather than a general impairment of transcription. Rpb1 loss itself is also unlikely to initiate this response, as γH2A accumulation precedes substantial Rpb1 loss following As(III) exposure, while *rpb1-1* RNAPII can remain chromatin-associated after inactivation (14, 61). Moreover, phleomycin also reduces Rpb1 abundance, indicating that this effect is not specific to arsenic. Our biochemical findings suggest that As(III) may associate with the RNAPII complex without necessarily inhibiting RNA elongation. Indeed, arsenic can impair certain functions of a protein while leaving others intact. For example, arsenic sulfide inhibits autoubiquitination of the E3 ligase c-CBL without abolishing its ability to ubiquitinate the oncogenic fusion kinase BCR–ABL (80). Similarly, arsenic disrupts intramolecular interactions within the androgen receptor and reduces its recruitment to chromatin, without affecting ligand binding, nuclear translocation, or DNA binding *in vitro* (81). These observations raise the possibility that As(III) induces conformational changes in RNAPII, potentially affecting its interactions with transcription factors, chromatin association, or transitions between functional states without necessarily inhibiting RNA elongation. Such perturbations could promote the formation or exposure of aberrant ssDNA-containing structures.

Importantly, rapid As(III)-induced checkpoint activation under conditions of limited oxidative stress is not restricted to yeast. In human A549 cells, acute As(III) exposure induces γH2AX accumulation and ATR phosphorylation without detectable ATM activation or a substantial increase in ROS. The non-additive γH2AX response to As(III) and transcription inhibitors, together with arsenic-affinity recovery of human RPB1, raises the possibility that RNAPII-associated processes may also contribute to checkpoint activation in mammalian cells. However, a functional requirement for RNAPII remains to be established. The evolutionary conservation of RNAPII, including cysteine-coordinated zinc-binding motifs in its subunits (82, 83), raises the possibility that As(III) may perturb similar structural features of the complex in different organisms. Beyond affecting transcription, such perturbations could compromise DNA repair, given the established role of RNAPII in transcription-coupled repair and evidence that transcription promotes HR at DSBs (84). Reduced repair efficiency could, in turn, promote the persistence of DNA lesions and genome instability. Furthermore, arsenic-induced reductions in mRNA and protein abundance reported in human cells (85–87) may partly reflect RNAPII dysfunction, alongside other cellular stress responses.

Although our findings are consistent with ssDNA-associated checkpoint signaling, the origin of the underlying DNA structures remains uncertain. The transcription bubble itself represents one possible source of ssDNA, although its limited size and partial enclosure within RNAPII may constrain its ability to elicit a robust checkpoint response (88). An alternative possibility involves transcription-associated DNA topology. Transcription generates positive supercoiling ahead of RNAPII and negative supercoiling behind it, normally relaxed by topoisomerases (89). As(III)-induced RNAPII dysfunction could alter the balance between transcription-generated supercoiling and topoisomerase-mediated relaxation, favoring the local accumulation of torsional stress. Excessive negative supercoiling could promote local DNA unwinding and ssDNA exposure, as demonstrated in transcription-associated systems in yeast and mammalian cells (90–92). Conversely, accumulated positive supercoiling could further impede RNAPII elongation (93). Such locally exposed ssDNA could bind RPA and recruit Mec1–Ddc2, potentially promoting γH2A phosphorylation. This model could explain why As(III)-induced γH2A accumulation does not require ongoing DNA replication or extensive DNA end resection, as transcription persists outside S phase and topologically driven DNA unwinding could expose ssDNA without nucleolytic processing. Moreover, numerous transient and spatially heterogeneous transcription-associated perturbations could account for widespread γH2A signaling independently of cohesin-mediated chromatin looping or substantial burden of canonical DSBs. Whether As(III) induces such topological alterations *in vivo* and whether they are sufficient to activate checkpoint signaling remain to be determined.

Taken together, our findings identify a substantial contribution of RNAPII-associated processes to early As(III)-induced checkpoint signaling, revealing an additional link between transcriptional perturbation and the cellular response to environmental genotoxic stress.

## MATERIALS AND METHODS

### Yeast strains

The *Saccharomyces cerevisiae* strains used in this study are listed in Table S1. Gene deletion or tagging was performed using a PCR-based method (94–96). Multiple gene mutants were obtained by genetic crossing of relevant haploids followed by tetrad dissection.

### Mammalian cells

A549 cells were sourced from the cell line collection of the Department of Molecular Physiology and Neurobiology, University of Wrocław. A549 cells are commonly used in toxicology studies and have been widely applied to investigate cellular responses to arsenic exposure.

### Yeast growth media

*S. cerevisiae* strains were grown at 30°C in YPD medium (1% yeast extract, 2% peptone, 2% glucose). The *rpb1-1* mutant was maintained at 25°C and, for experiments at the restrictive temperature, preincubated at 37°C for 30 min before further incubation at the same temperature with or without As(III) for 1 h. *S. pombe* spike-in cultures were grown at 30°C in YES medium (0.5% yeast extract, 3% glucose, supplemented with 225 mg/l adenine, leucine, and uracil). Solid media contained 2% agar. As(III) was used in the form of sodium (meta)arsenite (NaAsO₂; Sigma-Aldrich, S7400). Auxin-induced degradation (AID) was induced by the addition of 0.5 mM 3-indoleacetic acid (IAA; Sigma-Aldrich, I2886).

### Mammalian cell culture conditions

A549 cells were maintained in low-glucose DMEM (Biowest, L0064) supplemented with 10% fetal bovine serum (Biowest, S181H-500), 100 U/ml penicillin and 100 µg/ml streptomycin, and 2 mM L-glutamine. Cells were cultured at 37°C in a humidified atmosphere containing 5% CO₂. Unless stated otherwise, cells were grown for 48 h under these standard conditions prior to the start of experiments. As(III) was used in the form of sodium (meta)arsenite (NaAsO₂; Sigma-Aldrich).

### Yeast serial dilution growth assay (spot tests)

To assess the sensitivity of the relevant strains to As(III) or other DNA-damaging agents, mid-log cultures were normalized to the same OD600, 10-fold serially diluted, and spotted onto solid media containing the indicated drug concentrations. Spotting assays with As(III) were performed on synthetic complete (SC) medium (0.67% yeast nitrogen base without amino acids, supplemented with 225 mg/l adenine, leucine, histidine, tryptophan, and uracil, and 2% glucose). Spotting assays with Phl or MMS were performed on YPD plates. For each strain, several independent clones were tested. All serial dilution growth assays were performed at least three times, and representative images are shown.

### Yeast survival assay

Log-phase cultures were left untreated or exposed to the indicated DNA-damaging agent. Cells were then washed, serially diluted, and plated on solid agar plates. After 3 days of incubation at 30°C, colonies were counted. Survival was expressed as the percentage of colony-forming units (CFUs) relative to the corresponding time-zero sample, which was set to 100%. Data represent the mean ± standard deviation (SD) from at least three independent biological replicates.

### Cell cycle synchronization and analysis

To arrest yeast in G1 phase, cells were cultured in YPD in the presence of 5 µM α-factor for 2 h. G2/M block was achieved by incubating cells in YPD containing 15 µg/ml nocodazole (Sigma-Aldrich, M1404) for 2 h. G1 and G2/M arrests were confirmed by microscopic observations and FACS analysis. To measure DNA content, cell samples were fixed with 70% ethanol. Next, cells were digested with 0.25 mg/ml RNase (Thermo Scientific, EN0531) for 2 h at 50°C and with 5 mg/ml pepsin for 1 h at 37°C. Then, cells were sonicated and stained with 0.5 µM Sytox Green (Invitrogen, S7020) for 30 min in the dark. FACS analysis was performed using a Millipore Guava EasyCyte flow cytometer.

### Yeast whole-protein extracts isolation

Protein extracts were prepared using a trichloroacetic acid (TCA) precipitation protocol. Approximately 3 × 10⁷ cells of the indicated strains were harvested and incubated in 100 µl lysis buffer (2 M NaOH, 7% β-mercaptoethanol) for 1 min. Then, 100 µl of 50% TCA was added, and samples were mixed and centrifuged. The supernatant was discarded, and the pellet was washed with 1 M Tris-HCl, pH 8.0. The resulting protein pellet was resuspended in Laemmli sample buffer (50 mM Tris-HCl, pH 6.8, 2% SDS, 10% glycerol, 2% β-mercaptoethanol) and boiled for 5 min. For immunoblot analysis, 3 µl of each protein extract were routinely loaded per lane on SDS–PAGE gels.

### Isolation of whole-cell protein extracts from mammalian cells

A549 cells were seeded in 10-cm tissue culture dishes at 5 × 10⁵ cells per dish and cultured for 48 h before treatment with the indicated compounds. After treatment, dishes were washed on ice with ice-cold PBS, and cells were scraped directly into ice-cold RIPA buffer (50 mM Tris-HCl, pH 7.4, 150 mM NaCl, 1% NP-40, 0.5% sodium deoxycholate, 0.1% SDS, 1 mM MgCl₂) supplemented with 1 mM PMSF, protease inhibitor cocktail (Thermo Fisher, 78441), and 1 KU/ml Benzonase (Sigma-Aldrich, E1014). Collected cells were incubated on ice for 30 min and then centrifuged at 16,000 × g for 10 min at 4°C. The resulting whole-cell lysate was mixed 1:1 with 2x Laemmli sample buffer and boiled for 5 min.

### Immunoblotting

Protein extracts were routinely resolved on 4–20% gradient SDS–PAGE gels (Bio-Rad, 4568096) and transferred onto nitrocellulose membranes by semi-dry transfer. For analysis of Rad53 phosphorylation, proteins were separated on 7.5% SDS–PAGE gels (Bio-Rad, 4568026). Membranes were blocked in 5% non-fat dry milk in TBST (TBS: 20 mM Tris-HCl, 150 mM NaCl, pH 7.5, 0.1% Tween-20) and incubated with primary antibodies for 2 h, followed by incubation with secondary antibodies for 1 h in 1% milk in TBST. Signal was detected using a ChemiDoc MP imaging system (Bio-Rad). The following antibodies were used for Western blotting: 1:2000 anti-H2A S129-P (Abcam, ab15083), 1:5000 anti-H2A (Active-Motif, 39235), 1:20000 anti-Pgk1 (Abcam, ab113687), 1:2000 anti-Rad53 (Abcam, ab104232), 1:2000 anti-cMyc (Roche, 11667149001), 1:3000 anti-Pk (Bio-Rad, MCA1360G), 1:3000 anti-HA (Roche, 11867423001), 1:1000 anti-Rpb1 (Sigma-Aldrich, 05-952-I), 1:1000 anti-Rpb1 S2-P (Abcam, ab5095), 1:1000 anti-Rpb1 S5-P (Abcam, ab240740), 1:2000 anti-Rpb3 (Abcam, ab202893), 1:1000 anti-ATR T1989-P (Thermo Fisher, MA5-45973), 1:1000 anti-ATM S1981-P (Thermo Fisher, MA1-2020), 1:2000 anti-H2A.X S139-P (Cell Signaling, 9718), 1:1500 anti-H2A.X (Proteintech, 10856-1-AP), 1:4000 anti-β-actin (Sigma-Aldrich, A5441), 1:5000 anti-mouse (Jackson ImmunoResearch, 115-035-003), 1:5000 anti-rabbit (Jackson ImmunoResearch, 111-035-003). All experiments were performed at least three times. Representative images are shown. Densitometric analysis of γH2A immunoblot signals was performed using Image Lab software (Bio-Rad). γH2A signal intensities were normalized to the corresponding total H2A signals. Quantification was based on three independent biological replicates.

### Live yeast cell imaging

Yeast cells were observed with the Axio Imager M1 epifluorescence microscope (Carl Zeiss) equipped with a 100x immersion oil objective (Plan-Neofluar 100x / 1.30), the CFP, GFP and dsRED filter set and differential interference contrast (DIC). Z-stacked images were collected using an AxioCam MRc digital colour camera and processed with ZEN 3.7 software (Carl Zeiss). Data represent the mean ± standard deviation (SD) from at least three independent biological replicates. For microscopy-based quantification, statistical analyses were performed using the mean values from three independent biological experiments (n = 3).

### Immunofluorescence

A549 cells were seeded on glass coverslips placed in 12-well plates at 1.5 × 10⁴ cells per well and cultured for 48 h before fixation. Cells were fixed with 4% paraformaldehyde (PFA) in PBS for 12 min at 4°C, washed with PBS (3 × 5 min, RT), and permeabilized with 0.1% Triton X-100 in PBS for 3 min, followed by an additional PBS wash (3 × 5 min, RT). Coverslips were then blocked in 5% BSA in PBS for 60 min at RT and incubated with primary antibodies [1:200 anti-ATR T1989-P (Thermo Fisher, MA5-45973), 1:250 anti-ATM S1981-P (Thermo Fisher, MA1-2020), 1:800 anti-H2A.X S139-P (Cell Signaling, 9718), 1:200 anti-RPA32 (Abcam, ab76420)] overnight at 4°C. After incubation, coverslips were washed with PBS (3 × 5 min, RT). Primary antibodies were detected using fluorophore-conjugated secondary antibodies: goat anti-mouse Alexa Fluor 488 (1:1500; Thermo Fisher, A-11001) and goat anti-rabbit Alexa Fluor 488 (1:1000; Thermo Fisher, A-11034) for 60 min at RT. Following secondary antibody incubation, coverslips were washed again with PBS (3 × 5 min, RT) and mounted using Fluoroshield with DAPI (Merck, F6057). Stained cells were imaged using an Olympus FV1000 confocal microscope equipped with filter sets for DAPI and FITC/Alexa Fluor 488. Images were acquired with a Plan Apo 60×/1.4 NA oil-immersion objective. DAPI and Alexa Fluor 488 were excited with 405 nm and 473 nm diode lasers, respectively.

### ROS measurements in yeast

To analyse ROS levels in *S. cerevisiae*, WT cells were pre-loaded with MitoSOX (Thermo Fisher, M36005), CellROX (Thermo Fisher, C10444), 2′,7′-dichlorofluorescein (Sigma-Aldrich, 35845), or dihydroethidium (Sigma-Aldrich, D11347) at 5 µM for 15 min and then exposed to As(III). After 60 min, aliquots of cells were collected and immediately analyzed by flow cytometry using a Millipore Guava EasyCyte cytometer to measure fluorescence intensity. Untreated samples were used to determine baseline autofluorescence. Representative results are shown; experiments were performed at least three times.

### ROS measurements in mammalian cells

A549 cells were seeded on glass coverslips placed in 12-well plates at 1.5 × 10⁴ cells per well and cultured for 48 h. Cells were then incubated with 100 µM H₂O₂ in HBSS (Gibco, 14180046) for 30 min (positive control) or with 12.5 µM As(III) for 1 h or 2 h. After treatment, cells were washed once with HBSS. ROS staining was performed by incubating cells with 5 µM CellROX Deep Red (Thermo Fisher, C10422) in HBSS for 30 min at 37°C in the dark. Cells were then washed with HBSS, fixed with 4% PFA in PBS for 12 min at 4°C, washed again with PBS, and mounted using Fluoroshield mounting medium containing DAPI (Merck). Stained cells were imaged using an Olympus FV1000 confocal microscope equipped with filter sets for DAPI and Cy5.

### Arsenic uptake measurement

Approximately 1 × 10⁸ cells were treated or not with 0.5 mM As(III) for 1 h or 2 h. At time zero and after As(III) incubation, aliquots were plated on YPD agar to determine viable cell numbers. At the end of the treatment, 0.1% (w/v) sodium azide was added, cells were pelleted, and washed three times with distilled water. Pellets were resuspended in 800 µl ST buffer (1 M sorbitol, 20 mM Tris-HCl, pH 7.5), supplemented with 200 µl zymolyase solution (10 mg/ml in ST buffer), and incubated at 30°C for 55 min. β-mercaptoethanol was then added to a final concentration of 0.05 M and samples were incubated for an additional 5 min at 30°C. Cells were washed, resuspended in 2 ml of 0.5% (v/v) nitric acid (EMPLURA, Supelco, 1.00443.2500), and boiled for 10 min. Suspensions were centrifuged (15 min, 3,320 × g, 4°C) and arsenic concentrations in the supernatants were determined by atomic absorption spectrometry using a graphite furnace (contrAA 800 G, Analytik Jena). For calculations of intracellular arsenic concentration, an average yeast cell volume of 65 fL was assumed (97). Arsenic uptake measurements were performed three times in triplicate, and error bars represent mean ± standard deviation (SD).

### Chromatin endogenous cleavage (ChEC)

Approximately 1 × 10⁸ Rad52–MNase cells per sample were arrested with 0.1% sodium azide and permeabilized with 1% digitonin for 5 min. To activate micrococcal nuclease, cells were then incubated with 2 mM CaCl₂ at 30°C for 30 min. DNA was extracted by phenol–chloroform purification followed by ethanol precipitation. Equal amounts of total DNA were loaded, resolved on a 0.8% agarose gel, and stained with ethidium bromide. DNA intensities in each lane were quantified using a ChemiDoc MP imaging system (Bio-Rad) and Image Lab software and plotted as histograms. The experiment was repeated three times with similar results, and representative data are shown.

### Isolation of whole yeast chromosomes and pulsed-field gel electrophoresis (PFGE)

Preparation of agarose-embedded genomic DNA was performed using the CHEF Genomic DNA Plug Kit (Bio-Rad, 1703593) according to the manufacturer’s instructions with minor modifications. To ensure equal DNA content in the plugs, samples were collected at each time point, cell numbers were determined with a hemocytometer, and adjusted to 3 × 10⁷ cells per sample. Cells were embedded in 50 µl of 0.8% low-melting agarose and digested with 1 mg/ml lyticase for 2 h at 37°C, followed by overnight incubation with 1 mg/ml proteinase K at 50°C. Plugs were then washed four times in washing buffer (20 mM Tris-HCl, pH 8.0, 50 mM EDTA) and loaded onto the gel. PFGE was carried out using a CHEF-DR III system (Bio-Rad). Chromosomes were resolved in 1% agarose at 6 V/cm for 24 h with a 1–2 min switch time ramp and a 120° switch angle at 14°C. Gels were stained with ethidium bromide and imaged using a ChemiDoc MP system (Bio-Rad) and Image Lab software.

### Chromatin immunoprecipitation (ChIP)

For calibrated ChIP, approximately 4 × 10⁸ *S. cerevisiae* cells were mixed with ∼1 × 10⁸ *Schizosaccharomyces pombe* cells expressing Rad21–6HA, fixed in 1% formaldehyde for 15 min at room temperature (RT), and quenched with 150 mM glycine for 5 min. Cells were pelleted, washed once with TBS, and resuspended in 600 µl FA lysis buffer (50 mM HEPES-KOH, pH 7.5, 140 mM NaCl, 1 mM EDTA, 0.1% sodium deoxycholate, 1% Triton X-100, protease inhibitor cocktail [Sigma-Aldrich, P8215], 1 mM PMSF). Cells were lysed by four cycles of bead beating (6,500 rpm, 30 s) and lysates were sonicated to an average DNA fragment length of ∼600 bp. After centrifugation for 30 min at 15,000 rpm, cleared supernatants were transferred to new tubes and 1% of each was kept as an input sample. For immunoprecipitation, 5 µl of antibody against the *S. cerevisiae* protein of interest [anti-histone H2A phospho S129 (Abcam, ab15083), anti-histone H2A phospho S129 (Active Motif, 39271), anti-c-myc (Roche, 11667149001), anti-HA (Roche, 11867423001), anti-Rad51 (Invitrogen, PA534905), anti-Rpb1 (Sigma-Aldrich, 05-952-I), anti-RPA (Agrisera, AS07 214), anti-GFP (Invitrogen, A-11122)] and 2 µl of anti-HA (Roche, 11867423001) recognizing Rad21–6HA were added to the lysates, and samples were incubated overnight at 4°C with rotation. The next day, DNA-protein complexes were captured with Protein G Dynabeads (Invitrogen, 10004D) and sequentially washed with FA-lysis buffer, FA-500 buffer (50 mM HEPES–KOH pH 7.5, 500 mM NaCl, 1% Triton X-100, 0.1% sodium deoxycholate, 1 mM EDTA), LiCl wash buffer (10 mM Tris–HCl pH 8.0, 250 mM LiCl, 0.5% NP-40, 0.5% sodium deoxycholate, 1 mM EDTA) and TE (10 mM Tris–HCl pH 7.5, 1 mM EDTA) on a rotator for 5 min each time. Beads and input samples were then resuspended in elution buffer (50 mM Tris–HCl pH 7.5, 1% SDS, 10 mM EDTA) and incubated O/N at 65°C on a thermal shaker. The next day, magnetic beads were removed, and all samples were treated with RNase A (100 µg/ml; Thermo Scientific, EN0531) for 1 h at 37°C, followed by a 1 h incubation at 65°C in the presence of proteinase K (1 mg/ml). DNA from H2A-S129-P and Rpb1 ChIP samples was purified using a ChIP DNA Clean & Concentrator kit (Zymo Research, D5205). Sequencing libraries for these samples were prepared using the Novogene NGS DNA Library Prep Set (Novogene, PT004). DNA from RPA, Rad52–YFP, Rad51 and Rad59–13cMyc ChIP samples was extracted by phenol–chloroform purification and ethanol precipitation. Libraries were generated using the xGen™ ssDNA & Low-Input DNA Library Preparation Kit (Integrated DNA Technologies, 10009859) according to the manufacturer’s instructions. Libraries were sequenced by Novogene on a NovaSeq X Plus platform. All ChIP–seq experiments were conducted in at least two independent biological replicates, with the exception of H2A S129-P ChIP–seq performed once in H2A S129A cells, WT and *ku70*Δ cells arrested in G1 as well as in WT cells treated with Phl. Datasets have been deposited in the Gene Expression Omnibus (GEO) under accession number GSE318138. Chromatin immunoprecipitation for ChIP–qPCR was performed as described above, followed by quantitative PCR as previously described (98).

### ChIP-seq data processing and analysis

Paired-end ChIP-seq reads were processed using Cutadapt (99) to remove adapter sequences and low-quality terminal bases (Phred quality threshold of 30). Reads shorter than 60 bp after trimming were discarded. Filtered ChIP and input reads were aligned to a combined reference genome of *Saccharomyces cerevisiae* (sacCer3) and *Schizosaccharomyces pombe* (ASM294v2) using Bowtie2 (100). Alignment was performed using the *--very-sensitive* preset with a maximum fragment length of 1,000 bp. Alignment files were sorted and processed using SAMtools (101). Properly paired, primary alignments were retained, while unmapped, secondary, and supplementary alignments were excluded. PCR duplicates were marked using samtools markdup. Reads were filtered according to mapping quality (MAPQ), using thresholds of ≥10 for Rad51, Rad52, and Rad59; ≥20 for phosphorylated H2A (H2A-P) and RPA; and ≥30 for Rpb1. The MAPQ threshold of 20 was also applied to H2A-P ChIP-seq datasets obtained from *ku70*Δ cells. For analyses involving telomeric regions, the MAPQ threshold was set to 0 to retain reads mapping to repetitive sequences. Alignments corresponding to the chromosomes of *S. cerevisiae* and *S. pombe* were subsequently separated, excluding mitochondrial sequences. Spike-in normalization was performed as described by Hu *et al*. (102), using reads mapped to the *S. cerevisiae* and *S. pombe* genomes in both ChIP and input samples. Spike-in normalization and peak calling were performed using MACS3 (103) through a custom computational pipeline. Duplicate reads were retained during peak calling using the *--keep-dup all* option. To estimate background signal, MACS3 was used to model local background using multiple window sizes (1 kb, 10 kb, the estimated fragment length d, and a genome-wide background). The normalized ChIP signal and local background were compared using *macs3 bdgcmp* to calculate Poisson-based p-values, adjusted q-values, and fold enrichment. Narrow peaks were identified using bdgpeakcall, whereas broad enriched regions were identified using bdgbroadcall. Peak calling was performed separately using p-value- and q-value-based significance tracks, with a threshold of 0.05 for narrow and strongly enriched regions and 0.10 for weaker regions contributing to broad peaks. The minimum peak length was set to the estimated fragment length (d), with maximum gap lengths of 150 bp for narrow and strongly enriched regions and 600 bp for weaker broad regions. For visualization and downstream analyses, spike-in-normalized bigWig signal tracks (104) were generated from *S. cerevisiae* ChIP and corresponding input alignments using bamCoverage from deepTools (105). Sample-specific spike-in normalization factors were applied to account for differences in spike-in representation between samples. Coverage was calculated in 25-bp bins. Duplicate reads were retained during bigWig generation, as the *--ignoreDuplicates* option was not used. Mapped reads, normalized coverage tracks, and identified peaks were inspected using the Integrative Genomics Viewer (IGV). Metagene profiles, heatmaps, and additional visualizations were generated using deepTools.

For MNase-seq analysis, raw sequencing data from wild-type cells were retrieved from the Gene Expression Omnibus (GEO; accession number GSE69400) (106, 107) and processed to generate nucleosome maps as described previously (98).

Gene Ontology (GO) enrichment analysis was performed using the PANTHER Overrepresentation Test (Fisher’s exact test with FDR correction), separately for genes with increased or decreased Rpb1 occupancy, using the 5,764 analyzed genes as the reference set.

### As-biotin pull-down from native yeast extracts

Logarithmically growing yeast cells were harvested and resuspended in 500 µl of TNTM buffer (50 mM Tris-HCl, pH 7.5, 300 mM NaCl, 1% Triton X-100, 2.5 mM MgCl₂) supplemented with protease inhibitor cocktail [Sigma-Aldrich, P8215] and 1 mM PMSF. Cells were disrupted using glass beads with intermittent cooling on ice, and the lysates were clarified by centrifugation. The resulting extracts were treated with 1 KU/ml Benzonase (Sigma-Aldrich, E1014) and 20 µl RNase A (Thermo Scientific, EN0531) for 30 min at 4°C. After collecting 10 µl as input, the remaining extract was divided into four aliquots and incubated for 15 min with rotation under the following conditions: no additions, 100 µM As-biotin, 100 µM As(III), or both compounds (100 µM each). Subsequently, 15 µl of streptavidin beads, prewashed with TNTM buffer, were added to each sample and incubated for 15 min at room temperature. The beads were washed with TNTM buffer, resuspended in Laemmli sample buffer, and analyzed by immunoblotting.

### As-biotin pull-down from native mammalian cell extracts

A549 cells were cultured as described above for the isolation of whole-cell protein extracts from mammalian cells. Cells were scraped directly into ice-cold TNTM buffer (50 mM Tris-HCl, pH 7.5, 300 mM NaCl, 1% Triton X-100, 2.5 mM MgCl₂) supplemented with protease inhibitor cocktail (Thermo Fisher, 78441) and 1 mM PMSF and incubated for 30 min at 4°C with rotation. Lysates were clarified by centrifugation and treated with 1 kU/ml benzonase (Sigma-Aldrich, E1014) and 20 µl RNase A (Thermo Scientific, EN0531) for 30 min at 4°C. After collecting 10 µl as input, the remaining lysate was divided into four aliquots and incubated for 15 min with rotation under the following conditions: no additions, 100 µM As-biotin, 100 µM As(III), or both compounds (100 µM each). Subsequently, 15 µl of prewashed streptavidin beads were added to each sample and incubated for 15 min at room temperature. The beads were washed with TNTM buffer, resuspended in Laemmli sample buffer, boiled, and analyzed by immunoblotting.

### RNAPII complex purification for As-biotin experiments

Approximately 10¹¹ logarithmically growing untagged or Rpb1-HTP (His₆–TEV cleavage site–2× protein A) tagged cells were harvested by centrifugation, extensively washed with water and YEB buffer (25 mM HEPES–KOH pH 7.9, 100 mM KCl, 2 mM MgCl₂, 5% glycerol, 0.1% Triton X-100, 1 mM DTT, protease inhibitors [Sigma-Aldrich, P8215], 1 mM PMSF), and finally resuspended in 4 ml YEB. Cells were lysed in a bead beater at 4°C, and whole-cell extracts were centrifuged for 15 min at 15,000 rpm. The resulting protein extracts were incubated with 2 kU/ml benzonase (Sigma-Aldrich, E1014) and 100 µg/ml RNase A (Thermo Scientific, EN0531) for 1 h at 4°C. The extracts were then incubated with IgG Sepharose beads (Cytiva, 17096901) for 3 h at 4°C with rotation. Next, the beads were washed several times with IPP buffer (10 mM Tris pH 7.9, 100 mM NaCl, 0.1% Triton X-100) and then with TEV buffer (50 mM Tris pH 7.9, 0.2 mM EDTA, 100 mM NaCl, 0.1% Triton X-100, 1 mM DTT). To release Rpb1, the IgG Sepharose beads were incubated overnight at 4°C in TEV buffer containing 400 U/ml TEV protease (Sigma-Aldrich, SAE0118). The next day, the beads were separated from the buffer, and the resulting eluate was incubated with 200 µl Ni-NTA resin (Qiagen, 30210) for 1 h at 4°C with rotation. The beads were washed three times with Ni buffer (20 mM sodium phosphate, 300 mM NaCl, 20 mM imidazole, pH 7.4), followed by protein elution by 5 min incubation at room temperature in Ni buffer containing 500 mM imidazole. The resulting eluate was loaded onto Zeba™ Spin Desalting Columns (Thermo Scientific, 89890). The purified RNAPII complex was suspended in IP buffer (20 mM sodium phosphate, 300 mM NaCl, pH 7.4). The RNAPII-containing solution was then divided into two aliquots. One aliquot was supplemented with 25 µM biotin-conjugated arsenic (As-biotin; TRC, TRC-B394970), whereas the other was left untreated. As-biotin was also added to the sample prepared from the untagged strain. All samples were incubated for 30 min at room temperature.

In parallel, 30 µl streptavidin magnetic beads were blocked for 20 min at 4°C in IP buffer containing 0.2% BSA, then extensively washed with IP buffer. After the 30-min incubation, 10 µl of the blocked streptavidin beads were added to each sample and incubated for 15 min at 4°C with occasional mixing. Lysates were then removed, the beads were washed twice with IP buffer and twice with IP buffer containing 0.25% NP-40 and 500 mM NaCl, washed again with IP buffer, and finally resuspended in IP buffer.

### RNAPII complex purification for RNA elongation assay

Yeast cells expressing FLAG-tagged Rpb1 were grown in 2 L of YPD medium to an OD600 of approximately 1.5, harvested by centrifugation, washed with ice-cold water, and frozen at −80°C. Cells were resuspended in 6 mL of lysis buffer (200 mM KCl, 20 mM Tris-HCl, pH 8.0, 5 mM magnesium acetate, 20% glycerol, and 1× cOmplete protease inhibitor cocktail; Roche, 11697498001) and disrupted with zirconia beads in 2-mL tubes using five 8-min cycles, with cooling on ice between cycles. Lysates were clarified by centrifugation at 16,000 × g for 30 min and supplemented with 0.5% IGEPAL CA-630. The clarified extracts were incubated overnight with pre-equilibrated anti-FLAG magnetic agarose beads (Pierce Anti-DYKDDDDK Magnetic Agarose; Thermo Fisher Scientific, A36797) at a ratio of 10–15 µL beads per 1 mL extract, with continuous rotation. The beads were washed five times with lysis buffer, followed by six washes with wash buffer (200 mM potassium acetate, 20 mM HEPES, pH 7.8, 1 mM MgCl₂, and 20% glycerol). RNAPII was released by incubation with 12 µL HaloTEV protease (Promega, G6602) in wash buffer for 1 h at 25°C with shaking at 900 rpm. The bead-free eluate was flash-frozen in liquid nitrogen and stored at −80°C. The RNAPII preparation was characterized by mass photometry, which revealed a population with an apparent molecular mass of approximately 520–530 kDa, consistent with the expected molecular mass of the tagged yeast RNAPII complex (∼514 kDa) and the presence of residual TEV protease.

### *In vitro* RNAPII transcription elongation assay

RNA–DNA–DNA scaffolds were assembled by mixing an RNA primer, a complementary DNA template strand, and a non-template DNA strand (10 pmol each) in 15 µL of ultrapure water (A&A Biotechnology, 005-500). The mixture was heated to 95°C for 5 min and allowed to cool to room temperature. The sequences of the RNA and DNA oligonucleotides used for scaffold assembly are provided in SI Appendix, Table S2. For each reaction, 2 pmol of the preassembled RNA–DNA–DNA scaffold was incubated with a constant volume of purified RNAPII preparation for 10 min at room temperature. Transcription was initiated by adding an equal volume of transcription reaction mixture. Elongation reactions were performed in transcription buffer containing 60 mM (NH₄)₂SO₄, 20 mM HEPES–KOH (pH 7.6), 8 mM MgSO₄, 10 µM ZnCl₂, and 10% glycerol. Reactions contained 1 mM NTPs and the indicated concentrations of As(III). Samples were incubated at 28°C for 10 min. Following incubation, each reaction was adjusted to 250 µL with ultrapure water and mixed with 600 µL TRIzol reagent (Invitrogen, 15596026). After incubation for 5 min at room temperature, 170 µL chloroform was added, followed by an additional 3-min incubation. Samples were centrifuged at 12,000 × g for 15 min at 4°C, and the aqueous phase was transferred to a fresh tube. RNA was precipitated by adding 430 µL isopropanol and 1 µL GlycoBlue Coprecipitant (Invitrogen, AM9516), followed by incubation at −20°C for 30 min and centrifugation at 20,000 × g for 20 min at 4°C. RNA pellets were washed with 860 µL of 75% ethanol, centrifuged at 15,000 × g for 15 min at 4°C, air-dried for 10 min, and resuspended in 10 µL ultrapure water. Purified RNA was mixed with 2× RNA Loading Dye (Thermo Scientific, R0641), heat-denatured at 80°C for 5 min, and separated on 20% denaturing polyacrylamide gels containing 8 M urea. Fluorescently labeled RNA products were visualized using a Typhoon FLA 9500 fluorescence scanner.

### Bottom-up LC–MS analysis

Sodium deoxycholate was added to bead samples to a final concentration of 1%, followed by incubation at 65 °C for 10 min. Trypsin (300 ng) was then added, and on-bead digestion was carried out overnight at 37 °C. The next day, the digest was separated from the beads, acidified, and sodium deoxycholate was removed by centrifugation. The supernatant was desalted using STAGE tips (108). The resulting peptide pellet was resuspended in 0.1% formic acid (FA) and 3% acetonitrile (ACN). LC–MS analyses were performed on an M-Class ACQUITY UPLC system coupled to a Synapt XS HDMS mass spectrometer equipped with a nanoESI source. Mobile phase A was H₂O containing 0.1% FA, and mobile phase B was ACN containing 0.1% FA. Peptides were separated on a C18 BEH analytical column (75 µm × 250 mm) maintained at 65 °C using an 8–40% B linear gradient over 24 min at a flow rate of 300 nL/min. A 3-minute trapping step was performed prior to analytical separation. Data were acquired in positive-ion mode (ESI+) using data-dependent acquisition (DDA). MS and MS/MS scan times were 0.2 s, with m/z ranges of 250–1250 (MS) and 100–1800 (MS/MS). Analyser mode was set to Resolution. For each MS survey scan, the six most intense precursor ions with charge states 2+, 3+, or 4+ were selected for MS/MS, with one scan permitted per transition. Collision energy was applied as an m/z-dependent ramp in the trap cell (start LM/HM: 15–40 V; end LM/HM: 18–50 V). Source conditions were fine-tuned. (Glu¹)-Fibrinopeptide B was acquired in parallel as a lock-mass reference, and lock-mass correction was applied post-acquisition. Raw data were processed in Progenesis QI for Proteomics (Nonlinear Dynamics). Spectra were first searched against the PRIDE Contaminants Library using the following parameters: precursor tolerance, 7.5 ppm; fragment tolerance, 25 ppm; minimum fragments per peptide, 10; and shared hits across charge states enabled. An .mgf file was then exported (excluding queries assigned to contaminants and unrelated to yeast proteins) and searched with Mascot Server v2.8.0.1 (Matrix Science) against the *Saccharomyces cerevisiae* protein sequence database (UP000000757), with the streptavidin sequence appended (UniProt entry). Mascot search parameters were: peptide mass tolerance, 7.5 ppm; fragment mass tolerance, 25 ppm; maximum protein mass, 1 MDa; enzyme, trypsin; maximum missed cleavages, 2; variable modifications, methionine oxidation and protein N-terminal acetylation; and PSM-level FDR, 1%. Protein hits were grouped into families. The Progenesis protein output was exported, and spectral counts for RNAPII complex subunits were compared across samples.

## Data availability

All data needed to evaluate the conclusions in the paper are present in the article and/or its Supporting Information. The next-generation sequencing data (unprocessed read files and processed coverage files) can be accessed at GEO (https://www.ncbi.nlm.nih.gov/geo/) with accession number GSE318138. Raw LC–MS data have been deposited in the Zenodo Research Repository (DOI: https://doi.org/10.5281/zenodo.18412788).

## Acknowledgments

We thank Andrés Aguilera, Camilla Bjorkegren, Dana Branzei, Frédéric Beckouët, Jennifer Cobb, Elisa Dultz, Brian Luke, Adele Marston, Felix Prado, Rodney Rothstein, Barbara Dirac Svejstrup, Lorraine Symington, José Antonio Tercero, and Helle Ulrich for providing yeast strains and plasmids. We thank Dorota Dziadkowiec for insightful discussions during the course of this project and for critically reading the manuscript. This study was supported by a National Science Centre (Poland) grant (2022/47/B/NZ2/00355) to I.L.

## Author contributions

I.L. designed research; A.N., E.M.-D., K.K., M.T., A.D., D.P.B., A.P.K., I.L. performed research; A.N., P.T., K.K., M.T., A.D., D.P.B., T.W.T., A.P.K., I.L. analyzed data; and I.L. wrote the paper.

The authors declare no competing interest.

