## Supplementary information for "RNA polymerase II links arsenic exposure to chromosome-wide γH2A signaling"

**This PDF file includes:**
Tables S1 to S2
Figure legends for Figs. S1–S13
Figures S1 to S13
Unprocessed, uncropped immunoblot and gel images (Source Data)

**Other supporting materials for this manuscript include the following:**
Datasets S1 to S2

**Table S1. Yeast strains, cell lines and plasmids used in this study.**

| Yeast strains | | |
|---|---|---|
| **Genotype** | **Source** | **Strain** |
| *Saccharomyces cerevisiae* W303*: MATa leu2-3,112 trp1-1 can1-100 ura3-1 ade2-1 his3-11,15* | R. Rothstein | W303-1a |
| *Saccharomyces cerevisiae* W303*: MATa hta1-S129A::HIS3 hta2-S129A::TRP1 SCC1-9PK::TRP1 URA3::GPD-TK7* | J.Cobb | JC-1597 |
| *Saccharomyces cerevisiae* W303*: MATa yku70Δ::kanMX6* | Lab collection | IL006 |
| *Saccharomyces cerevisiae* W303: *MATa ADE2 bar1::LEU2 RAD52-YFP* | R. Rothstein | W3749-14C |
| *Saccharomyces cerevisiae* W303*: MATa Rad59-13cMyc-KanMX6* | This work | AN001 |
| *Saccharomyces cerevisiae* W303: *MATa exo1Δ::kanMX6* | Lab collection | AN003 |
| *Saccharomyces cerevisiae* BY474*: MATa his3Δ1 leu2Δ0 ura3Δ0 met15Δ0 RAD52-MN::HIS3MX6* | F. Prado | BYR52MN |
| *Saccharomyces cerevisiae* W303: *MATa ADE2 bar1::LEU2 RFA1-YFP* | R. Rothstein | W3775-12C |
| *Saccharomyces cerevisiae* W303: *MATa sgs1Δ::kanMX6* | Lab collection | MC012 |
| *Saccharomyces cerevisiae* W303: *MATa exo1Δ::kanMX6 sgs1Δ:: LEU2* | This work | AN0010 |
| *Saccharomyces cerevisiae* W303*: MATα sgs1-D664Δ ADE2* | R. Rothstein | W5927-20A |
| *Saccharomyces cerevisiae* W303: *MATa mre11Δ::LEU2* | D. Branzei | FY1643 |
| *Saccharomyces cerevisiae* W303: *MATa mre11-H125N-NatMX* | L. Symington | LSY3366 |
| *Saccharomyces cerevisiae* W303: *MATa ura::ADH1 promoter-OsTIR1–9myc::URA3, Scc1-Pk3-aid::KanMX4* | F. Beckouët | K18154 |
| *Saccharomyces cerevisiae* W303*: MATa rho⁰* | Lab collection | AN012 |
| *Saccharomyces cerevisiae* W303*: MATa yno1Δ::kanMX6* | Lab collection | AN013 |
| *Saccharomyces cerevisiae* W303*: MATa rpb1-1* | C. Björkegren | CB3295 |
| *Saccharomyces cerevisiae* W303*: MATα rpb1-1* | C. Björkegren | CB3291 |
| *Saccharomyces cerevisiae* W303: *MATα RFA1-mCherry::hphNT1* | C. J. A. Tercero | YCL48 |
| *Saccharomyces cerevisiae* DF5: *RFA1-GFP::HISMX3, TRP1::AFB2, RPO21-AID*-9cMyc::natNT2* | H. Ulrich | 5578 |
| *Saccharomyces cerevisiae* W303: *RPB2::3V5-IAA7-KANMX*, *HIS3::GPD1-OsTIR* | E. Dultz | KWY8449 |
| *Saccharomyces cerevisiae* BY4741: *Rpo21-HTP::URA* | B. D. Svejstrup | YLM594 |
| *Saccharomyces cerevisiae* W303: *rpb1S751F* | A. Aguilera | WSR8-5A |
| *Saccharomyces cerevisiae* W303: *rpb9 ::HIS3* | A. Aguilera | WRP9-3C |

| | | |
|---|---|---|
| *Saccharomyces cerevisiae BY4741: MATa his3Δ1 leu2Δ0 met15Δ0 ura3Δ0 rpb1::Rpb1-mCherry-Avi-TEV-His-FLAG* | T.Turowski | TT1 |
| *Schizosaccharomyces pombe: rad21::rad21-6HA:KanMX6 ade6-210 leu1-32 ura4-D18 his3-D1 arg3-D4* | A.Marston | AM635 |
| **Cell lines** | | |
| **Cell line** | **Source** | **Identifier** |
| A549 | Department of Molecular Physiology and Neurobiology, University of Wrocław | RRID: CVCL_0023 |
| **Plasmids** | | |
| Plasmid *pRS426 pGPD, 2μ URA3* | B. Luke | pBL189 |
| Plasmid *pRS426 pGPD-RNH1-HA, 2μ URA3* | B. Luke | pBB39 |

**Table S2. RNA and DNA oligonucleotides used in the RNAPII elongation assay.**

| Content | Sequence |
|---|---|
| RNA Primer (FAM labelled) | FAM-CUAUACUCCUUUGAUGC |
| Template DNA | CATCTGGAACAAAAAAAAAATCTCTTAAAATCTGCTGAGGATGGTTTCG ATCCATCGACGAGACGCATCAAAGGAGAGTGAGATCG |
| Non-Template DNA | CGATCTCACTCTCCTTTGATGCGTCTCCAACATCTCTCACCCATCTCCA CAGCAGATTTTAAGAGATTTTTTTTTTGTTCCAGATGGACAGCGAGACT AGAGTGAC |

**Figure Legends**

**SI Appendix, Fig. S1. Acute As(III) exposure elicits a specific, dose-dependent, and transient γH2A response.** (A) Clonogenic survival of WT cells exposed to the indicated concentrations of As(III) for 1 or 2 h. Data are expressed as mean ± SD of three biological replicates relative to untreated cultures. (B) Intracellular As concentration in WT cells treated with 0.5 mM As(III) for 1 or 2 h, measured by atomic absorption spectrometry. Data represent mean ± SD from three independent experiments, each performed in triplicate. Arsenic was not detected in untreated cells. (C) γH2A immunoblot after 1 h exposure to As(III), Fe, Cu, Co, Ni, or Cd; all metals were used at 0.5 mM except Cd, which was used at 50 μM. (D) Dose-dependent γH2A induction in WT cells treated with the indicated concentrations of As(III) for 1 h. (E) Early time course of γH2A formation after addition of 0.5 mM As(III). (F) Extended time course of γH2A formation after exposure to 0.5 mM As(III) for the indicated times. (G) Serial-dilution growth assays of WT and H2A-S129A cells on control plates or plates containing the indicated concentrations of As(III) or phleomycin (Phl). Representative results from at least three independent experiments are shown. Pgk1 and Ponceau S staining served as loading and total-protein controls, respectively, in the immunoblots.

**SI Appendix, Fig. S2. Reproducibility and specificity of γH2A ChIP–seq profiling.** (A) Spike-in-normalized γH2A ChIP–seq profiles of chromosome III from three independent biological replicates of logarithmically growing WT cells left untreated or exposed to 0.5 mM As(III) for 1 h. Replicates 1 and 2 were performed using an anti-γH2A antibody from Abcam, whereas replicate 3 was performed using an anti-γH2A antibody from Active Motif. (B) Pairwise Spearman correlations of genome-wide γH2A ChIP–seq signals across all untreated and As(III)-treated biological replicates. (C) γH2A ChIP–seq profiles of chromosomes III and IV in H2A-S129A cells left untreated or exposed to 0.5 mM As(III) for 1 h. Profiles from a single biological experiment are shown. (D) γH2A ChIP–qPCR measurements at selected ORF, ARS, centromeric, pericentromeric, telomere-proximal, and rDNA loci in WT cells left untreated or exposed to 0.5 mM As(III) for 1 h. Data are shown as mean ± SD of three independent biological replicates. Statistical significance was assessed using a two-tailed t test.

**SI Appendix, Fig. S3.** As(III) does not produce prominent RPA-enriched domains. (A) RPA ChIP–seq profiles along chromosome III from two independent biological replicates of logarithmically growing WT cells left untreated or exposed to 0.5 mM As(III) for 1 h. (B) RPA metaprofiles across scaled ORFs and 0.5-kb flanking regions under the same conditions. (C) Representative DNA-content profiles measured by flow cytometry in logarithmically growing WT cells left untreated or after 1 or 2 h of exposure to 0.5 mM As(III).

**SI Appendix, Fig. S4. As(III) induces modest, broadly distributed changes in HR-factor occupancy.** (A, C and E) ChIP–seq profiles of Rad52, Rad51, and Rad59 along chromosome III in logarithmically growing WT cells left untreated or exposed to 0.5 mM As(III) for 2 h. Two biological replicates are shown. (B, D and F) Representative metaprofiles of Rad52, Rad51, and Rad59 occupancy across ORFs under the same conditions.

**SI Appendix, Fig. S5. Dose-dependent differences in the kinetics and genomic γH2A response to phleomycin.** (A) γH2A immunoblot of logarithmically growing WT cells exposed to 50 μg/ml phleomycin (Phl) for the indicated times. Pgk1 and Ponceau S staining served as loading and total-protein controls, respectively, in the immunoblots. (B) γH2A immunoblot of logarithmically growing WT cells exposed to 2.5 μg/ml Phl for the indicated times. (C) Spike-in-normalized genome-wide γH2A ChIP–seq profiles along chromosome IV in WT cells treated with 0.5 mM As(III), 2.5 or 50 μg/ml Phl for 1 h. Data are from a single biological experiment.

**SI Appendix, Fig. S6. Oxidative stress and cohesin-mediated chromatin organization are not major drivers of As(III)-induced γH2A.** (A) Flow-cytometric analysis of ROS-sensitive dye fluorescence (MitoSOX, CellROX, DCF, and DHE) in WT cells left untreated or exposed to 0.5 mM As(III) for 1 h. (B and C) γH2A immunoblotting in cells lacking mitochondrial respiration (*rho0*) (B) or the NADPH oxidase *YNO1* (C), left untreated or exposed to 0.5 mM As(III) for 1 h. Pgk1 and Ponceau S staining served as loading and total-protein controls, respectively, in the immunoblots. (D) γH2A immunoblotting in Scc1–AID–3Pk cells with or without 0.5 mM auxin pretreatment for 1 h, and in untagged control cells, followed by exposure to 0.5 mM As(III) for 1 h. Scc1–AID–3Pk depletion was monitored by anti-Pk immunoblotting. (E) Quantification of the Rad52–MN ChEC data shown in Fig. 2G.

**SI Appendix, Fig. S7. As(III)-induced γH2A signaling in G2/M- and G1-arrested cells.** (A) Spike-in-normalized γH2A ChIP–seq profiles along chromosome III from two independent biological replicates of G2/M-arrested WT cells left untreated or exposed to 0.5 mM As(III) for 1 h. (B) DNA-content profiles of logarithmically growing WT cells and WT cells arrested in G1 with α-factor before and after 1 h exposure to 0.5 mM As(III). (C) Spike-in-normalized γH2A ChIP–seq profiles along chromosomes III and IV in G1-arrested WT cells left untreated or exposed to 0.5 mM As(III) for 1 h. Data are from a single biological experiment. (D) DNA-content profiles of logarithmically growing *ku70*Δ cells and *ku70*Δ cells arrested in G1 before and after 1 h exposure to 0.5 mM As(III). (E) Spike-in-normalized γH2A ChIP–seq profiles along chromosomes III and IV in G1-arrested *ku70*Δ cells left untreated or exposed to 0.5 mM As(III) for 1 h. Data are from a single biological experiment.

**SI Appendix, Fig. S8. As(III)-induced checkpoint signaling persists in resection-defective cells.** (A and B) Immunoblot analysis of γH2A and Rad53 phosphorylation in WT and *sgs1*Δ *exo1*Δ cells (A), or WT and *mre11*Δ cells (B), left untreated or exposed to 0.5 mM As(III) for 1 h. Pgk1 and Ponceau S staining served as loading and total-protein controls, respectively. (C) Serial-dilution growth assays of WT, *exo1*Δ, *sgs1*Δ, *sgs1*Δ *exo1*Δ, *sgs1*-D664, *mre11*Δ, and *mre11*-H125N cells on control plates or plates containing 0.5 mM As(III).

**SI Appendix, Fig. S9. Effects of Rpb1 and Rpb2 depletion on As(III)-induced γH2A signaling.** (A) Representative DNA-content profiles of Rpb1-AID cells left untreated or subjected to the indicated auxin treatments, with or without subsequent exposure to 0.5 mM As(III) for 1 h. (B) Immunoblot analysis of Rpb2–3V5–AID, γH2A, and total H2A in cells subjected to the indicated auxin treatments, with or without exposure to 0.5 mM As(III) for 1 h. Rpb2–3V5–AID depletion was monitored by anti-Pk immunoblotting. Pgk1 and Ponceau S staining served as loading and total-protein controls, respectively.

**SI Appendix, Fig. S10. Effects of different transcriptional perturbations on γH2A signaling.** (A and B) γH2A immunoblot analysis of WT cells treated with 3 µg/ml (A) or 20 µg/ml (B) thiolutin (Thio), with or without 0.5 mM As(III). Cells were pretreated with thiolutin for 1 h and subsequently exposed or not to As(III) for an additional 1 h. (C and D) γH2A immunoblot analysis of WT, *rpb1*-S751F, and *rpb9*Δ cells at 30°C (C) or after a shift to 37°C for 1 h (D), with or without subsequent exposure to 0.5 mM As(III) for 1 h. Pgk1 and Ponceau S staining served as loading and total-protein controls, respectively.

**SI Appendix, Fig. S11. Additional analyses of γH2A responses to RNAPII perturbation.** (A) Spike-in-normalized γH2A ChIP–seq profiles along chromosome III in *rpb1-1* cells maintained at 25°C or shifted to 37°C. Two independent biological replicates are shown. (B) γH2A ChIP–seq metaprofiles across scaled ORFs in *rpb1-1* cells at 25°C or after a shift to 37°C. (C and D) γH2A immunoblot analysis of WT (C) and *rpb1-1* (D) cells shifted to 37°C and then left untreated or exposed to 0.25 mM As(III). (E and F) γH2A immunoblot analysis of WT cells treated for 1 h with 0.01% MMS (E) or 50 µg/ml Phl (F), with or without 0.5 mM As(III). Pgk1 and Ponceau S staining served as loading and total-protein controls, respectively, in the immunoblotting experiments. (G) PFGE analysis of chromosomal DNA from *rpb1-1* cells maintained at 25°C or shifted to 37°C. Cells treated with 50 µg/ml Phl for 1 h at 37°C served as a positive control for chromosome fragmentation. (H) Representative fluorescence microscopy images of Rfa1–mCherry and Rad52–YFP foci in WT and *rpb1-1* cells. Corresponding quantifications are shown in Fig. 3I and J. (I) γH2A immunoblot analysis of cells with or without *RNH1* overexpression, left untreated or exposed to As(III). Rnh1–HA expression was detected by anti-HA immunoblotting.

**SI Appendix, Fig. S12. Changes in RNAPII subunit abundance following As(III) exposure and Rpb2 depletion.** (A–C) Immunoblot analysis of Rpb1 abundance in logarithmically growing WT cells exposed to 0.5 mM As(III) (A), 50 µg/ml phleomycin (Phl) (B), or 2.5 µg/ml Phl (C) for the indicated times. (D and E) Immunoblot analysis of Ser2-phosphorylated (D) and Ser5-phosphorylated (E) Rpb1 in WT cells exposed to 0.5 mM As(III) for the indicated times. (F) Immunoblot analysis of Rpb1, Rpb2–3V5–AID, Rpb3, and γH2A in cells left untreated or exposed to 0.5 mM As(III) for 1 or 2 h. (G) Immunoblot analysis of Rpb1, Rpb2–3V5–AID, Rpb3, and γH2A in cells left untreated or exposed to auxin for 1 or 2 h. Rpb2–3V5–AID was detected using anti-Pk antibodies. Pgk1 and Ponceau S staining served as loading and total-protein controls, respectively, in the immunoblots.

**SI Appendix, Fig. S13. Genomic and biochemical characterization of the relationship between As(III) and RNAPII.** (A) Spike-in-normalized Rpb1 ChIP–seq profiles along chromosome III from two independent biological replicates of logarithmically growing WT cells left untreated or exposed to 0.5 mM As(III) for 1 h. (B) SDS–PAGE analysis of endogenous RNAPII purified from yeast cells expressing Rpb1–HTP, with an untagged strain serving as a negative purification control. The positions of RNAPII subunits are indicated. Asterisks indicate unidentified co-purifying proteins. (C) Mass spectrometry analysis of As-biotin pull-downs performed with purified yeast RNAPII. Heatmap showing spectral counts for individual RNAPII subunits (Rpb1–Rpb12) in the As-biotin, beads-only, and untagged control samples. (D) Quantification of full-length RNA products generated in an *in vitro* transcription elongation assay using purified yeast RNAPII in the absence or presence of the indicated As(III) concentrations (0.001–1 mM). Full-length RNA products are expressed as a percentage of the total RNA signal. Data are presented as mean ± SD from three independent experiments.

Fig. S1

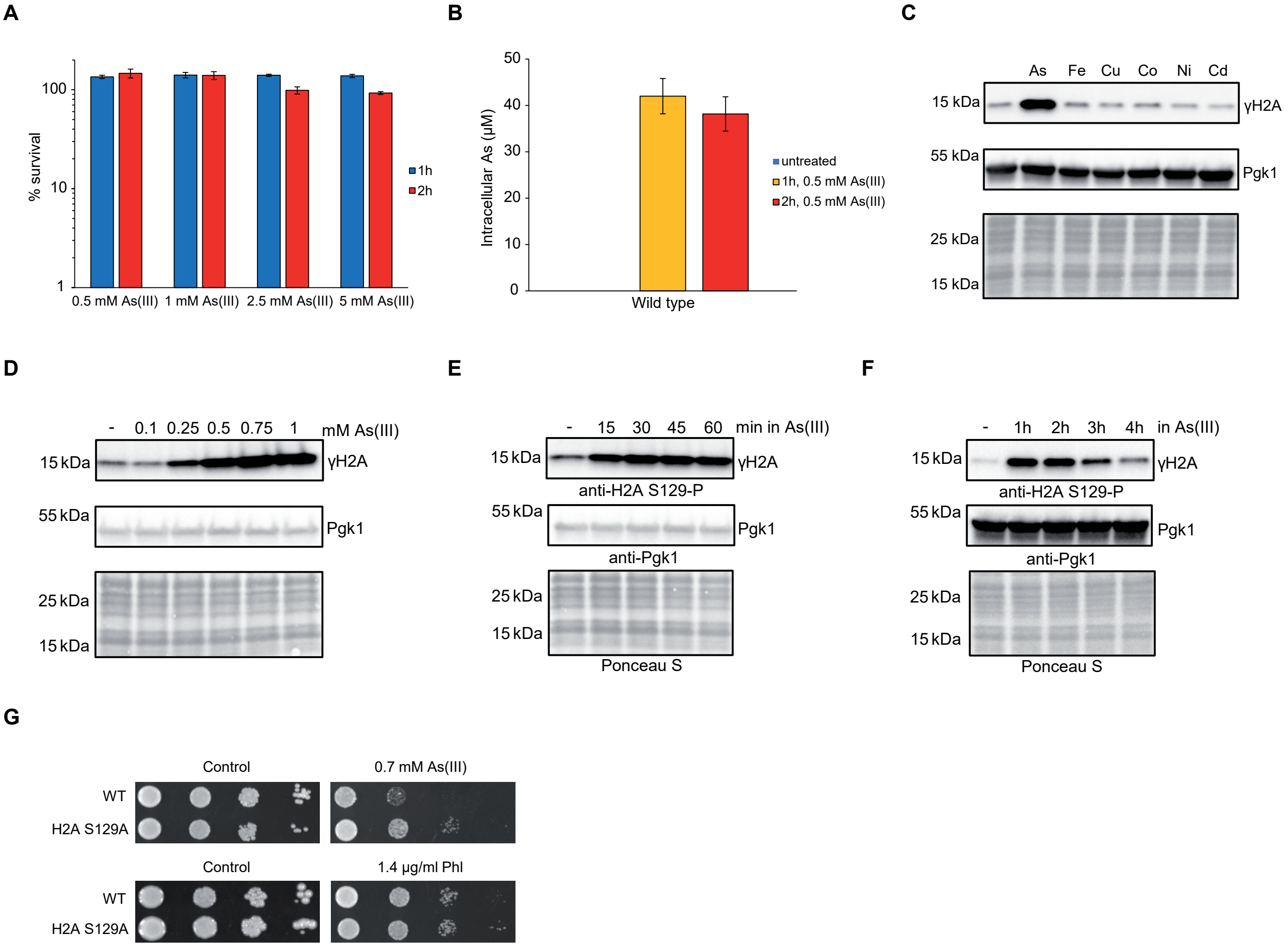
A
100
10
1
% survival
1h
2h
0.5 mM As(III)
1 mM As(III)
2.5 mM As(III)
5 mM As(III)
B
50
40
30
20
10
0
Intracellular As (µM)
untreated
1h, 0.5 mM As(III)
2h, 0.5 mM As(III)
Wild type
C
As
Fe
Cu
Co
Ni
Cd
15 kDa
γH2A
55 kDa
Pgk1
25 kDa
15 kDa
D
-
0.1
0.25
0.5
0.75
1
mM As(III)
15 kDa
γH2A
55 kDa
Pgk1
25 kDa
15 kDa
E
-
15
30
45
60
min in As(III)
15 kDa
γH2A
anti-H2A S129-P
55 kDa
Pgk1
anti-Pgk1
25 kDa
15 kDa
Ponceau S
F
-
1h
2h
3h
4h
in As(III)
15 kDa
γH2A
anti-H2A S129-P
55 kDa
Pgk1
anti-Pgk1
25 kDa
15 kDa
Ponceau S
G
Control
0.7 mM As(III)
WT
H2A S129A
Control
1.4 µg/ml Phl
WT
H2A S129A

**Fig. S2**

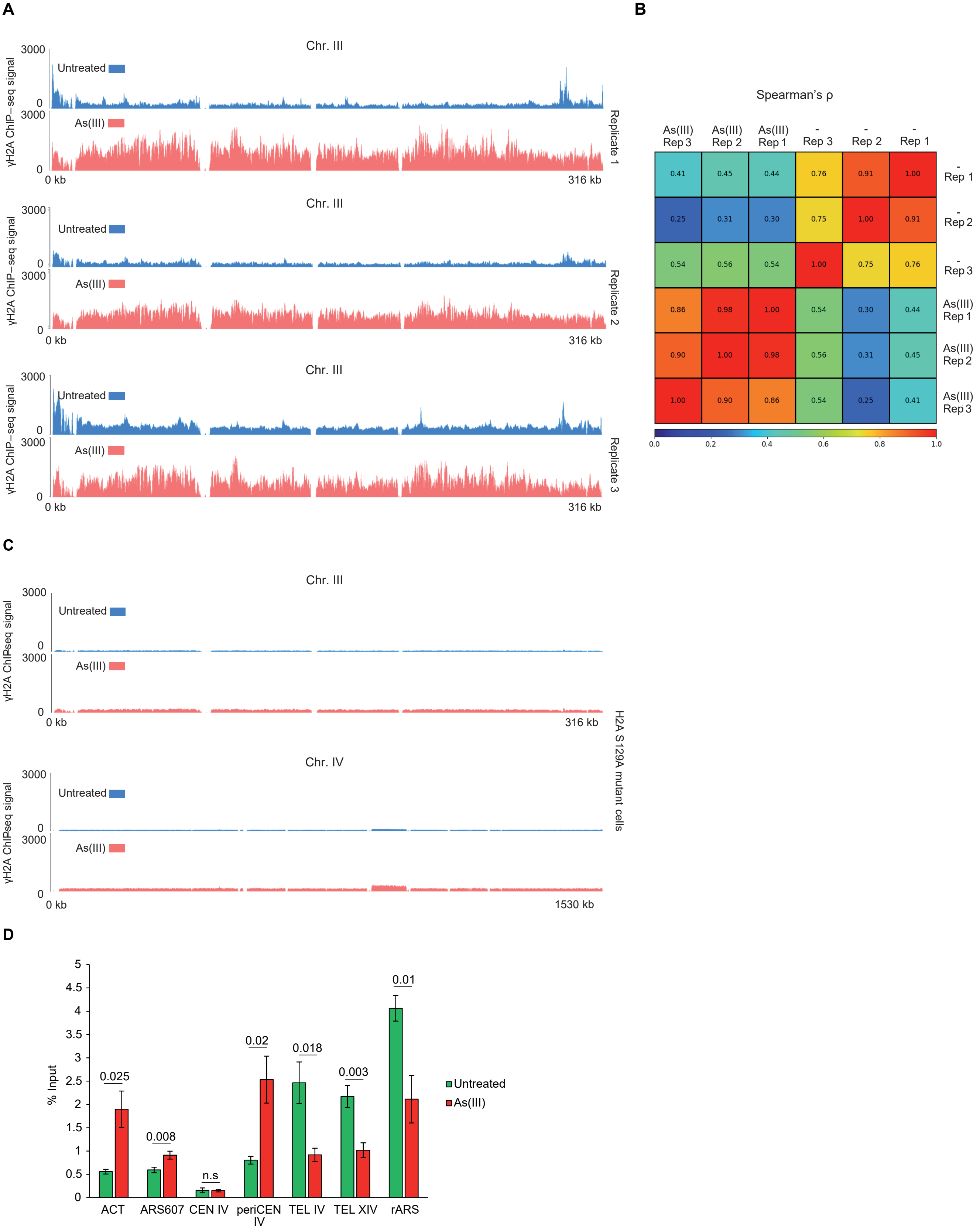
A
Chr. III
γH2A ChIP−seq signal
3000
0
Untreated
As(III)
0 kb
316 kb
Replicate 1
Replicate 2
Replicate 3
B
Spearman's ρ
As(III) Rep 3
As(III) Rep 2
As(III) Rep 1
- Rep 3
- Rep 2
- Rep 1
0.41 0.45 0.44 0.76 0.91 1.00
0.25 0.31 0.30 0.75 1.00 0.91
0.54 0.56 0.54 1.00 0.75 0.76
0.86 0.98 1.00 0.54 0.30 0.44
0.90 1.00 0.98 0.56 0.31 0.45
1.00 0.90 0.86 0.54 0.25 0.41
0.0 0.2 0.4 0.6 0.8 1.0
C
Chr. III
Chr. IV
γH2A ChIPseq signal
1530 kb
H2A S129A mutant cells
D
% Input
0 0.5 1 1.5 2 2.5 3 3.5 4 4.5 5
0.025
0.008
n.s
0.02
0.018
0.003
0.01
ACT
ARS607
CEN IV
periCEN IV
TEL IV
TEL XIV
rARS
Untreated
As(III)

Fig. S3

A

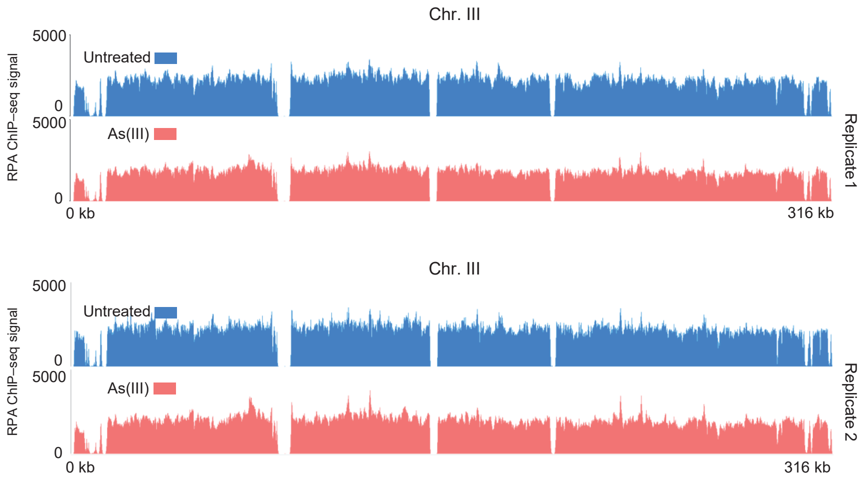
Chr. III
5000
Untreated
0
5000
As(III)
0
RPA ChIP–seq signal
0 kb
316 kb
Replicate 1
Chr. III
5000
Untreated
0
5000
As(III)
0
RPA ChIP–seq signal
0 kb
316 kb
Replicate 2

B

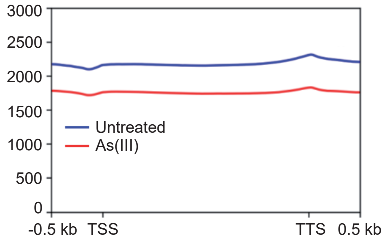
3000
2500
2000
1500
1000
500
0
Untreated
As(III)
-0.5 kb
TSS
TTS
0.5 kb

C

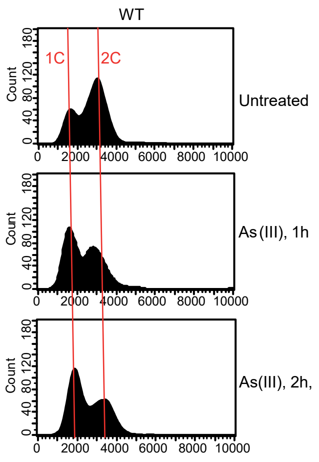
WT
1C
2C
Count
Untreated
As(III), 1h
As(III), 2h,
0
2000
4000
6000
8000
10000

Fig. S4

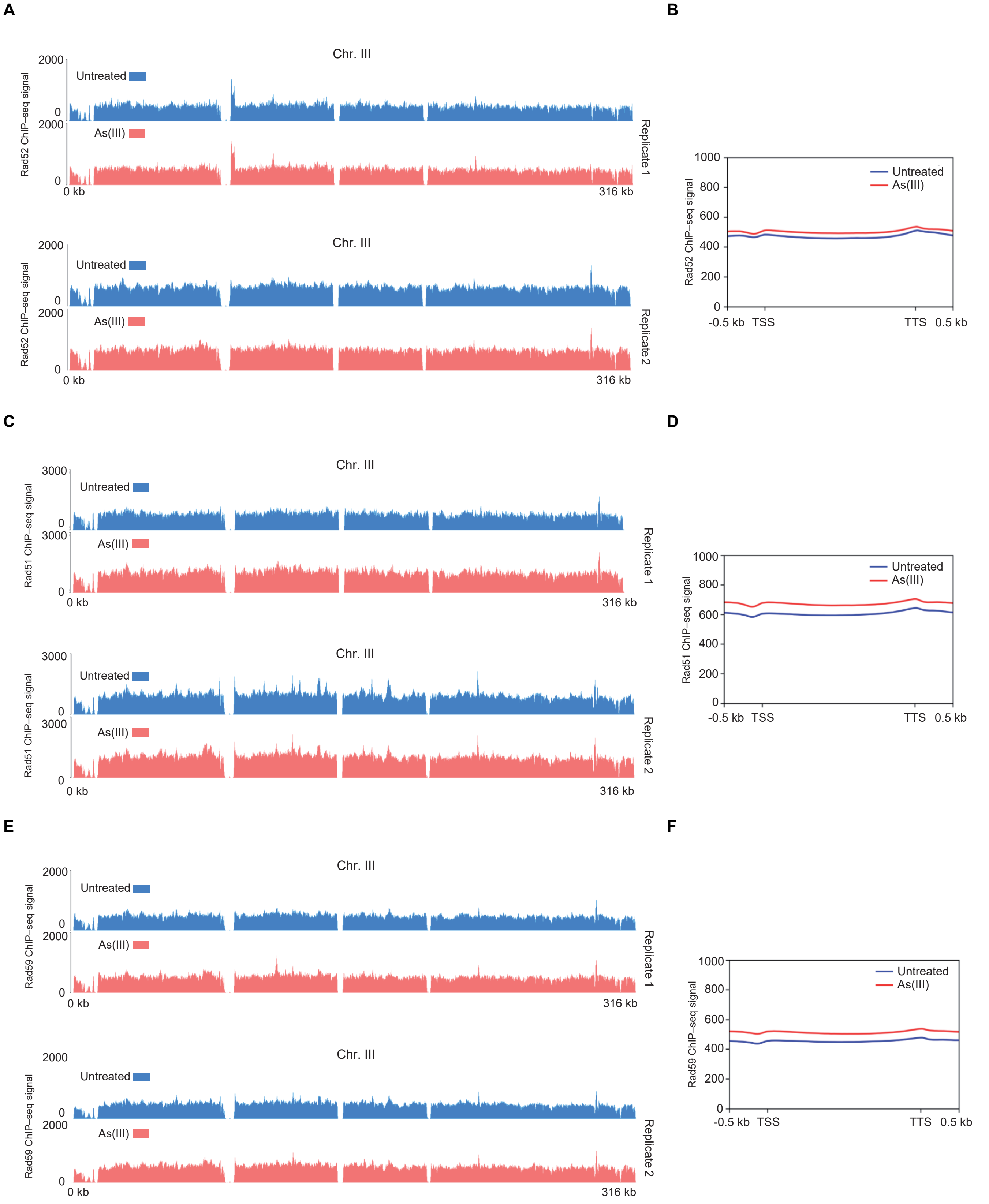
A
Chr. III
Rad52 ChIP–seq signal
Untreated
As(III)
2000
0
0 kb
316 kb
Replicate 1
Replicate 2
B
Rad52 ChIP–seq signal
1000
800
600
400
200
0
Untreated
As(III)
-0.5 kb
TSS
TTS
0.5 kb
C
Chr. III
Rad51 ChIP–seq signal
3000
0
Untreated
As(III)
0 kb
316 kb
Replicate 1
Replicate 2
D
Rad51 ChIP–seq signal
1000
800
600
400
200
0
Untreated
As(III)
-0.5 kb
TSS
TTS
0.5 kb
E
Chr. III
Rad59 ChIP–seq signal
2000
0
Untreated
As(III)
0 kb
316 kb
Replicate 1
Replicate 2
F
Rad59 ChIP–seq signal
1000
800
600
400
200
0
Untreated
As(III)
-0.5 kb
TSS
TTS
0.5 kb

**Fig. S5**

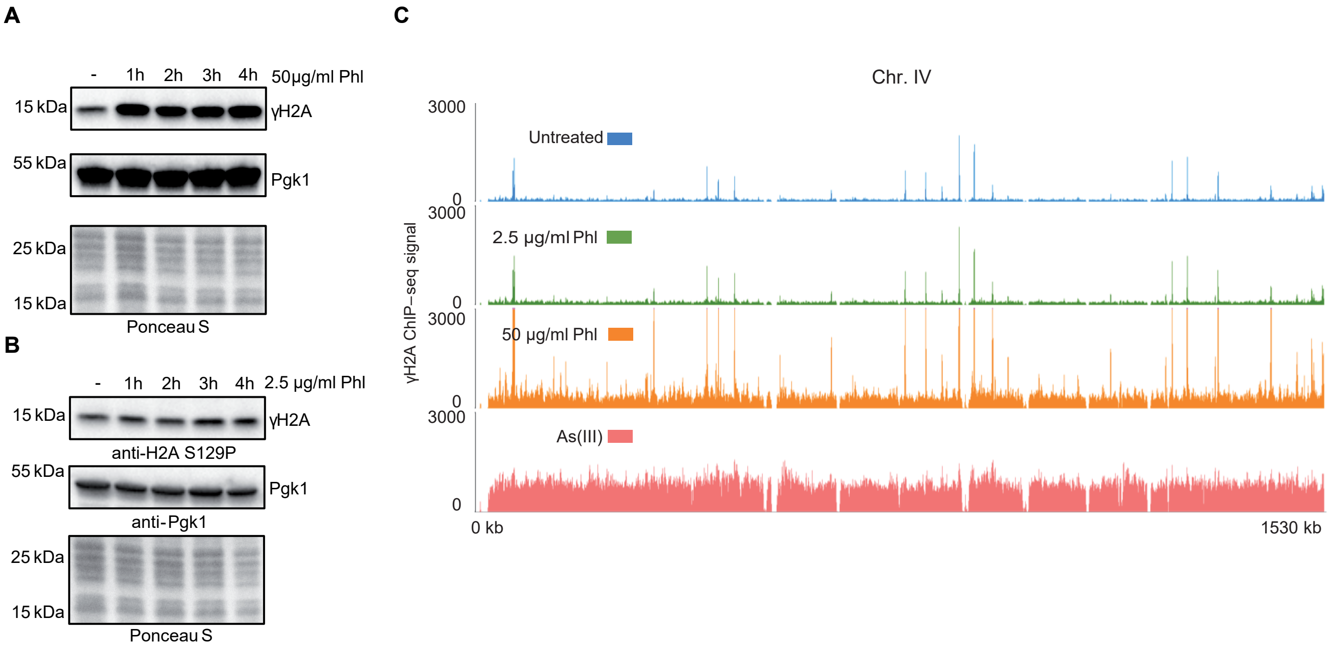
A
-
1h
2h
3h
4h
50μg/ml Phl
15 kDa
γH2A
55 kDa
Pgk1
25 kDa
15 kDa
Ponceau S
B
-
1h
2h
3h
4h
2.5 μg/ml Phl
15 kDa
γH2A
anti-H2A S129P
55 kDa
Pgk1
anti-Pgk1
25 kDa
15 kDa
Ponceau S
C
Chr. IV
3000
0
Untreated
2.5 μg/ml Phl
50 μg/ml Phl
As(III)
γH2A ChIP−seq signal
0 kb
1530 kb

Fig. S6

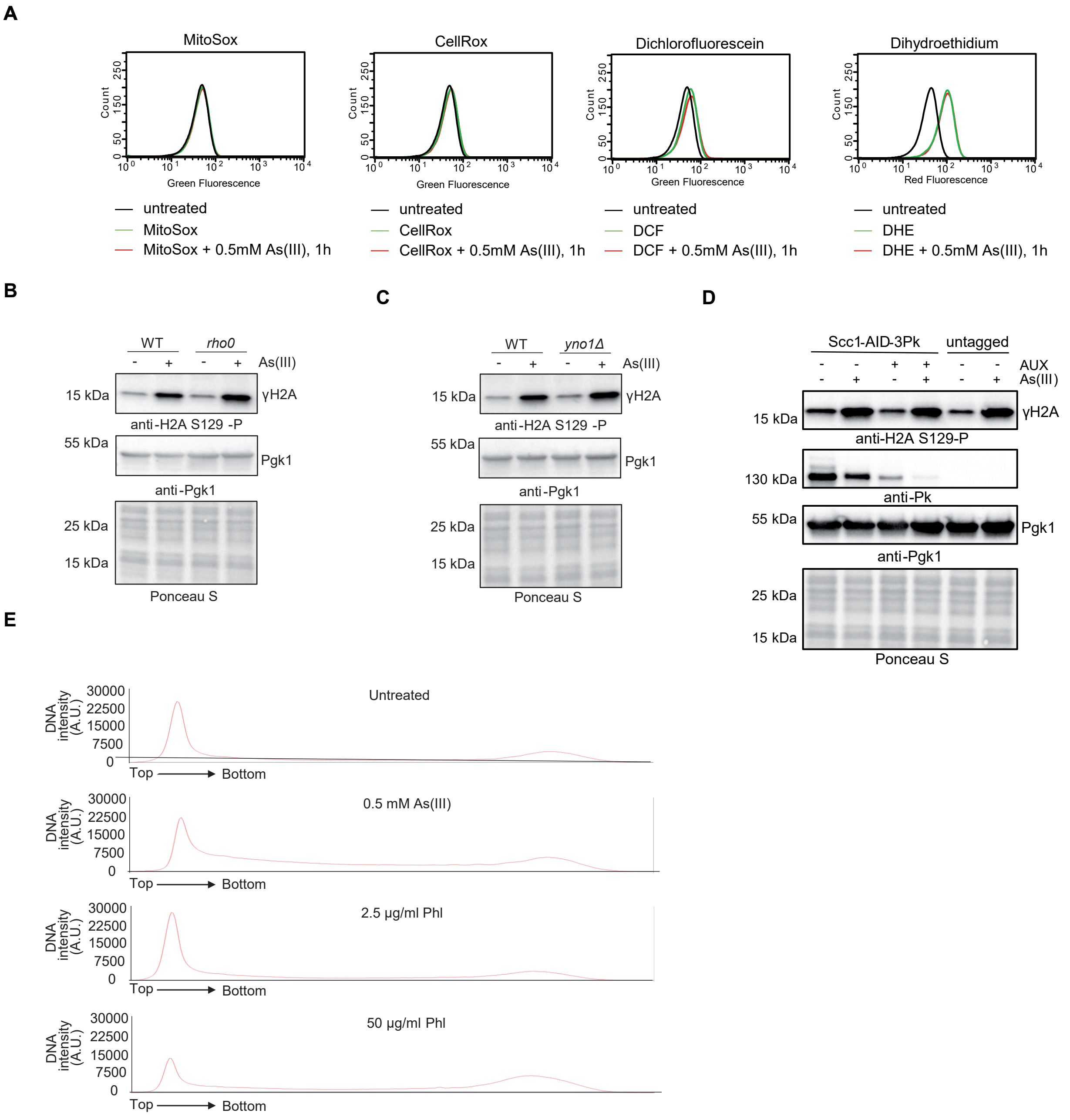
A
MitoSox
CellRox
Dichlorofluorescein
Dihydroethidium
Count
Green Fluorescence
Red Fluorescence
untreated
MitoSox
MitoSox + 0.5mM As(III), 1h
CellRox
CellRox + 0.5mM As(III), 1h
DCF
DCF + 0.5mM As(III), 1h
DHE
DHE + 0.5mM As(III), 1h
B
WT
rho0
As(III)
15 kDa
γH2A
anti-H2A S129 -P
55 kDa
Pgk1
anti-Pgk1
25 kDa
Ponceau S
C
yno1Δ
D
Scc1-AID-3Pk
untagged
AUX
anti-H2A S129-P
130 kDa
anti-Pk
E
DNA intensity (A.U.)
Untreated
0.5 mM As(III)
2.5 µg/ml Phl
50 µg/ml Phl
Top
Bottom

Fig. S7

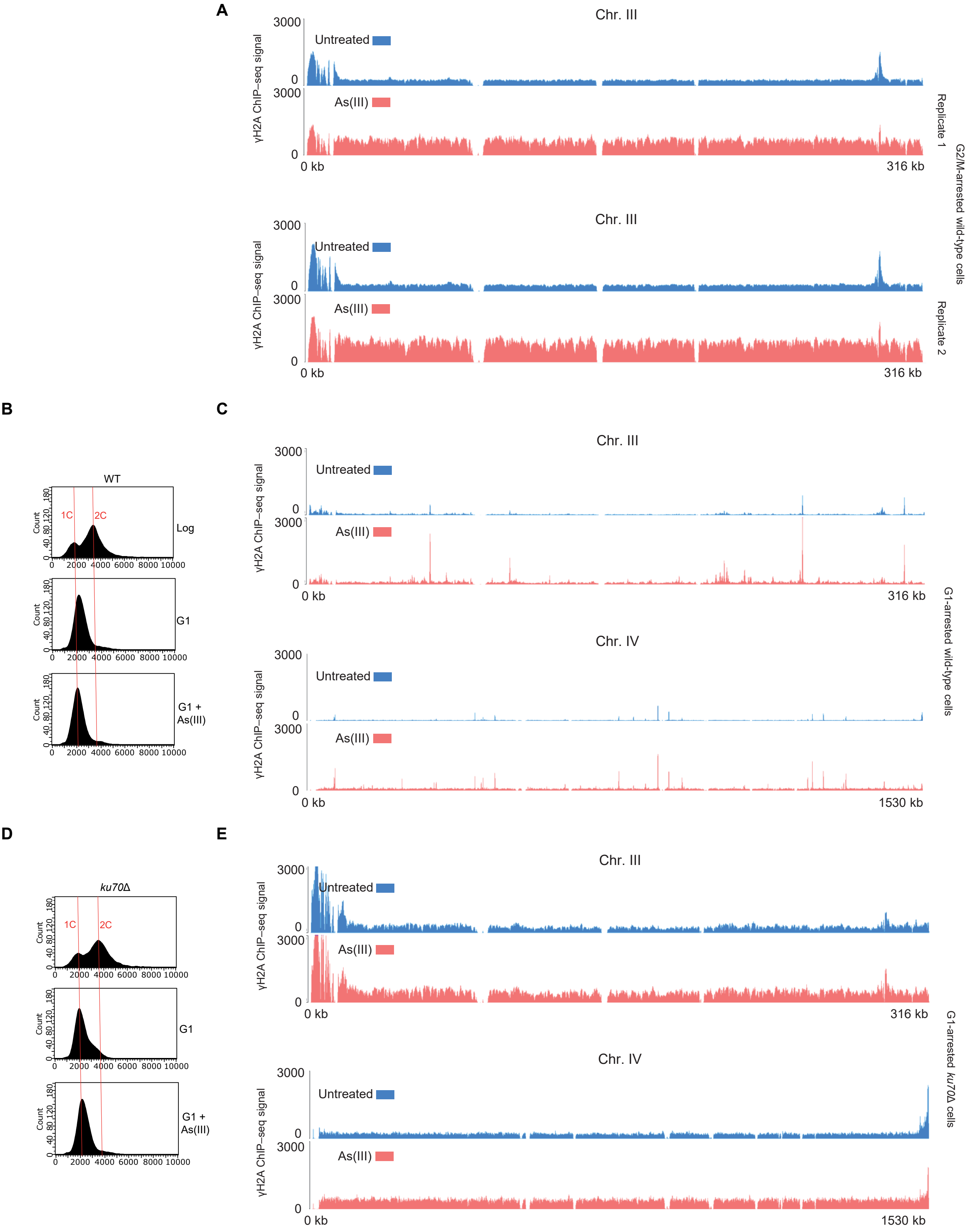
A
Chr. III
γH2A ChIP−seq signal
3000
0
Untreated
As(III)
0 kb
316 kb
Replicate 1
Replicate 2
G2/M-arrested wild-type cells
B
WT
1C
2C
Count
Log
G1
G1 + As(III)
0 2000 4000 6000 8000 10000
C
Chr. III
Chr. IV
1530 kb
G1-arrested wild-type cells
D
ku70Δ
E
G1-arrested ku70Δ cells

**Fig. S8**

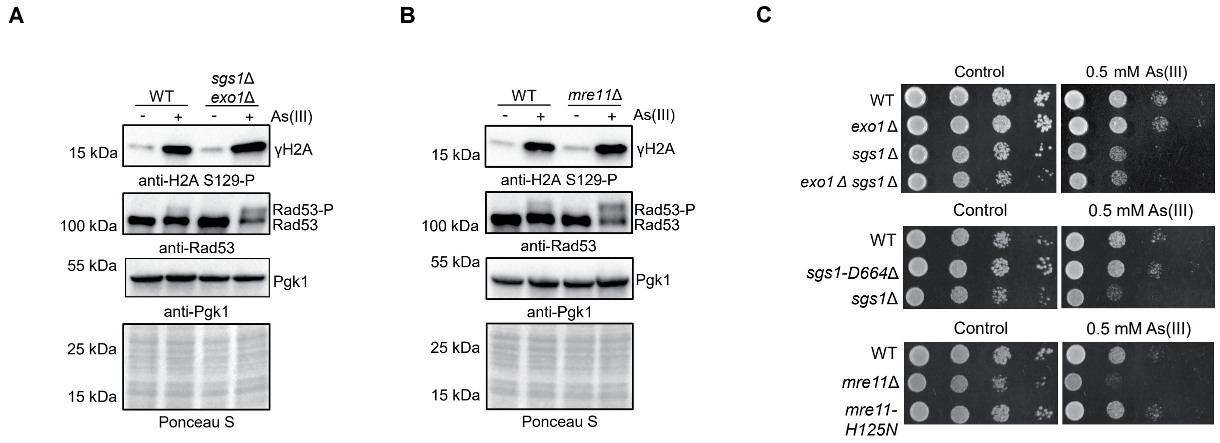
A
WT
sgs1Δ exo1Δ
- + - + As(III)
15 kDa
γH2A
anti-H2A S129-P
Rad53-P
Rad53
100 kDa
anti-Rad53
55 kDa
Pgk1
anti-Pgk1
25 kDa
15 kDa
Ponceau S
B
WT
mre11Δ
- + - + As(III)
15 kDa
γH2A
anti-H2A S129-P
Rad53-P
Rad53
100 kDa
anti-Rad53
55 kDa
Pgk1
anti-Pgk1
25 kDa
15 kDa
Ponceau S
C
Control
0.5 mM As(III)
WT
exo1 Δ
sgs1 Δ
exo1 Δ sgs1 Δ
Control
0.5 mM As(III)
WT
sgs1-D664Δ
sgs1Δ
Control
0.5 mM As(III)
WT
mre11Δ
mre11-H125N

**Fig. S9**

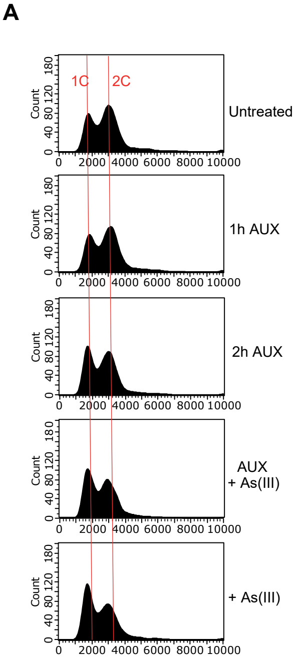
A
1C
2C
Count
Untreated
1h AUX
2h AUX
AUX
+ As(III)
+ As(III)
0 2000 4000 6000 8000 10000
0 40 80 120 180

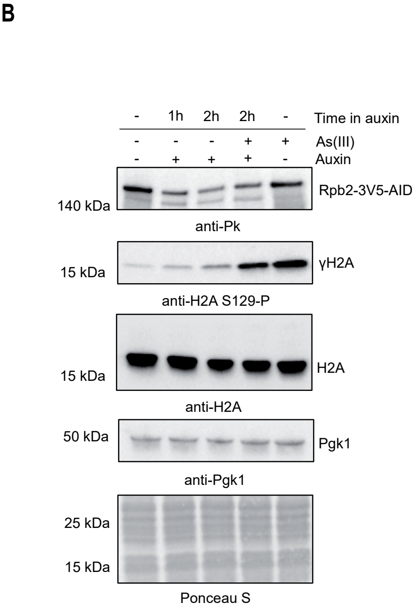
B
Time in auxin
- 1h 2h 2h -
As(III)
- - - + +
Auxin
- + + + -
140 kDa
Rpb2-3V5-AID
anti-Pk
15 kDa
γH2A
anti-H2A S129-P
15 kDa
H2A
anti-H2A
50 kDa
Pgk1
anti-Pgk1
25 kDa
15 kDa
Ponceau S

Fig. S10

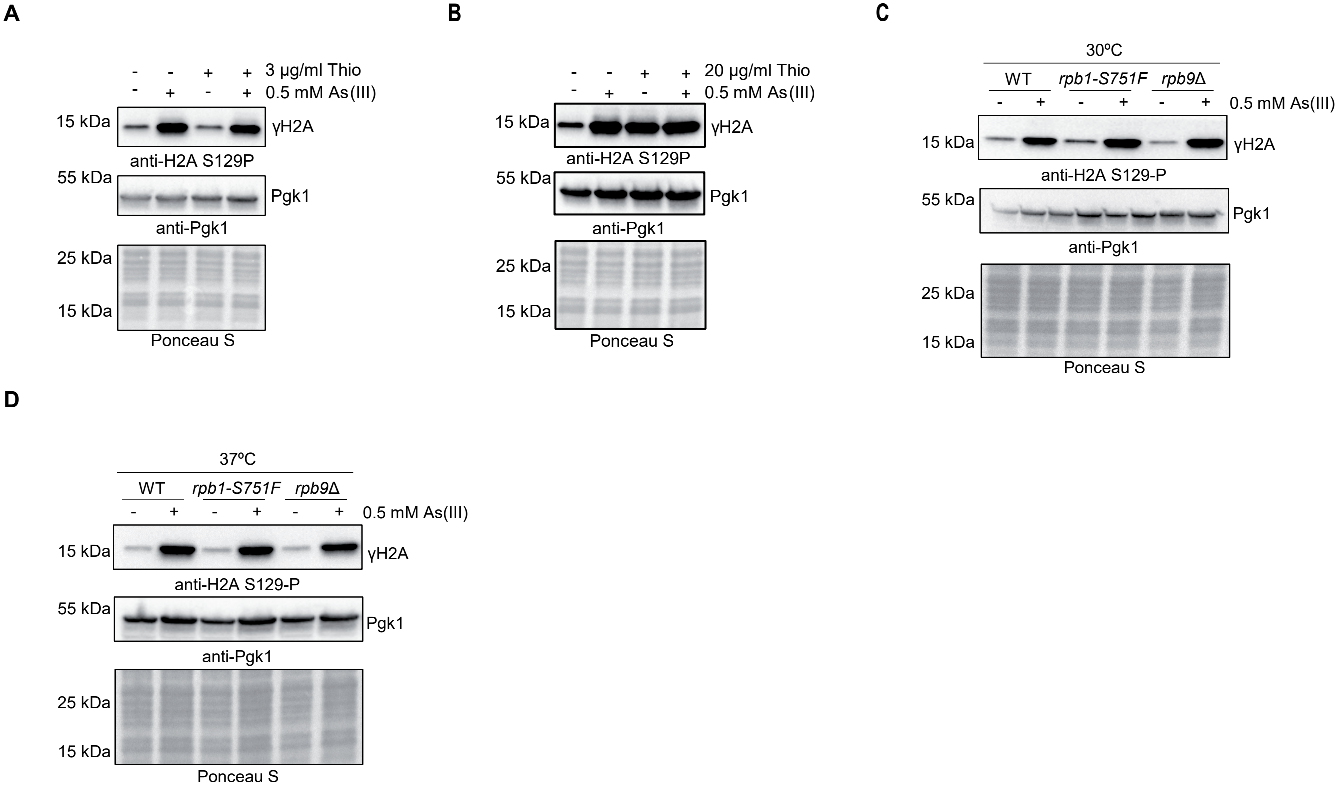
A
- - + + 3 µg/ml Thio
- + - + 0.5 mM As(III)
15 kDa
γH2A
anti-H2A S129P
55 kDa
Pgk1
anti-Pgk1
25 kDa
15 kDa
Ponceau S
B
- - + + 20 µg/ml Thio
- + - + 0.5 mM As(III)
15 kDa
γH2A
anti-H2A S129P
55 kDa
Pgk1
anti-Pgk1
25 kDa
15 kDa
Ponceau S
C
30ºC
WT
rpb1-S751F
rpb9Δ
- + - + - + 0.5 mM As(III)
15 kDa
γH2A
anti-H2A S129-P
55 kDa
Pgk1
anti-Pgk1
25 kDa
15 kDa
Ponceau S
D
37ºC
WT
rpb1-S751F
rpb9Δ
- + - + - + 0.5 mM As(III)
15 kDa
γH2A
anti-H2A S129-P
55 kDa
Pgk1
anti-Pgk1
25 kDa
15 kDa
Ponceau S

Fig. S11

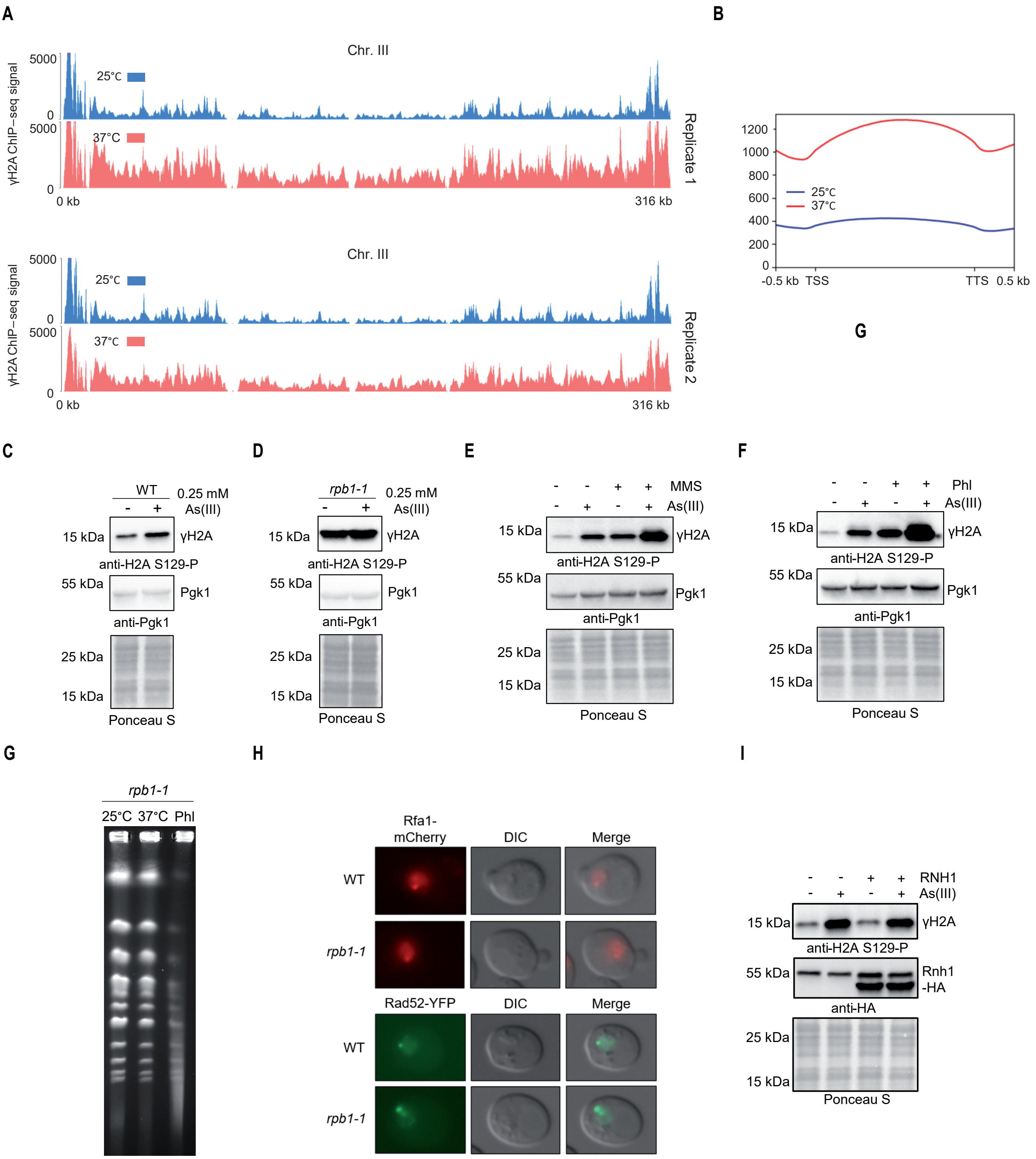

A
Chr. III
γH2A ChIP−seq signal
5000
0
25°C
37°C
0 kb
316 kb
Replicate 1
Replicate 2
B
1200
1000
800
600
400
200
0
25°C
37°C
-0.5 kb
TSS
TTS
0.5 kb
G
C
WT
0.25 mM As(III)
15 kDa
γH2A
anti-H2A S129-P
55 kDa
Pgk1
anti-Pgk1
25 kDa
15 kDa
Ponceau S
D
rpb1-1
0.25 mM As(III)
E
MMS
As(III)
F
Phl
As(III)
G
rpb1-1
25°C 37°C Phl
H
Rfa1-mCherry
DIC
Merge
WT
rpb1-1
Rad52-YFP
I
RNH1
As(III)
Rnh1 -HA
anti-HA

**Fig. S12**

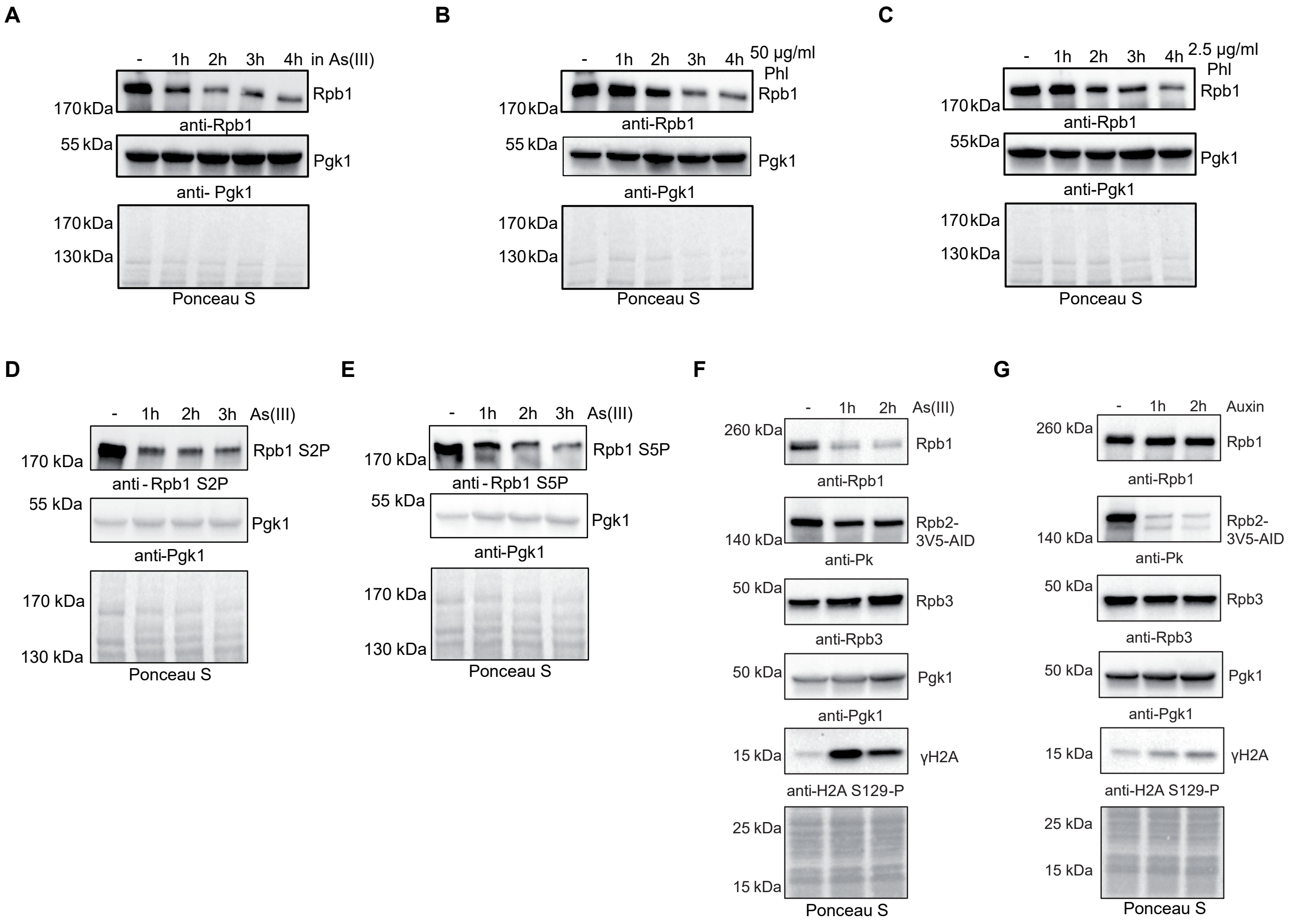
A
- 1h 2h 3h 4h in As(III)
170 kDa
Rpb1
anti-Rpb1
55 kDa
Pgk1
anti- Pgk1
170 kDa
130 kDa
Ponceau S
B
- 1h 2h 3h 4h 50 µg/ml Phl
170 kDa
Rpb1
anti-Rpb1
55 kDa
Pgk1
anti-Pgk1
170 kDa
130 kDa
Ponceau S
C
- 1h 2h 3h 4h 2.5 µg/ml Phl
170 kDa
Rpb1
anti-Rpb1
55kDa
Pgk1
anti-Pgk1
170 kDa
130 kDa
Ponceau S
D
- 1h 2h 3h As(III)
170 kDa
Rpb1 S2P
anti - Rpb1 S2P
55 kDa
Pgk1
anti-Pgk1
170 kDa
130 kDa
Ponceau S
E
- 1h 2h 3h As(III)
170 kDa
Rpb1 S5P
anti - Rpb1 S5P
55 kDa
Pgk1
anti-Pgk1
170 kDa
130 kDa
Ponceau S
F
- 1h 2h As(III)
260 kDa
Rpb1
anti-Rpb1
140 kDa
Rpb2-3V5-AID
anti-Pk
50 kDa
Rpb3
anti-Rpb3
50 kDa
Pgk1
anti-Pgk1
15 kDa
γH2A
anti-H2A S129-P
25 kDa
15 kDa
Ponceau S
G
- 1h 2h Auxin
260 kDa
Rpb1
anti-Rpb1
140 kDa
Rpb2-3V5-AID
anti-Pk
50 kDa
Rpb3
anti-Rpb3
50 kDa
Pgk1
anti-Pgk1
15 kDa
γH2A
anti-H2A S129-P
25 kDa
15 kDa
Ponceau S

Fig. S13

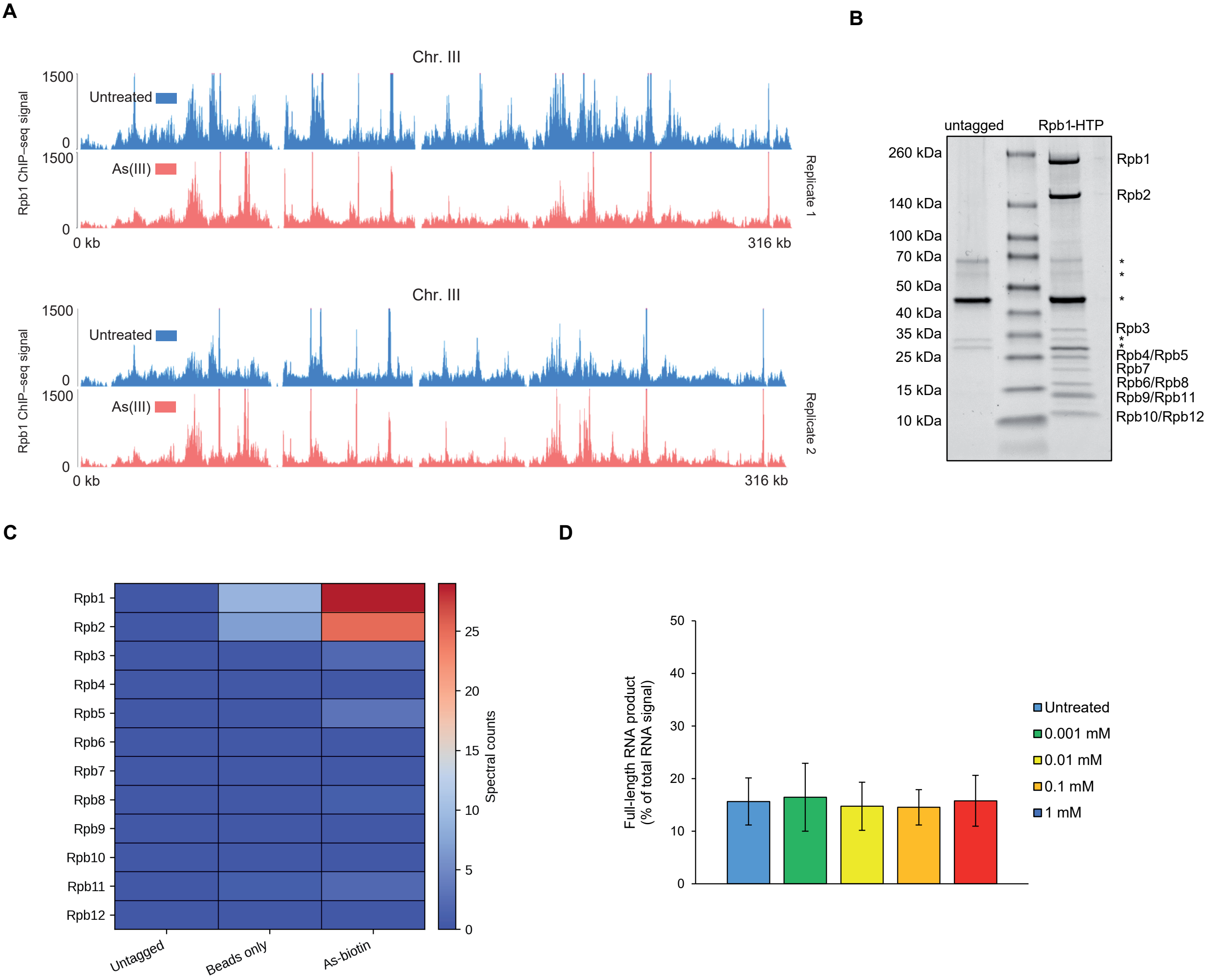
A
B
Chr. III
Rpb1 ChIP–seq signal
1500
0
Untreated
As(III)
0 kb
316 kb
Replicate 1
Replicate 2
untagged
Rpb1-HTP
260 kDa
140 kDa
100 kDa
70 kDa
50 kDa
40 kDa
35 kDa
25 kDa
15 kDa
10 kDa
Rpb1
Rpb2
Rpb3
Rpb4/Rpb5
Rpb7
Rpb6/Rpb8
Rpb9/Rpb11
Rpb10/Rpb12
C
D
Rpb1
Rpb2
Rpb3
Rpb4
Rpb5
Rpb6
Rpb7
Rpb8
Rpb9
Rpb10
Rpb11
Rpb12
Untagged
Beads only
As-biotin
Spectral counts
0
5
10
15
20
25
Full-length RNA product
(% of total RNA signal)
50
40
30
20
10
0
Untreated
0.001 mM
0.01 mM
0.1 mM
1 mM

### Unprocessed, uncropped immunoblot and gel images (Source Data)

**Fig. 1A**

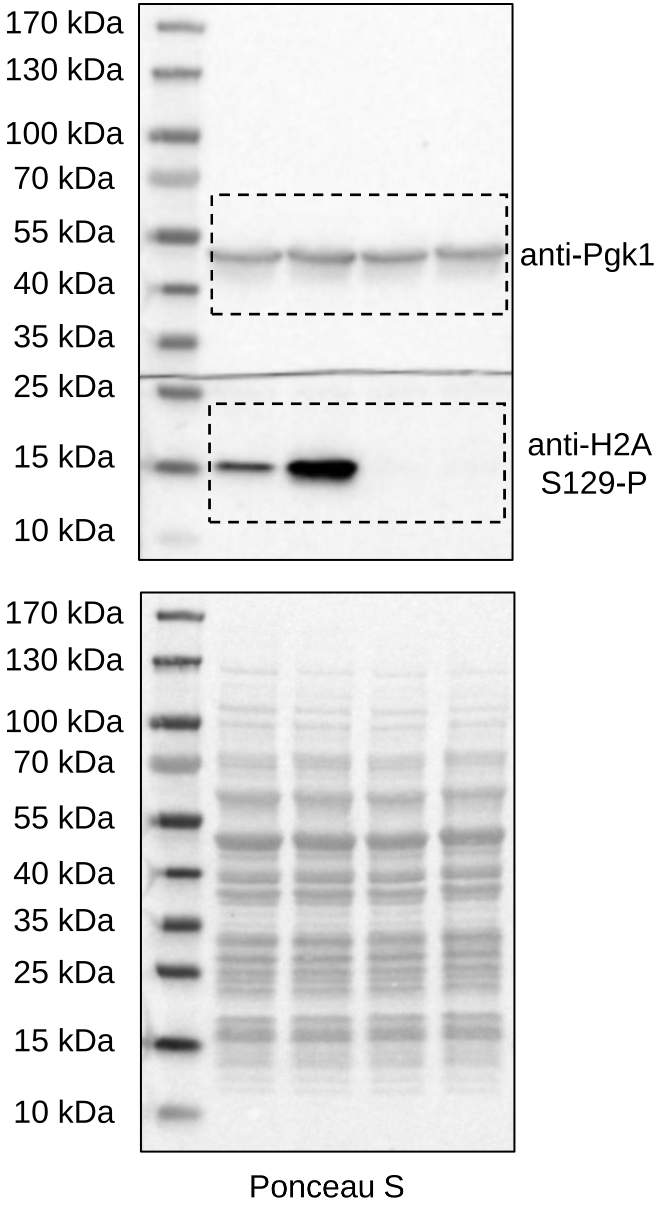
170 kDa
130 kDa
100 kDa
70 kDa
55 kDa
40 kDa
35 kDa
25 kDa
15 kDa
10 kDa
anti-Pgk1
anti-H2A S129-P
170 kDa
130 kDa
100 kDa
70 kDa
55 kDa
40 kDa
35 kDa
25 kDa
15 kDa
10 kDa
Ponceau S

**Fig. 2A**

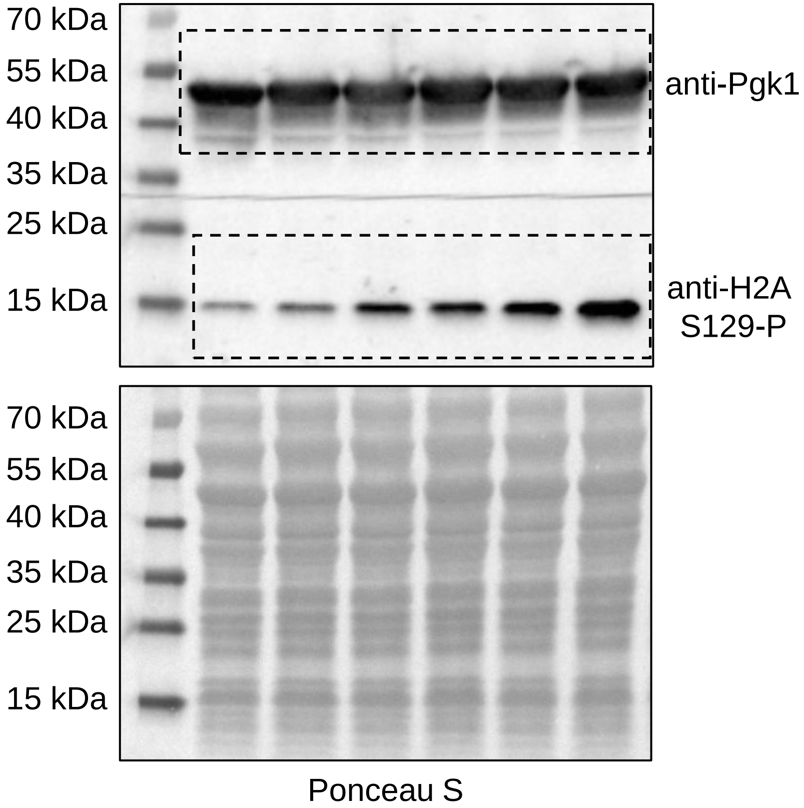
70 kDa
55 kDa
40 kDa
35 kDa
25 kDa
15 kDa
anti-Pgk1
anti-H2A S129-P
70 kDa
55 kDa
40 kDa
35 kDa
25 kDa
15 kDa
Ponceau S

#### Fig. 2B

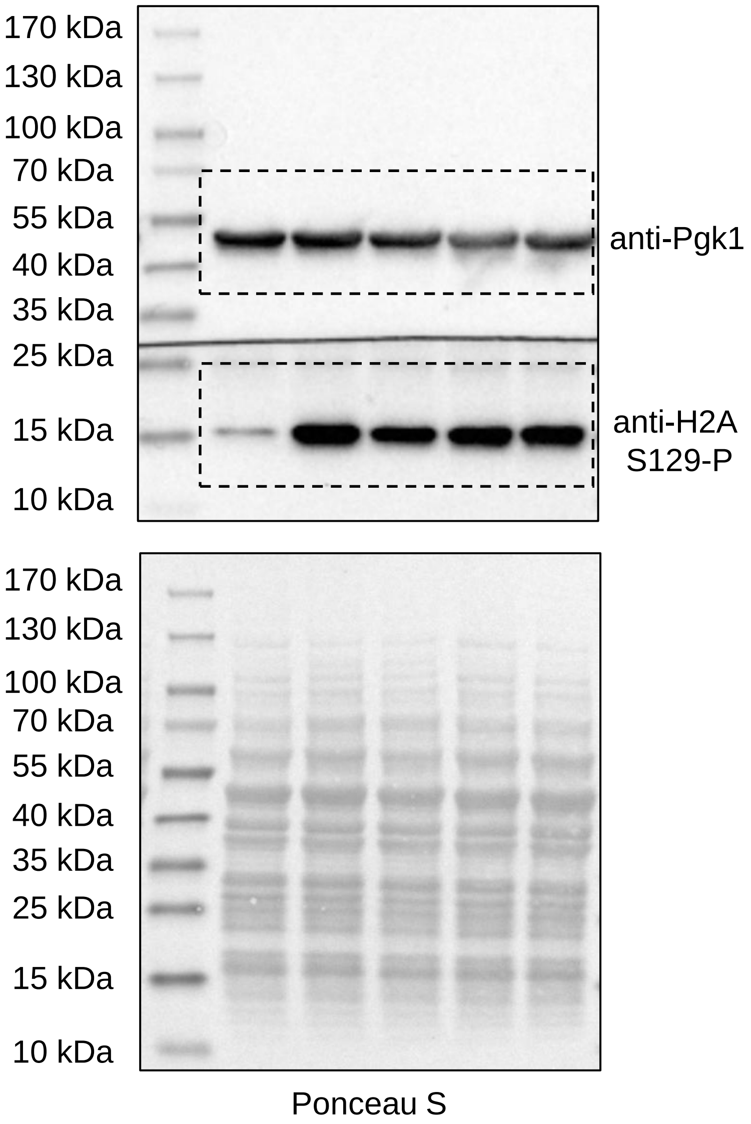
170 kDa
130 kDa
100 kDa
70 kDa
55 kDa
40 kDa
35 kDa
25 kDa
15 kDa
10 kDa
anti-Pgk1
anti-H2A S129-P
Ponceau S

#### Fig. 2E

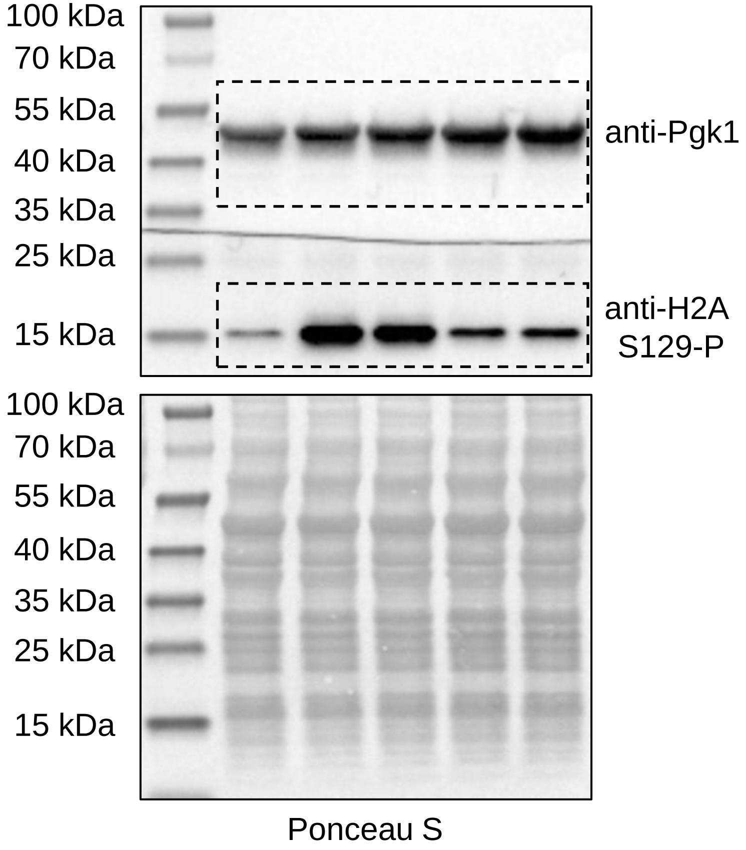
100 kDa
70 kDa
55 kDa
40 kDa
35 kDa
25 kDa
15 kDa
anti-Pgk1
anti-H2A S129-P
Ponceau S

**Fig. 3A**

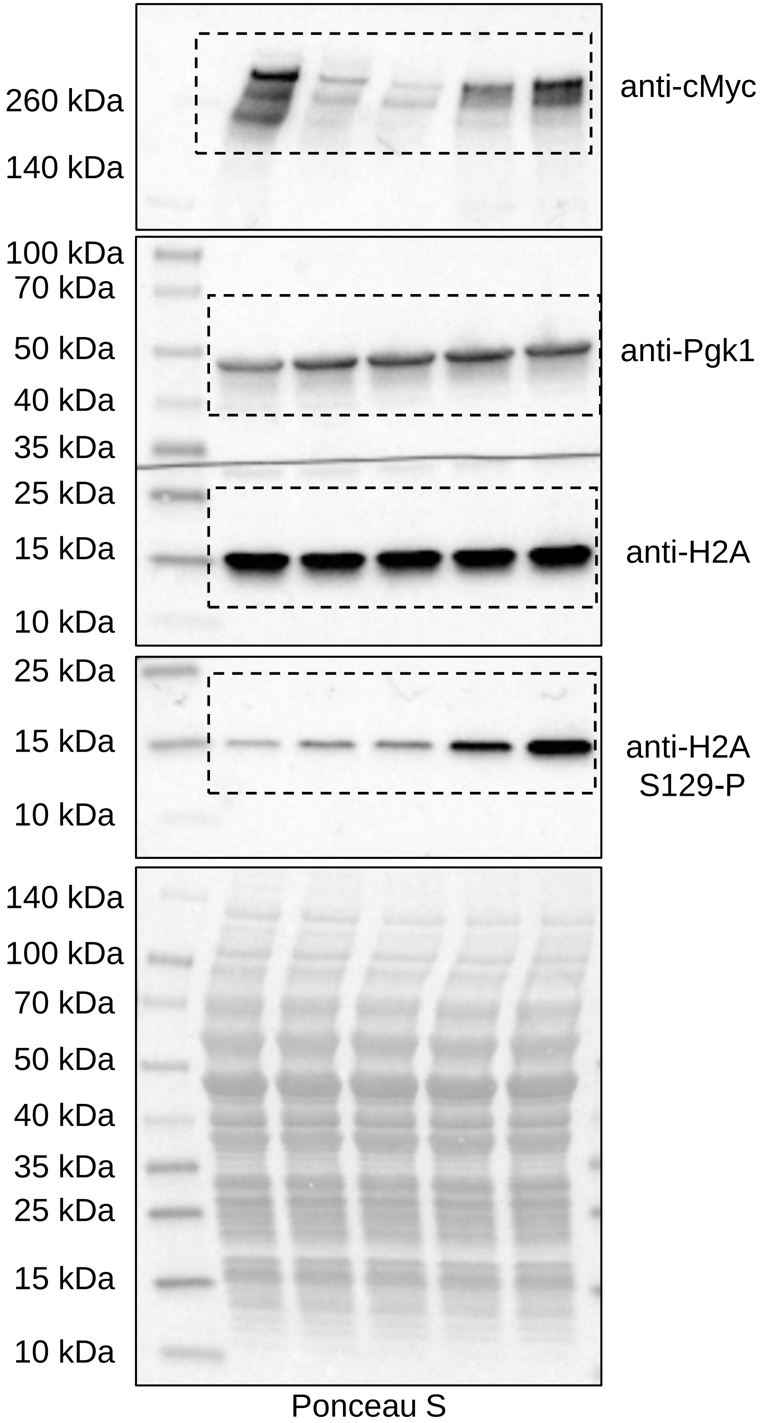

260 kDa
140 kDa
anti-cMyc
100 kDa
70 kDa
50 kDa
40 kDa
35 kDa
25 kDa
15 kDa
10 kDa
anti-Pgk1
anti-H2A
25 kDa
15 kDa
10 kDa
anti-H2A S129-P
140 kDa
100 kDa
70 kDa
50 kDa
40 kDa
35 kDa
25 kDa
15 kDa
10 kDa
Ponceau S


**Fig. 3C**

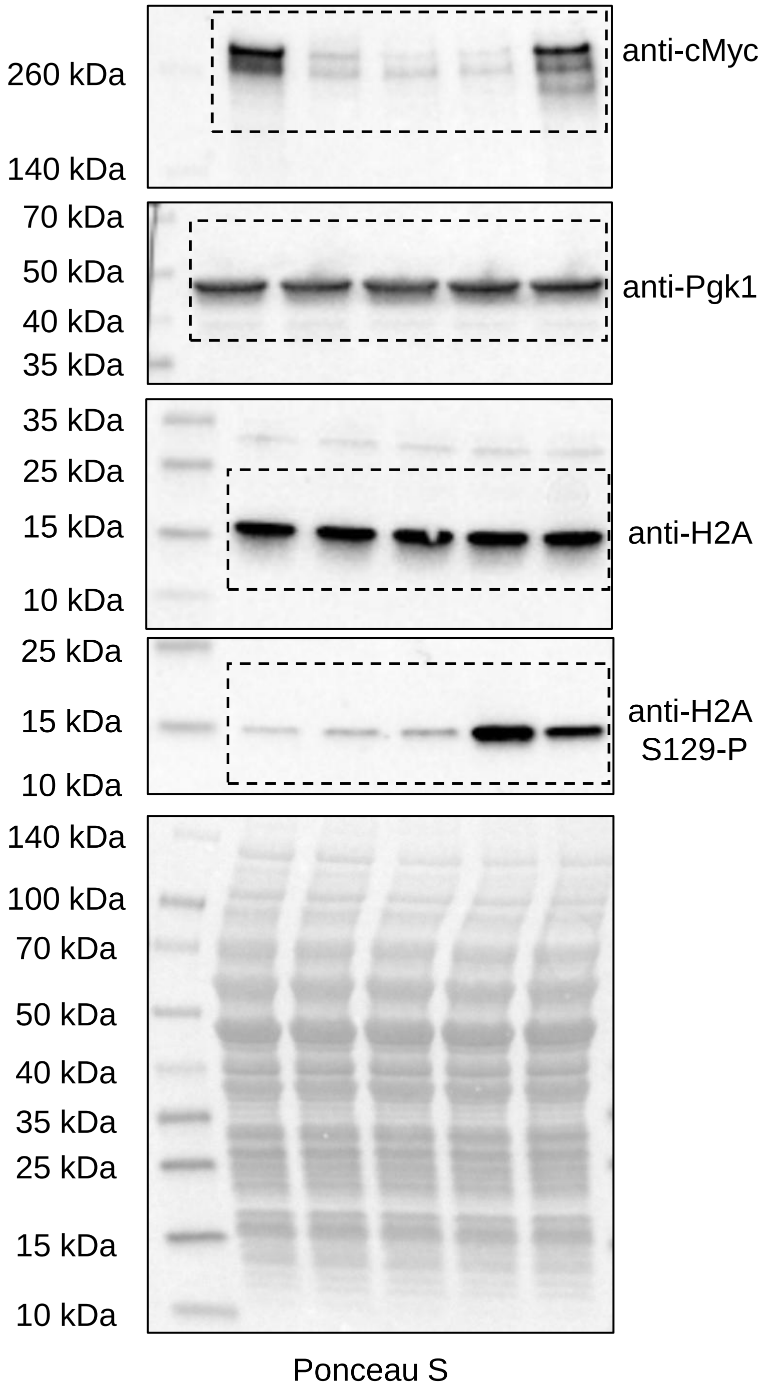

260 kDa
140 kDa
anti-cMyc
70 kDa
50 kDa
40 kDa
35 kDa
anti-Pgk1
35 kDa
25 kDa
15 kDa
10 kDa
anti-H2A
25 kDa
15 kDa
10 kDa
anti-H2A S129-P
140 kDa
100 kDa
70 kDa
50 kDa
40 kDa
35 kDa
25 kDa
15 kDa
10 kDa
Ponceau S

**Fig. 3D**

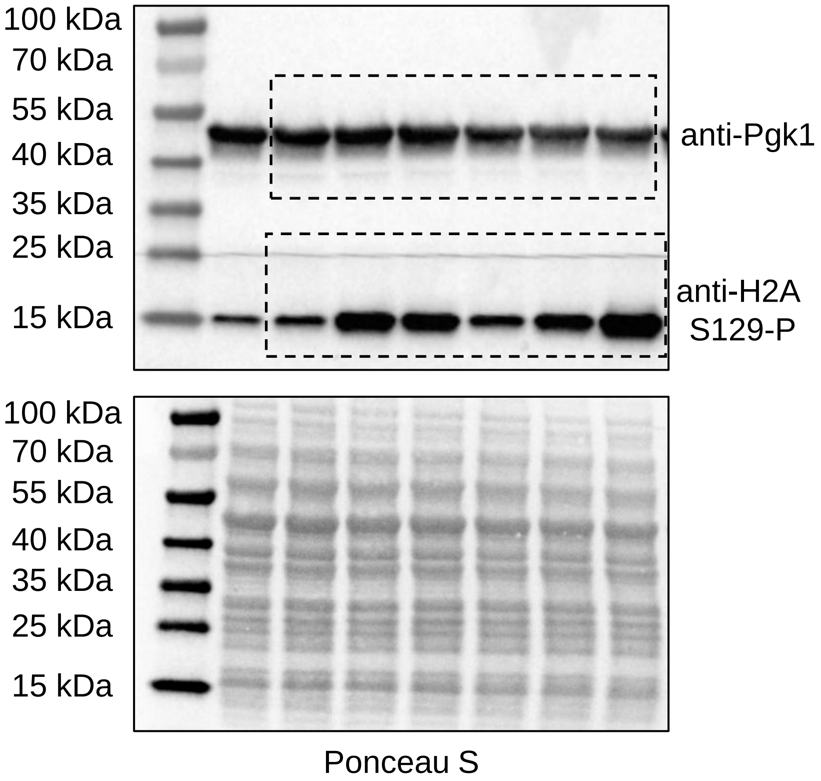
100 kDa
70 kDa
55 kDa
40 kDa
35 kDa
25 kDa
15 kDa
anti-Pgk1
anti-H2A
S129-P
100 kDa
70 kDa
55 kDa
40 kDa
35 kDa
25 kDa
15 kDa
Ponceau S

**Fig. 3E**

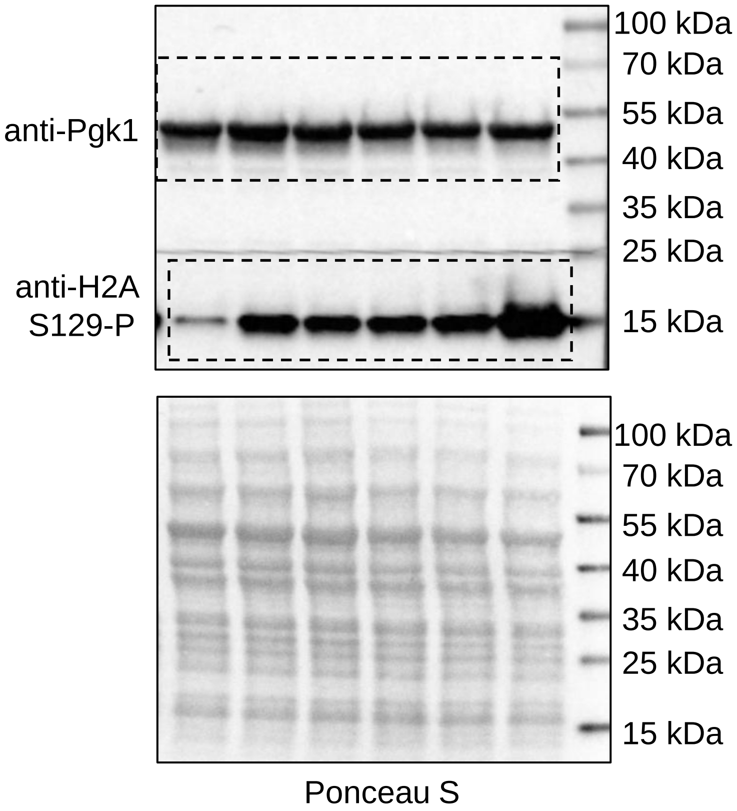
anti-Pgk1
anti-H2A
S129-P
100 kDa
70 kDa
55 kDa
40 kDa
35 kDa
25 kDa
15 kDa
100 kDa
70 kDa
55 kDa
40 kDa
35 kDa
25 kDa
15 kDa
Ponceau S

**Fig. 3G**

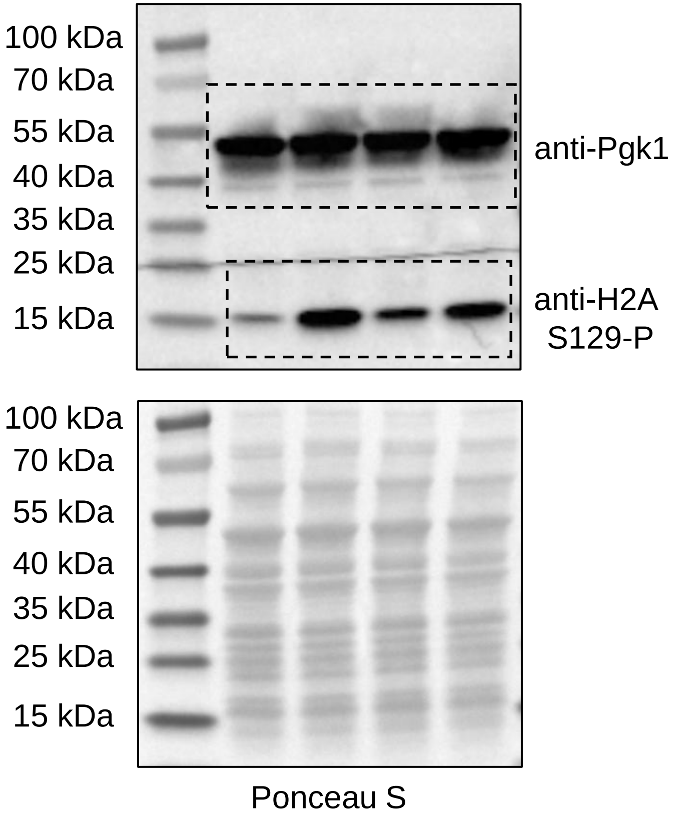
100 kDa
70 kDa
55 kDa
40 kDa
35 kDa
25 kDa
15 kDa
anti-Pgk1
anti-H2A S129-P
100 kDa
70 kDa
55 kDa
40 kDa
35 kDa
25 kDa
15 kDa
Ponceau S

**Fig. 3H**

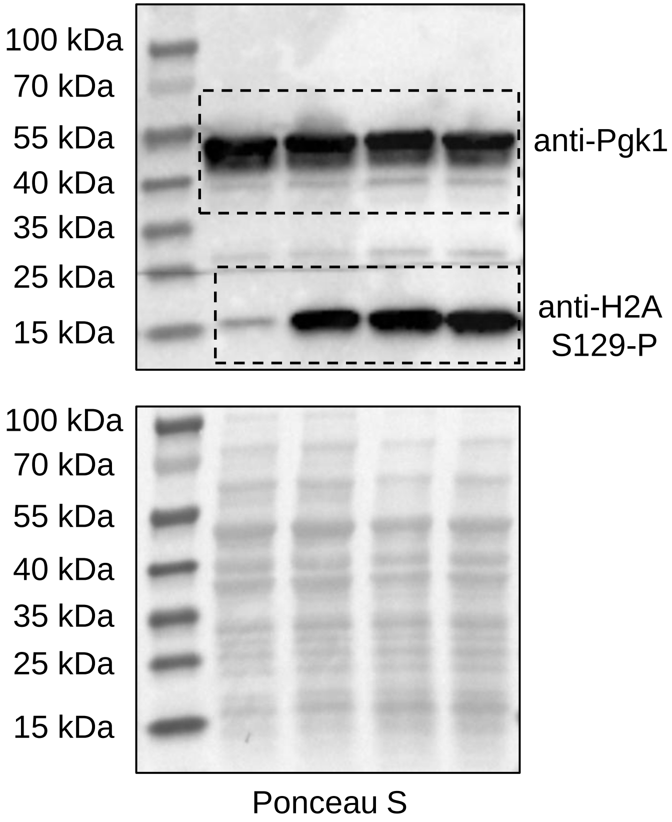
100 kDa
70 kDa
55 kDa
40 kDa
35 kDa
25 kDa
15 kDa
anti-Pgk1
anti-H2A S129-P
100 kDa
70 kDa
55 kDa
40 kDa
35 kDa
25 kDa
15 kDa
Ponceau S

**Fig. 4A**

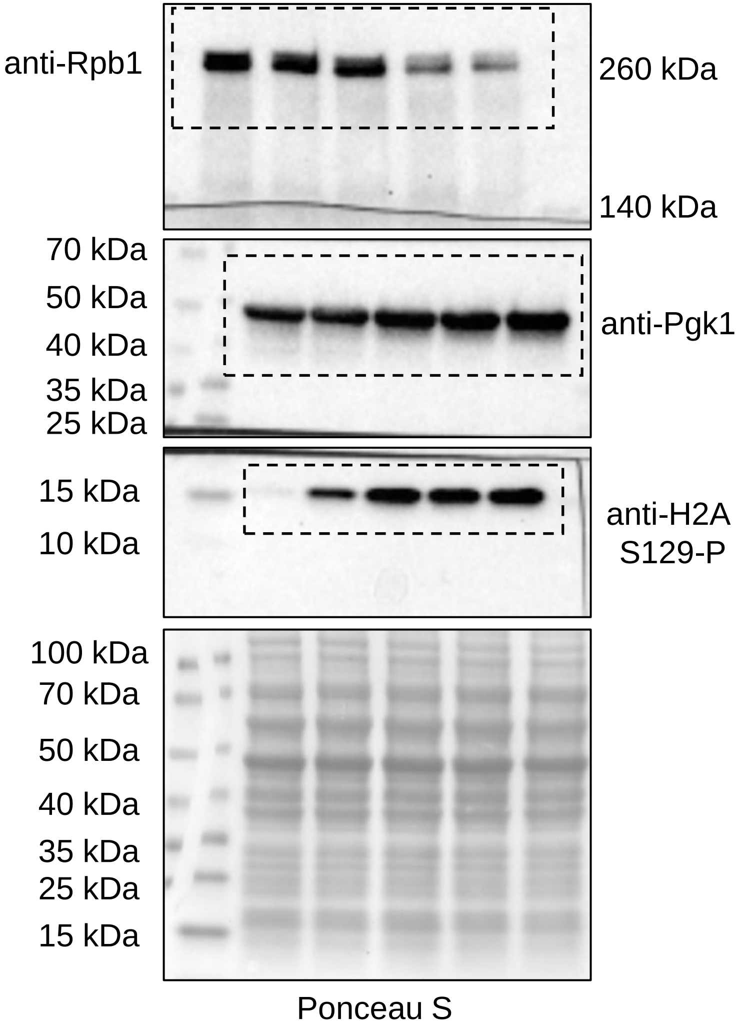
anti-Rpb1
260 kDa
140 kDa
70 kDa
50 kDa
40 kDa
35 kDa
25 kDa
anti-Pgk1
15 kDa
10 kDa
anti-H2A S129-P
100 kDa
70 kDa
50 kDa
40 kDa
35 kDa
25 kDa
15 kDa
Ponceau S

**Fig. 4D**

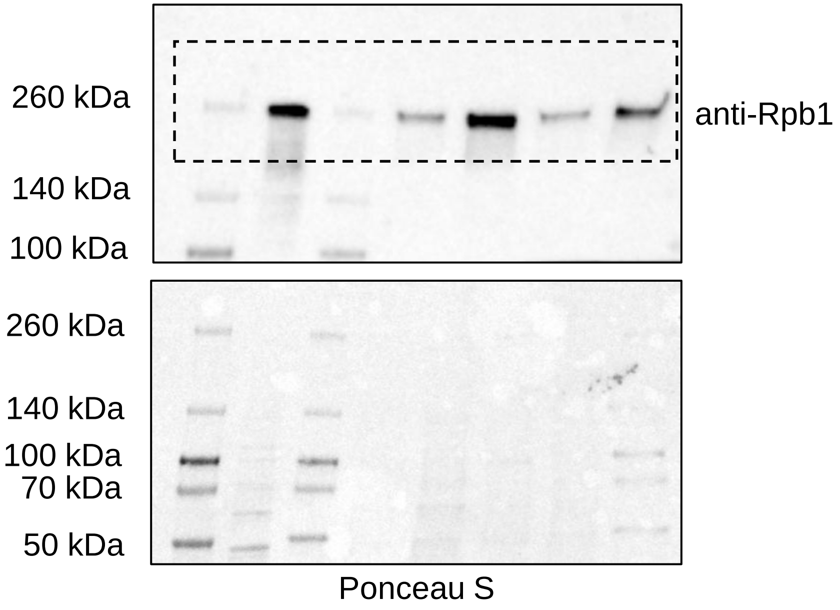
260 kDa
140 kDa
100 kDa
anti-Rpb1
260 kDa
140 kDa
100 kDa
70 kDa
50 kDa
Ponceau S

**Fig. 5E**

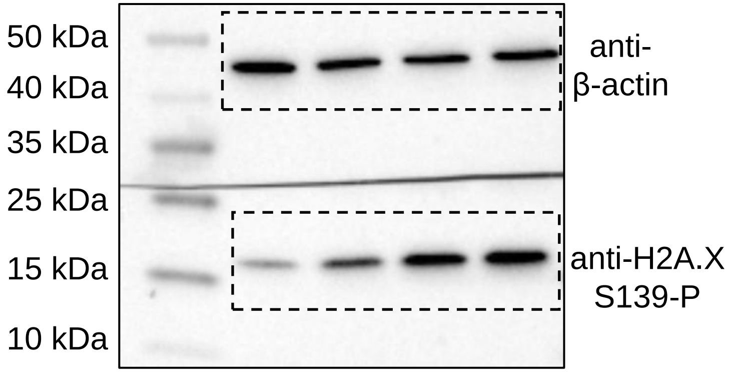


**Fig. 5E**

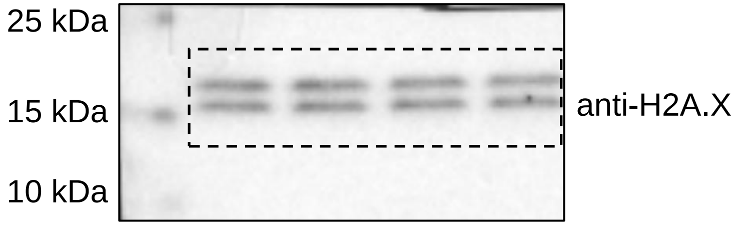


**Fig. 5E**

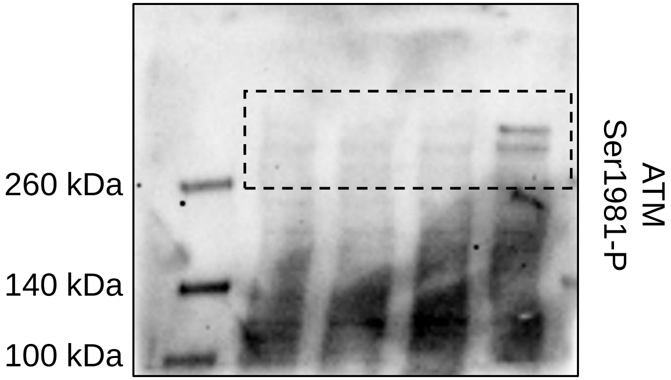


**Fig. 5E**

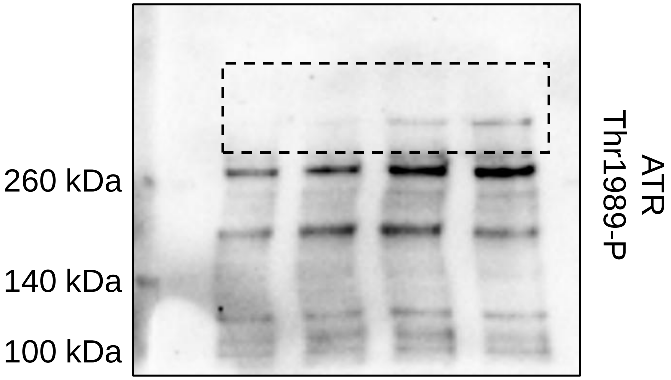


**Fig. 5H**

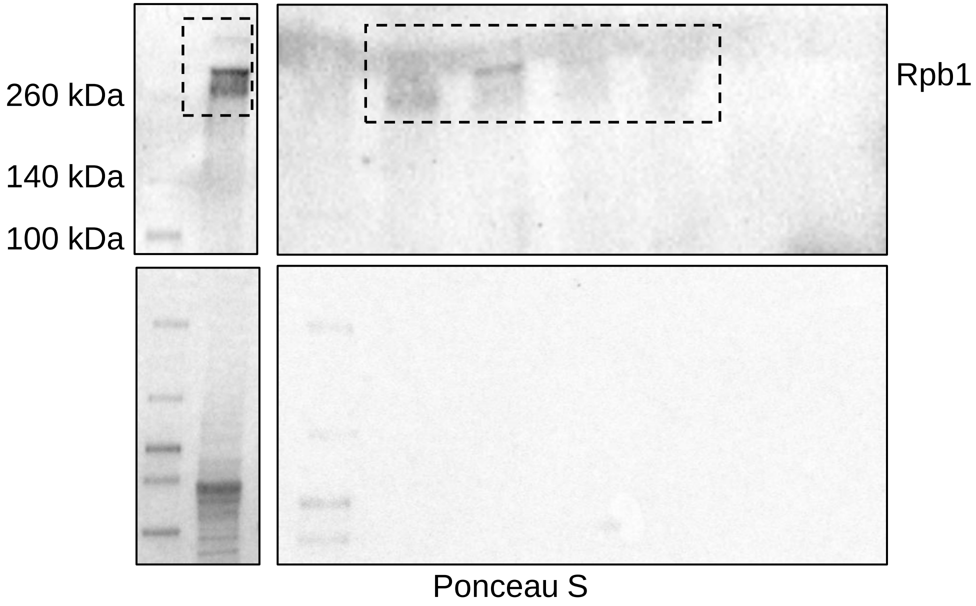

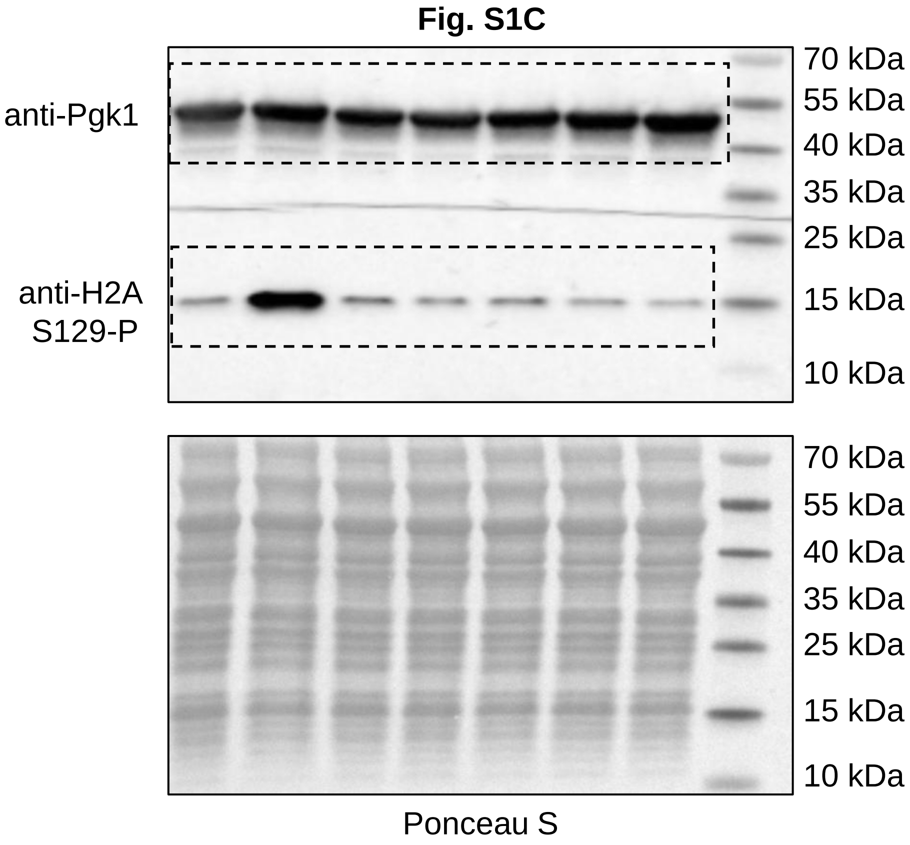
Fig. S1C
anti-Pgk1
anti-H2A
S129-P
70 kDa
55 kDa
40 kDa
35 kDa
25 kDa
15 kDa
10 kDa
70 kDa
55 kDa
40 kDa
35 kDa
25 kDa
15 kDa
10 kDa
Ponceau S

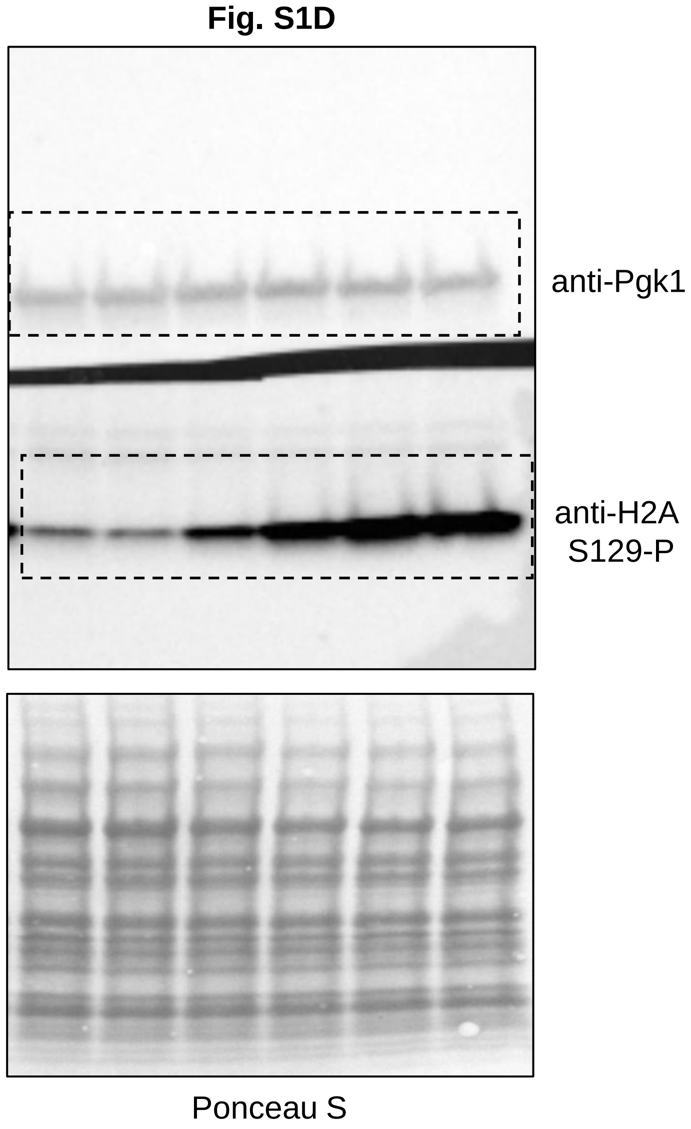
Fig. S1D
anti-Pgk1
anti-H2A
S129-P
Ponceau S

**Fig. S1E**

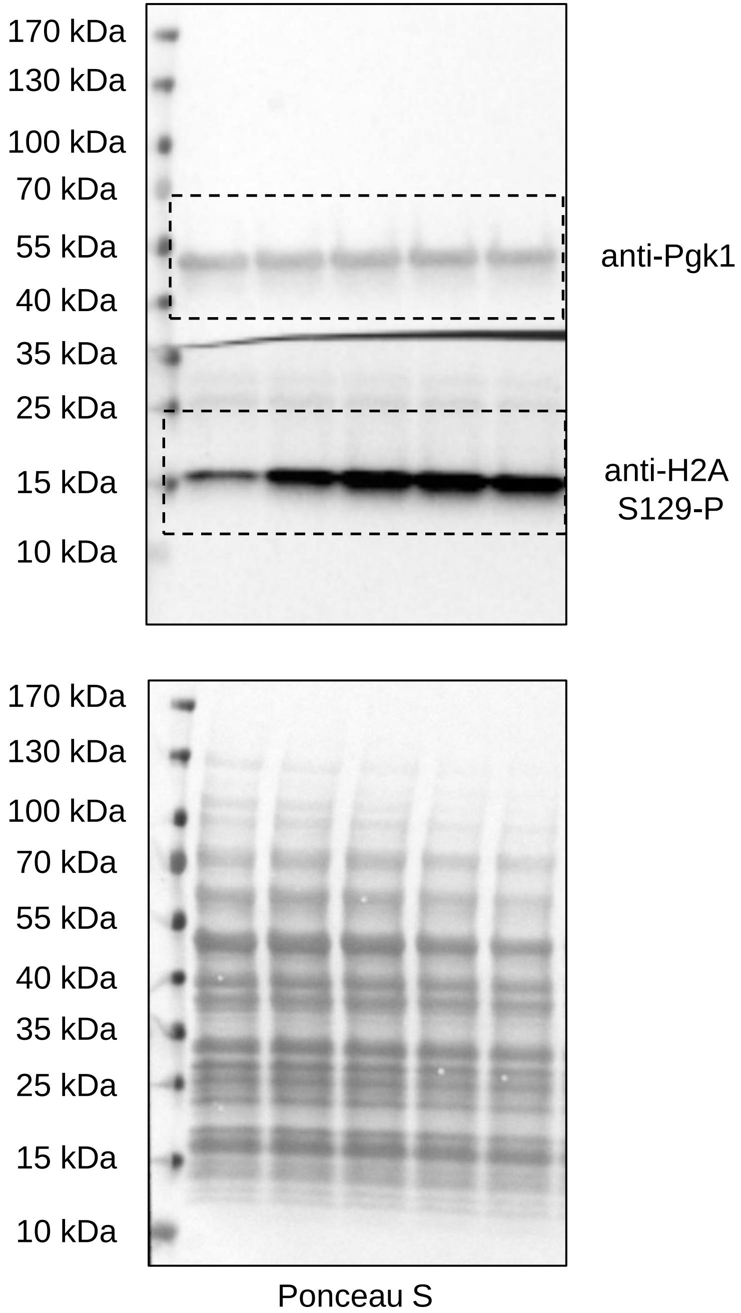


**Fig. S1F**

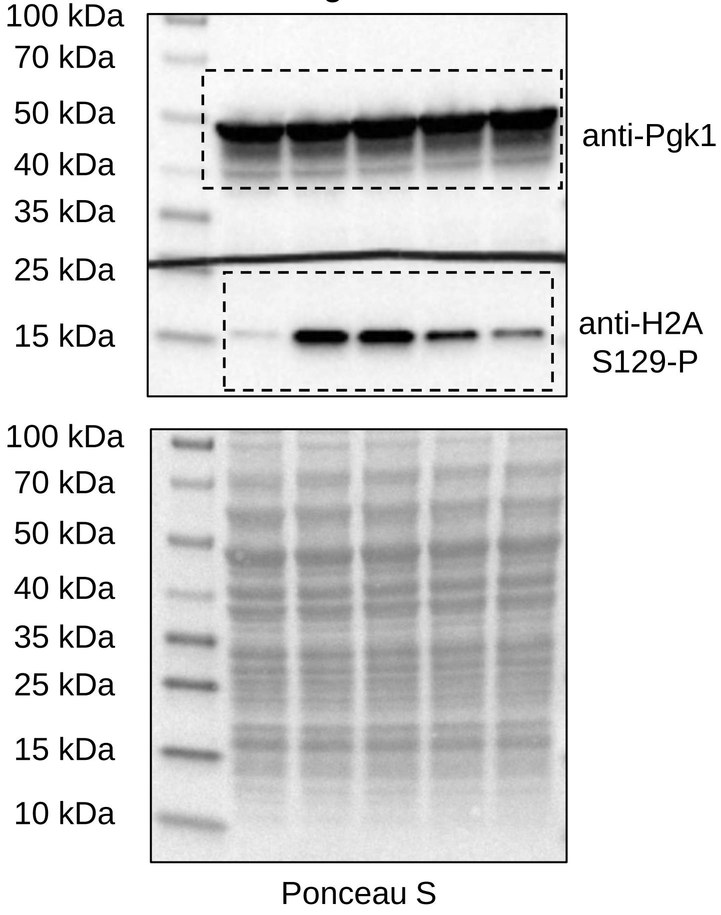

**Fig. S5A**

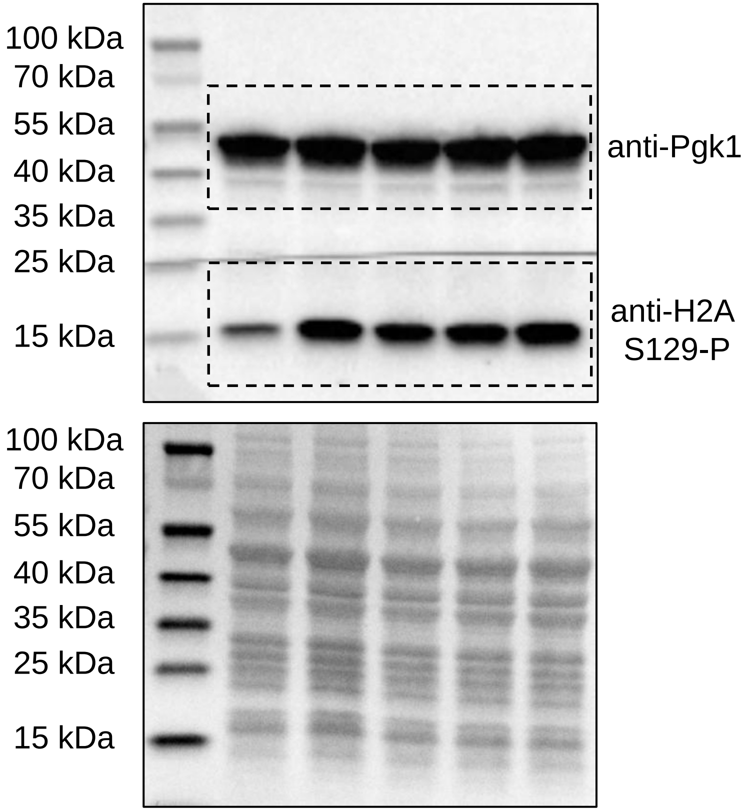


Ponceau S

**Fig. S5B**

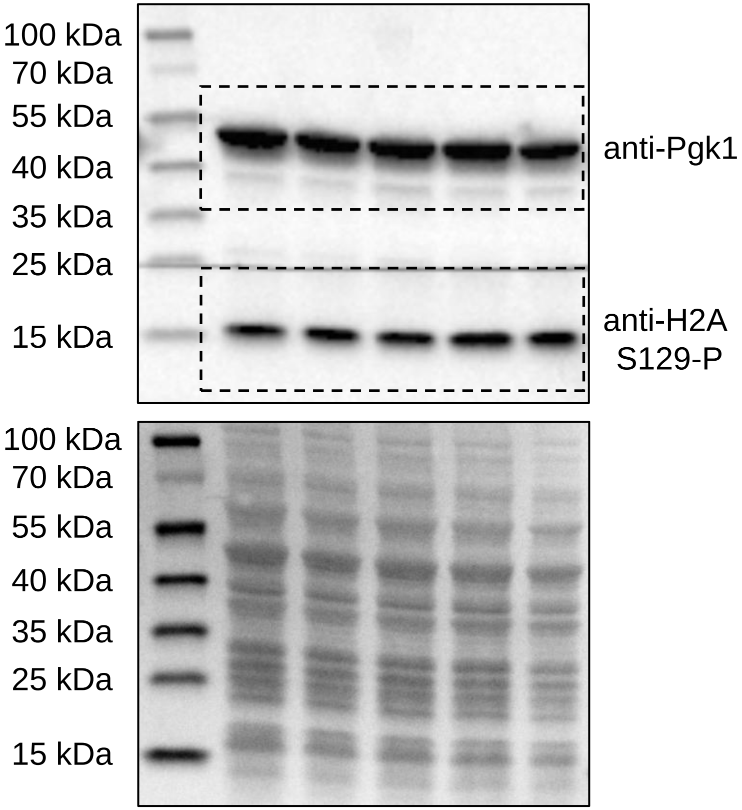


Ponceau S

**Fig. S6B**

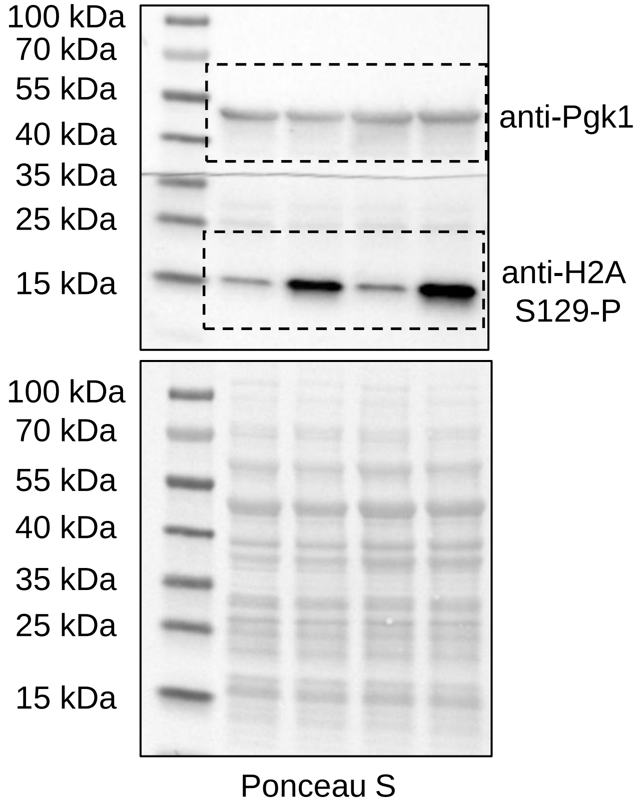


**Fig. S6C**

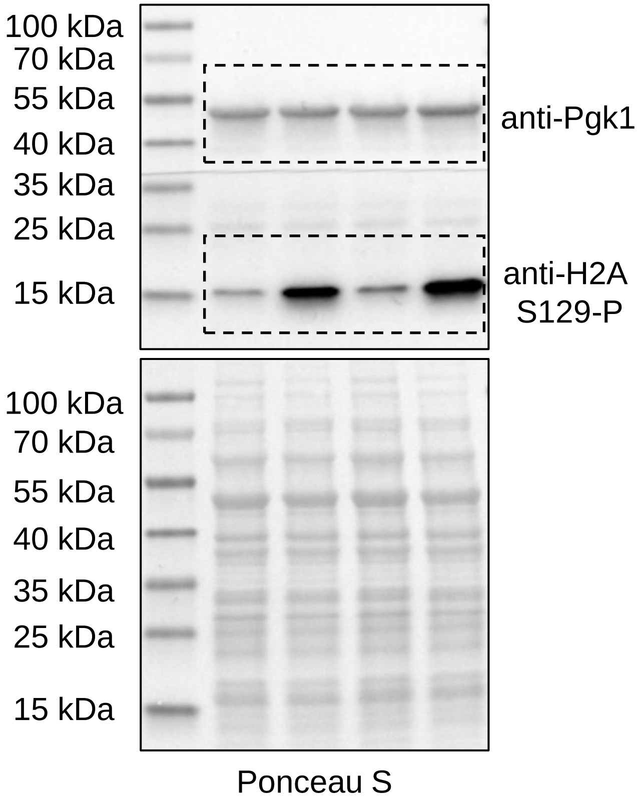


**Fig. S6D**

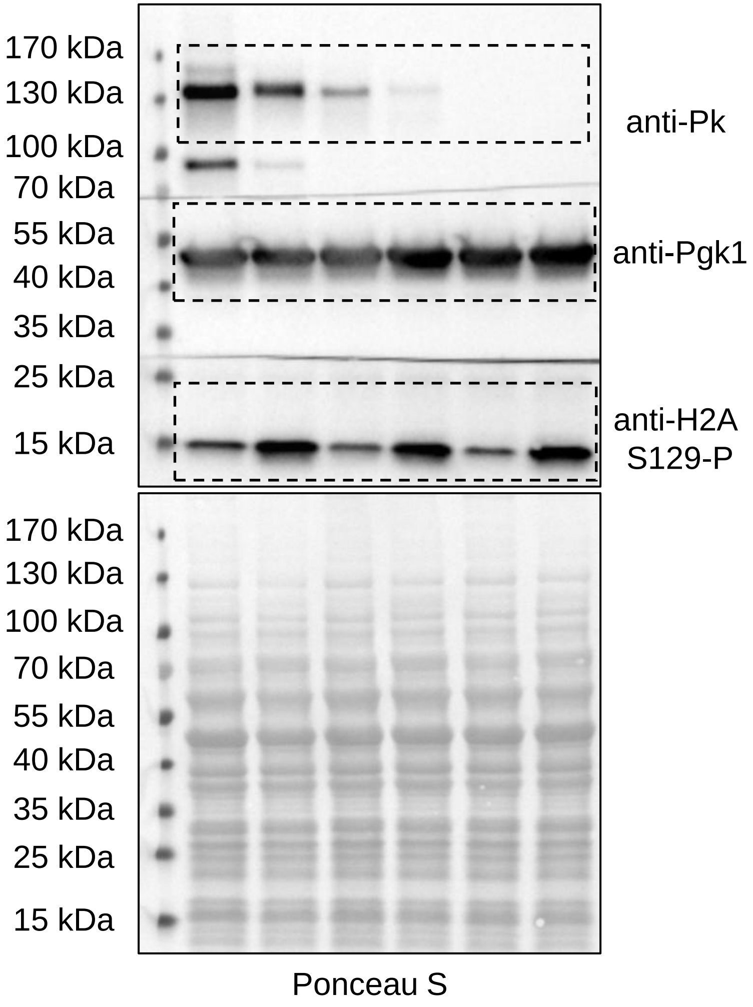

**Fig. S8A**

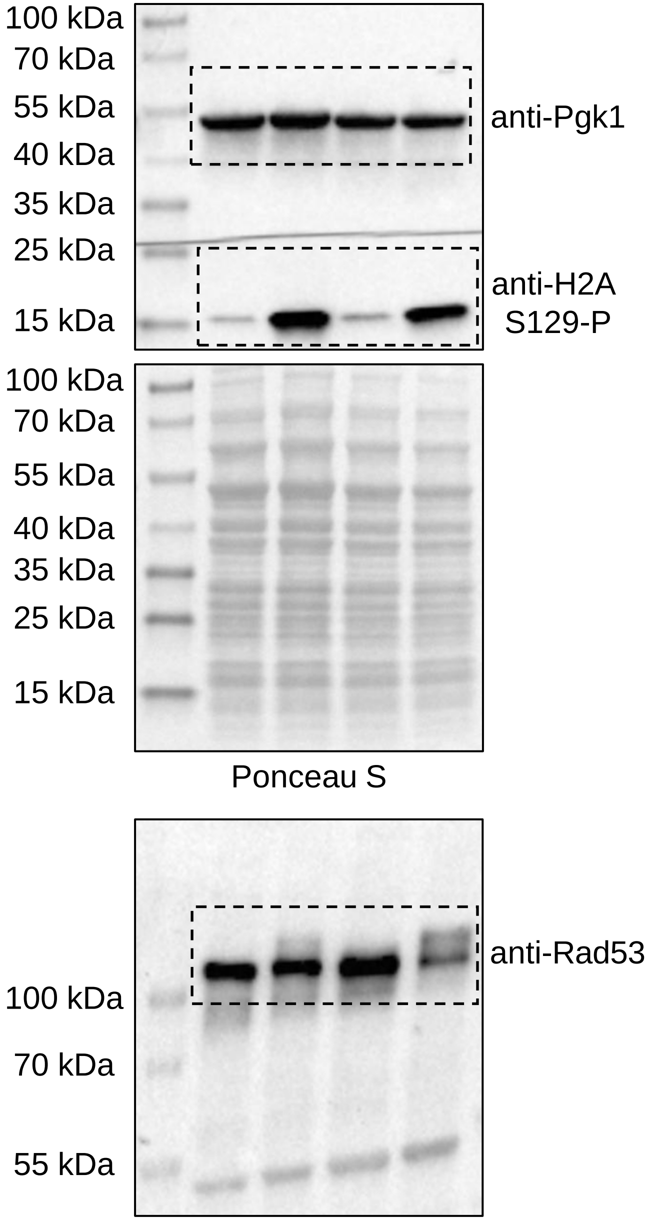


**Fig. S8B**

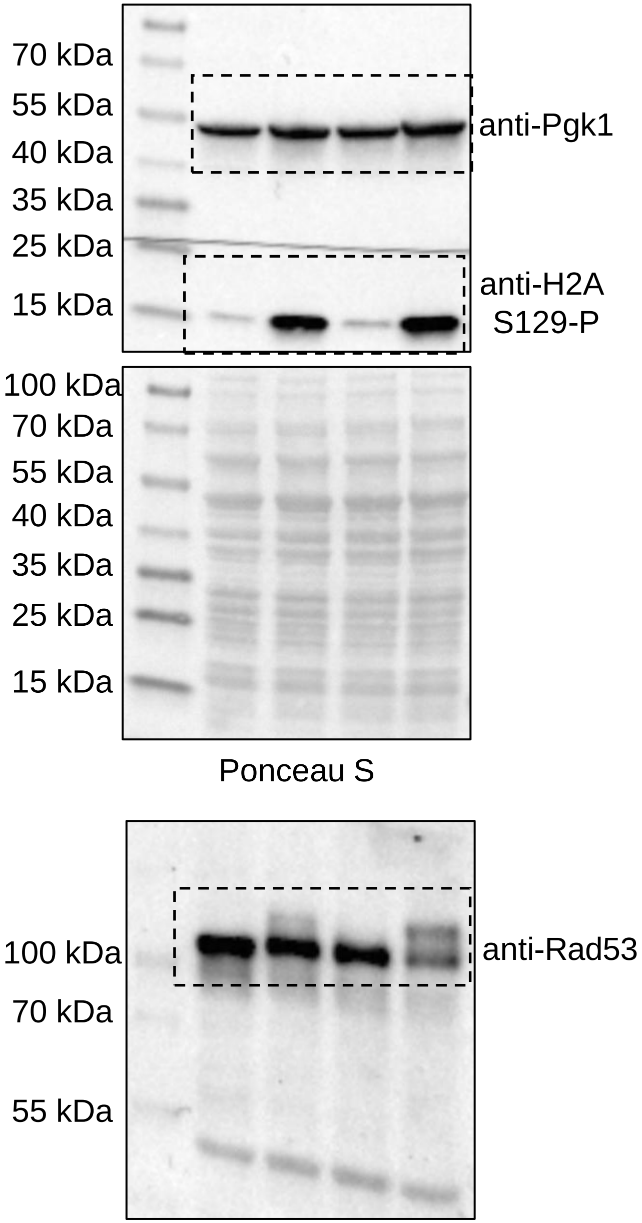

**Fig. S9B**

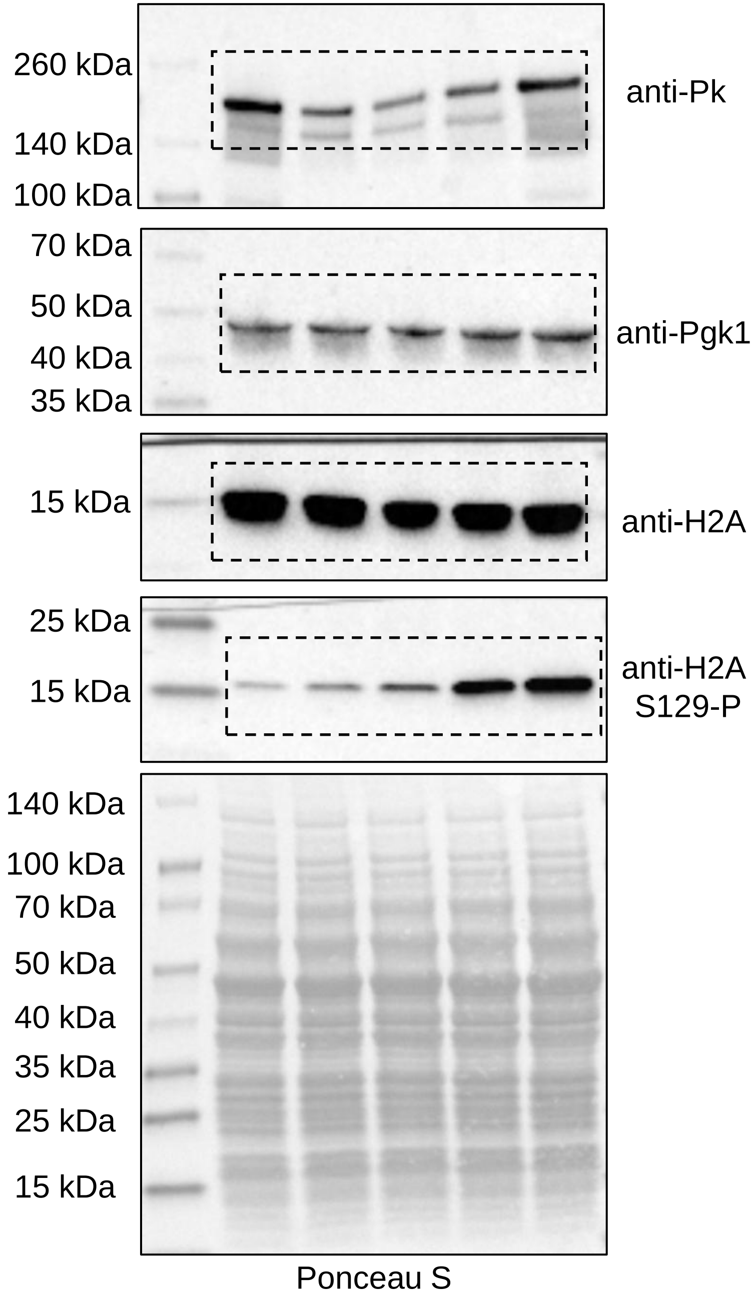
260 kDa
140 kDa
100 kDa
anti-Pk
70 kDa
50 kDa
40 kDa
35 kDa
anti-Pgk1
15 kDa
anti-H2A
25 kDa
15 kDa
anti-H2A
S129-P
140 kDa
100 kDa
70 kDa
50 kDa
40 kDa
35 kDa
25 kDa
15 kDa
Ponceau S

**Fig. S10A**

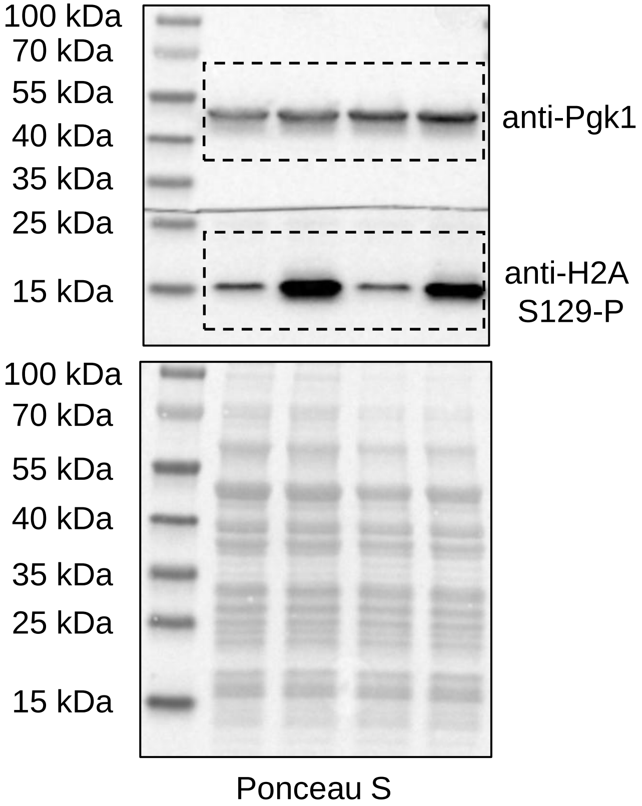


**Fig. S10B**

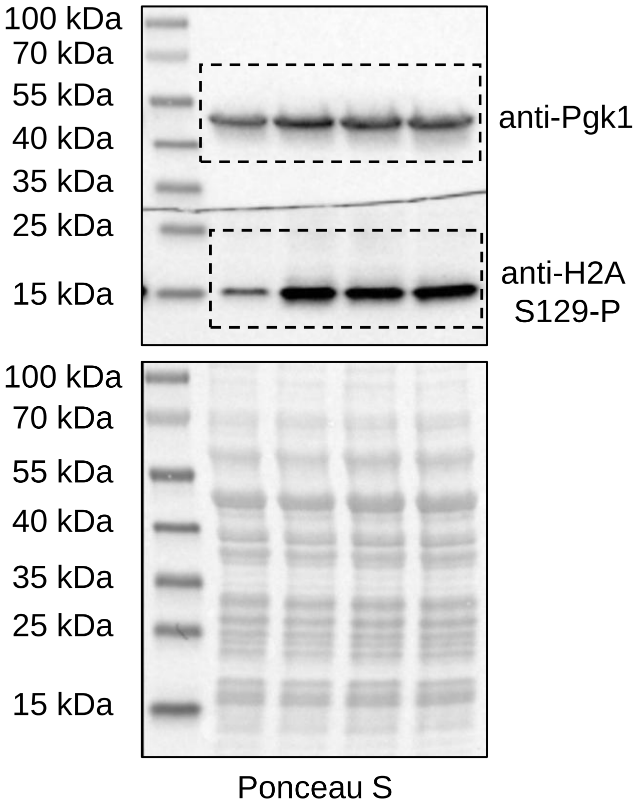

**Fig. S10C**

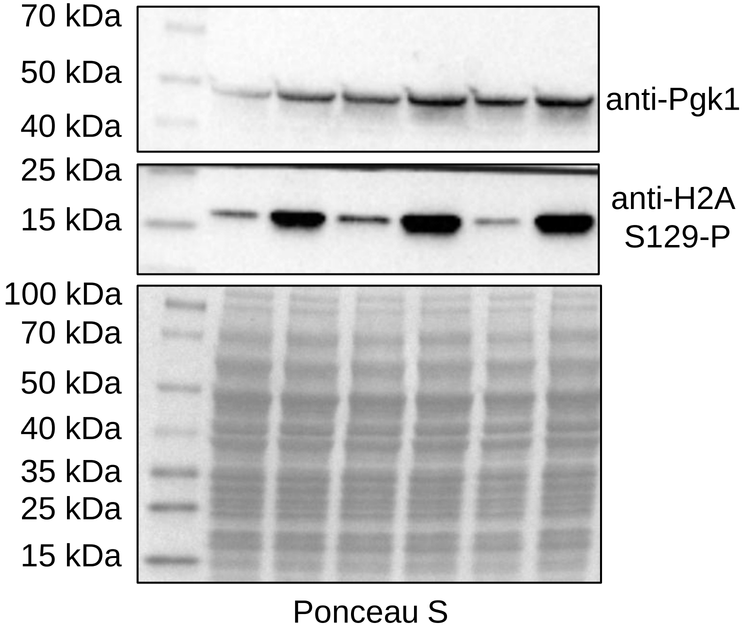


**Fig. S10D**

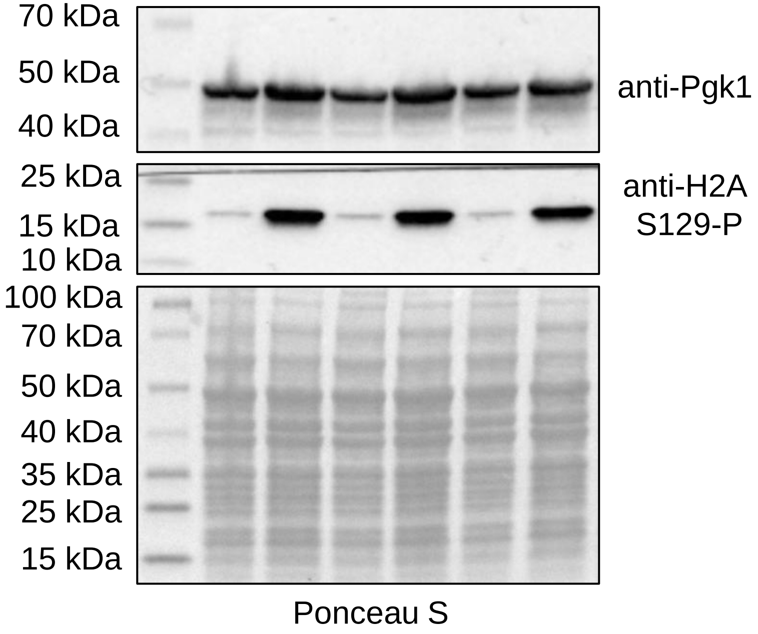

**Fig. S11C**

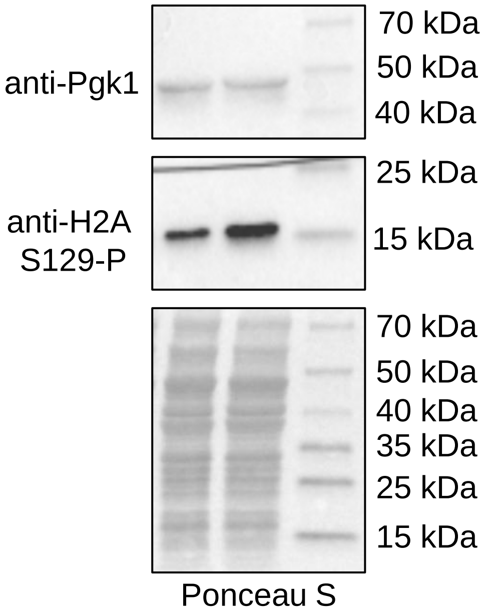


**Fig. S11D**

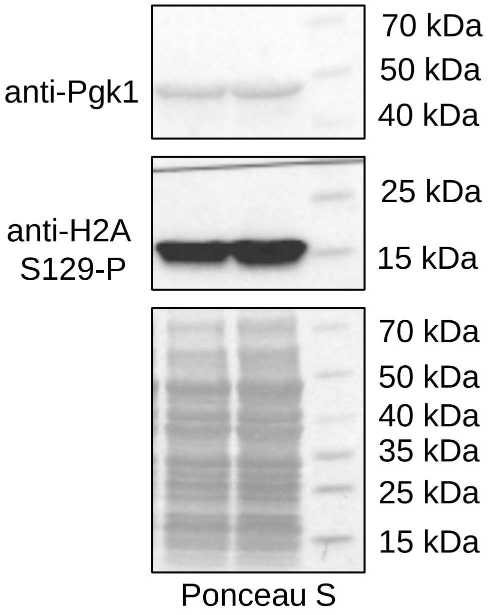

#### Fig. S11E

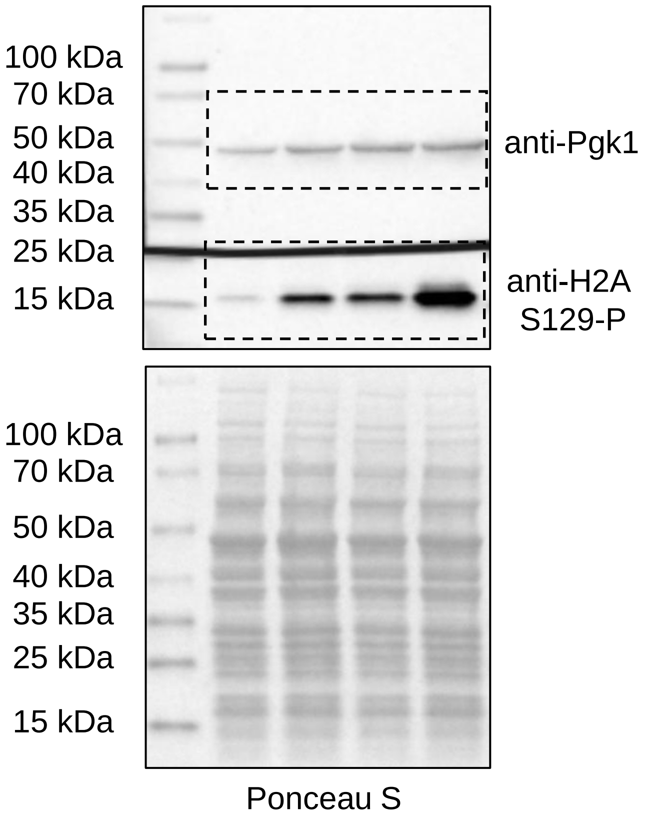


#### Fig. S11F

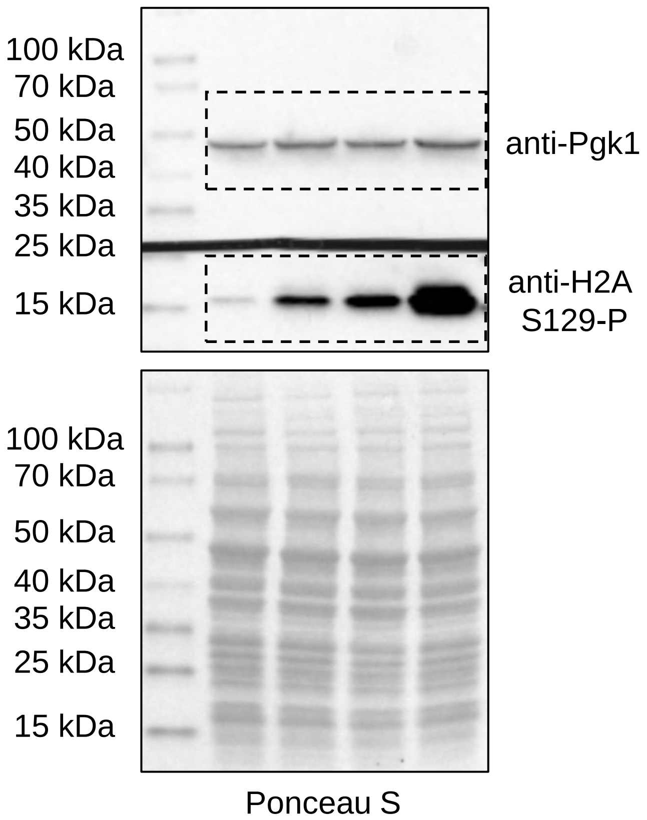


#### Fig. S11I

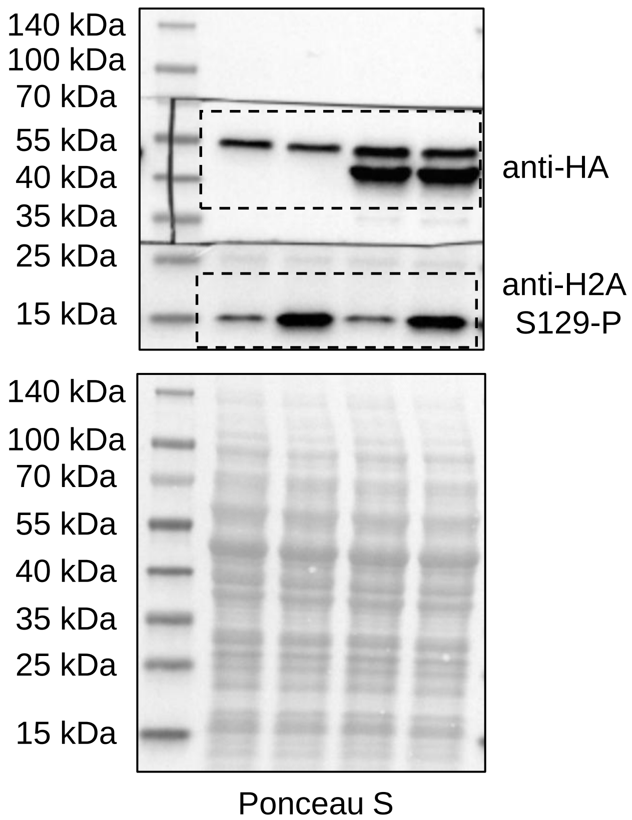

**Fig. S12A**

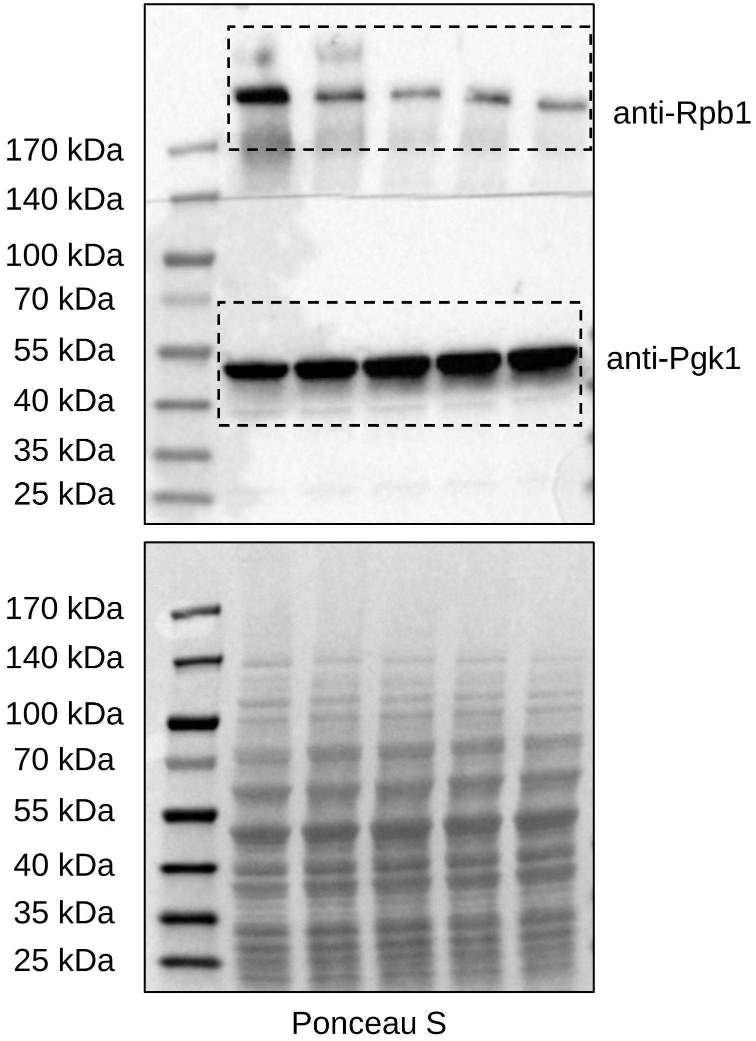


**Fig. S12B**

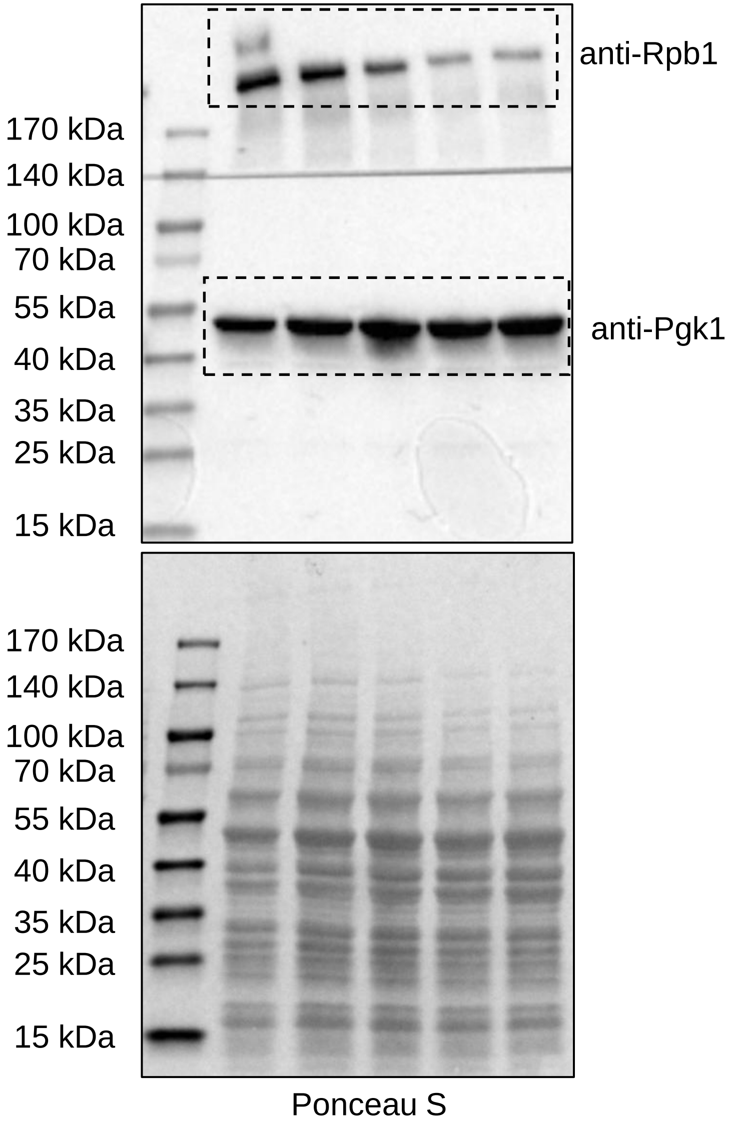


**Fig. S12C**

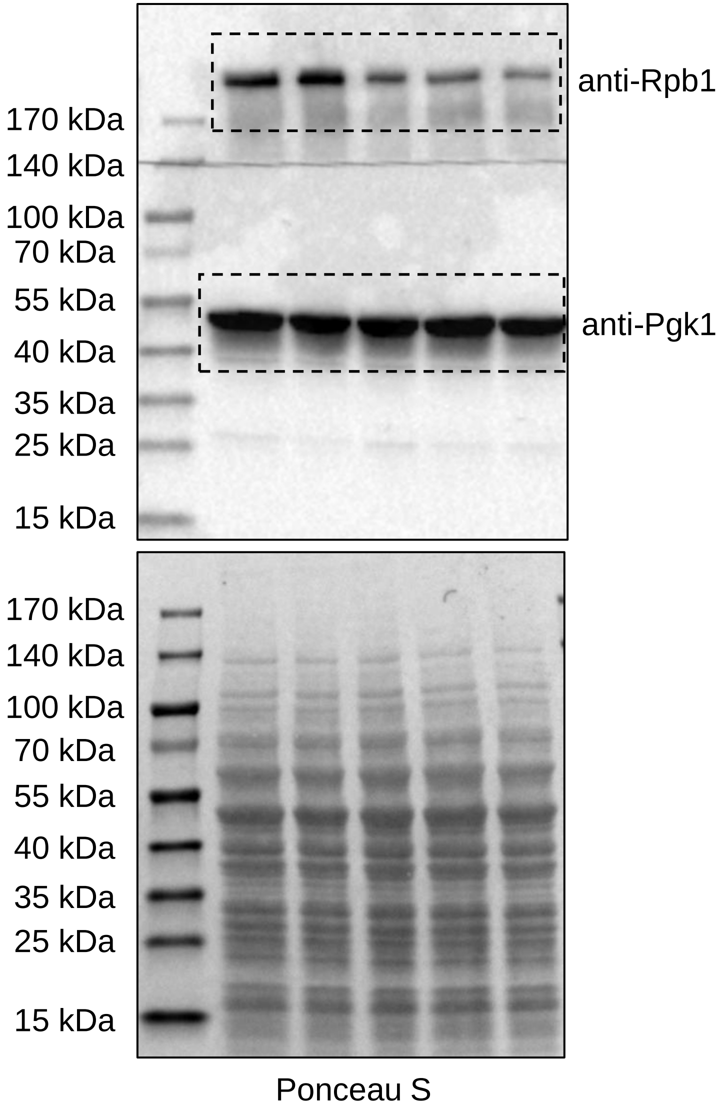

**Fig. S12D**

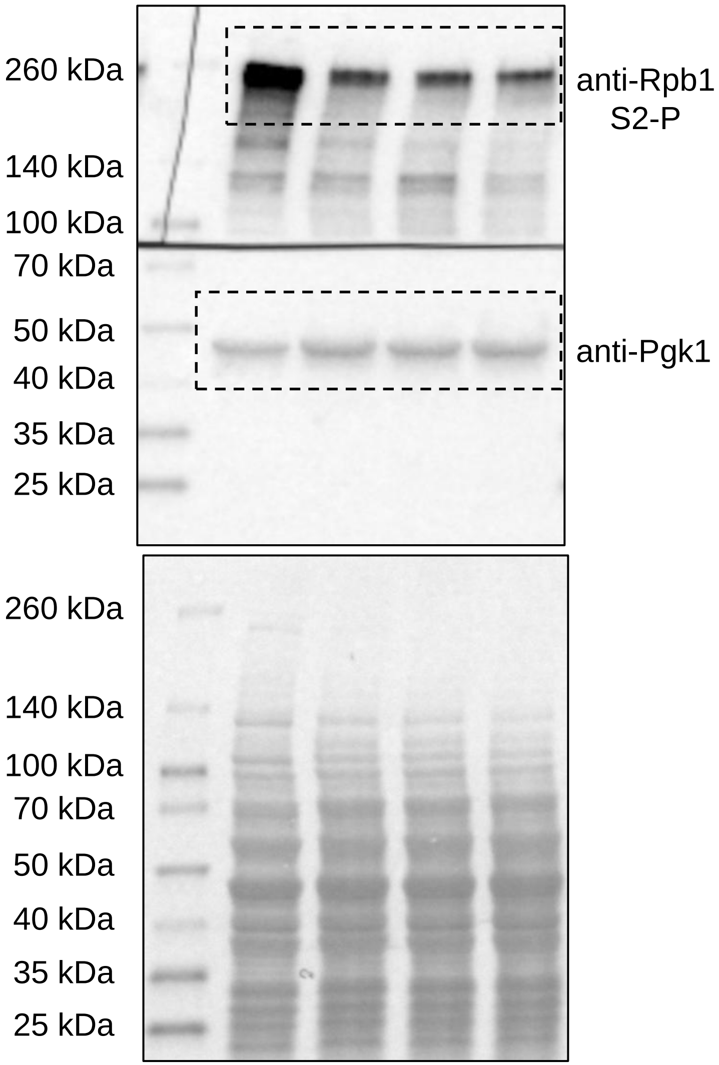


Ponceau S

**Fig. S12E**

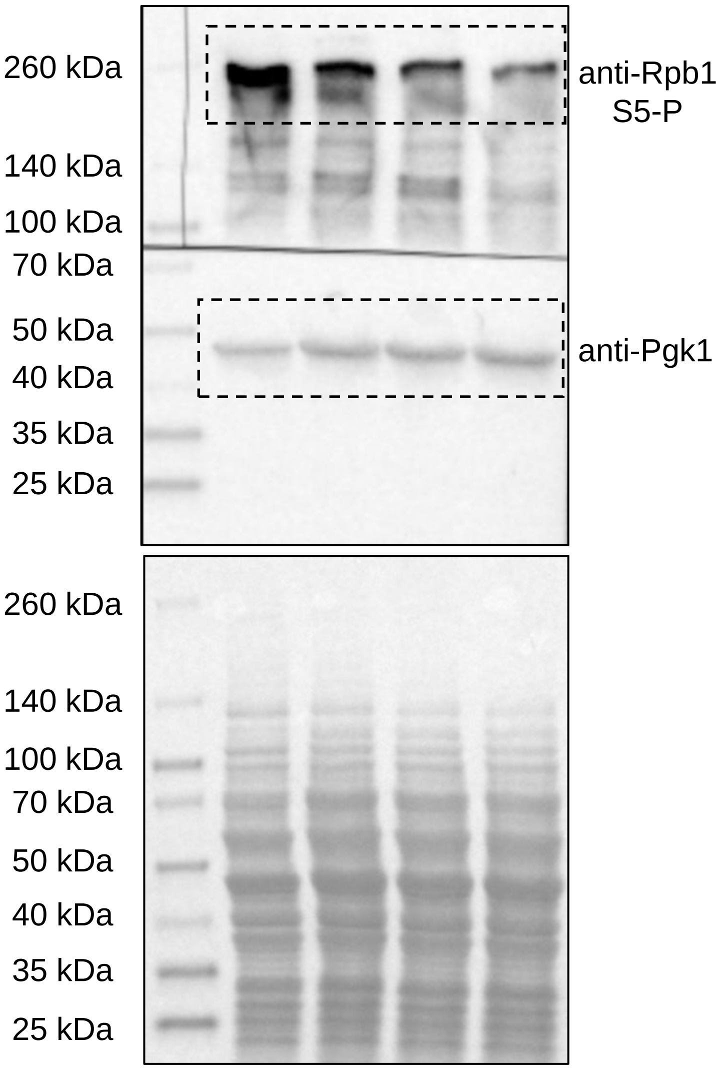


Ponceau S

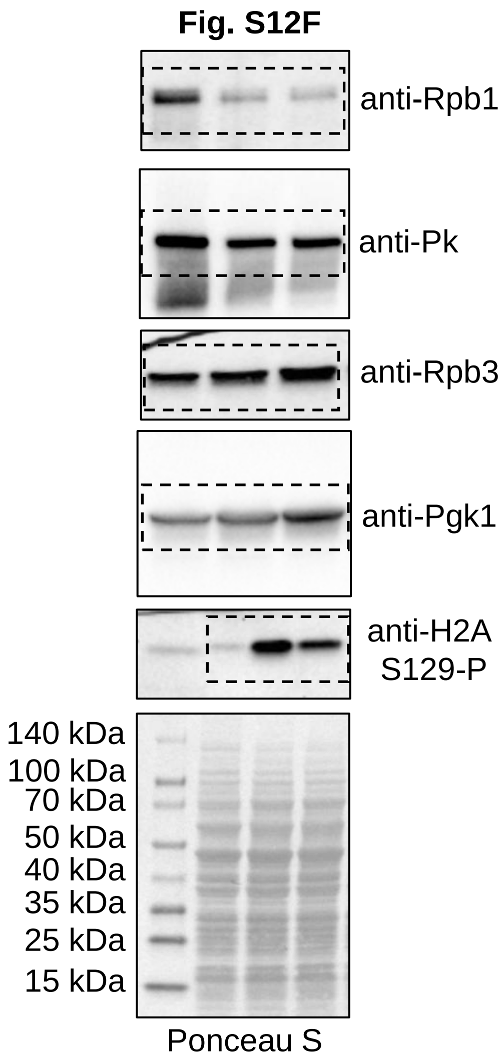
Fig. S12F
anti-Rpb1
anti-Pk
anti-Rpb3
anti-Pgk1
anti-H2A S129-P
140 kDa
100 kDa
70 kDa
50 kDa
40 kDa
35 kDa
25 kDa
15 kDa
Ponceau S

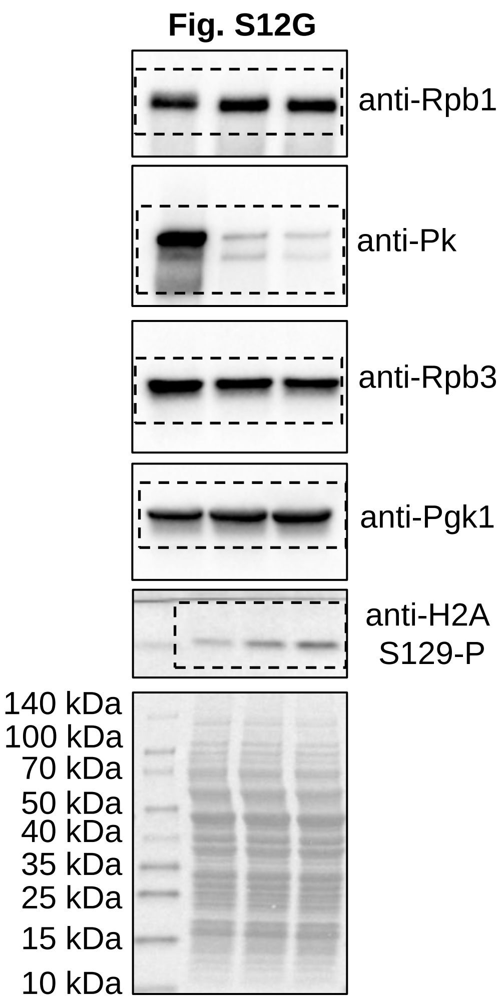
Fig. S12G
anti-Rpb1
anti-Pk
anti-Rpb3
anti-Pgk1
anti-H2A S129-P
140 kDa
100 kDa
70 kDa
50 kDa
40 kDa
35 kDa
25 kDa
15 kDa
10 kDa

**Fig. S13B**

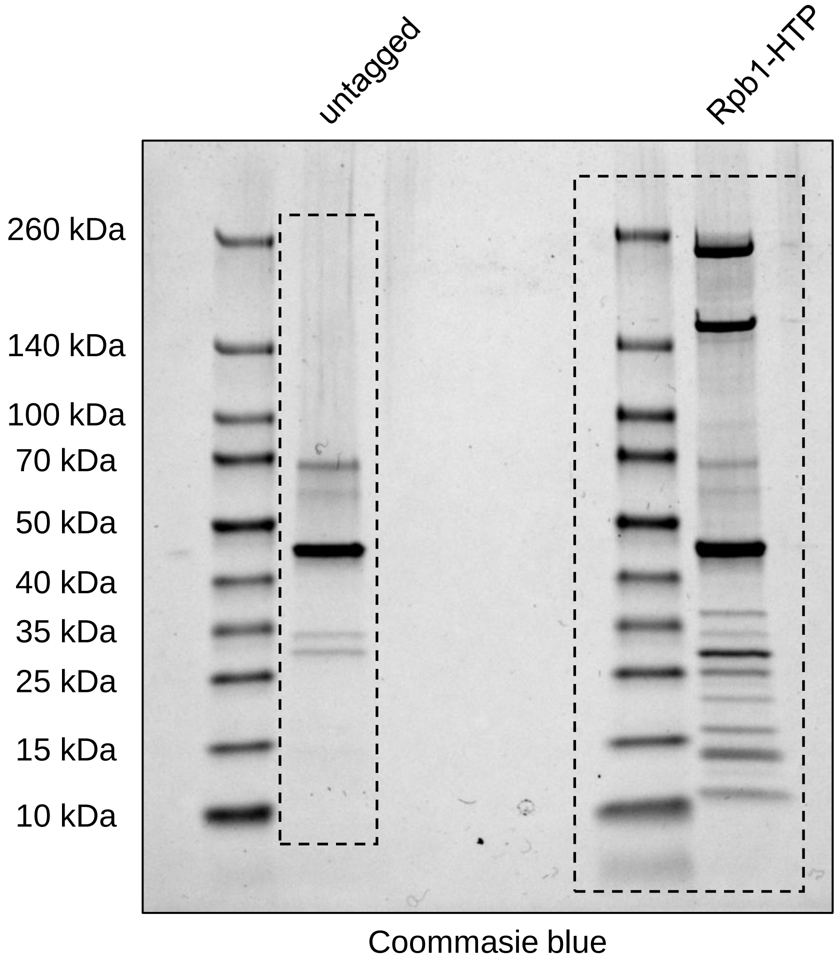
untagged
Rpb1-HTP
260 kDa
140 kDa
100 kDa
70 kDa
50 kDa
40 kDa
35 kDa
25 kDa
15 kDa
10 kDa
Coommasie blue
